# Cell type-specific prediction of 3D chromatin organization enables high-throughput *in silico* genetic screening

**DOI:** 10.1101/2022.03.05.483136

**Authors:** Jimin Tan, Nina Shenker-Tauris, Javier Rodriguez-Hernaez, Eric Wang, Theodore Sakellaropoulos, Francesco Boccalatte, Palaniraja Thandapani, Jane Skok, Iannis Aifantis, David Fenyö, Bo Xia, Aristotelis Tsirigos

## Abstract

The mammalian genome is spatially organized in the nucleus to enable cell type-specific gene expression. Investigating how chromatin organization determines this specificity remains a challenge. Methods for measuring the 3D chromatin organization, such as Hi-C, are costly and bear strong technical limitations, restricting their broad application particularly in high-throughput genetic perturbations. In this study, we present C.Origami, a deep neural network model that performs *de novo* prediction of cell type-specific chromatin organization. The C.Origami model enables *in silico* experiments to examine the impact of genetic perturbations on chromatin interactions in cancer genomes and beyond. In addition, we propose an *in silico* genetic screening framework that enables high-throughput identification of impactful genomic regions on 3D chromatin organization. We demonstrate that cell type-specific *in silico* genetic perturbation and screening, enabled by C.Origami, can be used to systematically discover novel chromatin regulatory mechanisms in both normal and disease-related biological systems.

## Introduction

In mammalian cells, interphase chromosomes are hierarchically organized into large compartments which consist of multiple topologically associating domains (TADs) at the sub-megabase scale^1^. Chromatin looping within TADs functions to restrict enhancer-promoter interactions at the kilobase scale for regulating gene expression^1–3^. The perturbation of TADs, such as disrupting TAD boundary, can lead to aberrant chromatin interactions and changes in gene expression^4–7^. As a result, mutations that disrupt 3D genome organization can substantially affect developmental programs and play important roles in genetic diseases and cancer^4,5,8,9^.

The higher-order organization of the genome is largely determined by intrinsic DNA sequence features known as *cis*-regulatory elements that are bound by *trans*-acting factors in a sequence specific manner^10^. For example, the location and orientation of CCCTC-binding factor (CTCF) binding sites act as a landmark for defining boundaries of TADs. Other factors, such as the cohesin complex proteins, act together to regulate chromatin interaction via loop extrusion^10,11^. While most TADs are conserved across cell types, a substantial amount (>10%) of TADs are dynamic and vary in different cells^12^. In addition, widespread cell type-specific chromatin-looping contributes to the precise regulation of gene expression^3,13^. These fine-scale chromatin interactions are controlled by chromatin remodeling proteins and transcription factors such as GATA1, YY1, and mediator proteins^2,14–16^. While the general organization of chromatin organization is largely well described, the current challenge is to reveal the principles underlying cell type-specific chromatin folding. Chromatin conformation capture technologies, such as Hi-C, are used for examining chromatin structure underlying gene regulation at fine-scales and across cell types^17,18^. However, these approaches are typically time- and resource-consuming, and require large cell numbers^18^. In addition, experimental tools are limited by the process of aligning sequencing reads to a specific reference genome, making it challenging for experiments involving *de novo* genome rearrangement. These limitations prohibit their wide-scale applications in investigating how chromatin organization determines cell type-specific gene expression, especially in gene regulation studies involving genetic perturbation and in rearranged chromosomes such as cancer genomes.

Owing to its ability to model complex interactions, deep learning has emerged as a powerful approach for studying genomic features. Leveraging *in silico* perturbations based on deep learning models could effectively reduce the resources required for *de novo* analyses of chromatin organization through experiments^19,20^. Since intrinsic features in DNA sequence of the genome partially determine its general folding principles, an approximate prediction of chromatin organization can be made using sequence alone^21–23^. However, due to the lack of specific genomic features which govern chromatin interactions^10^, approaches that rely solely on DNA sequence are unable to predict cell type-specific chromatin interactions^21–23^. Conversely, methods that rely only on chromatin profiles lack the consideration of DNA sequence features, thus generally requiring multiple epigenomic data to improve predictive power^24–29^. The limitations of current methods make them infeasible for *in silico* experiments studying how DNA sequence features and *trans*-acting factors work together to shape chromatin organization for accurate gene expression regulation.

We propose that an accurate *de novo* prediction of chromatin folding requires a model which effectively recognizes both DNA sequence and cell type-specific genomic features. Meanwhile, for the model to be practical, it should minimize the requirement for input information without performance loss. Based on these principles, we developed C.Origami, a deep neural network that synergistically integrates DNA sequence features and two essential cell type-specific genomic features: CTCF binding and chromatin accessibility signal. C.Origami achieved accurate *de novo* prediction of cell type-specific chromatin organization in both normal and rearranged genomes.

The high accuracy of C.Origami enables *in silico* genetic perturbation experiments that interrogate the impact on chromatin interactions, and moreover, allows systematic identification of cell type-specific regulation mechanisms of genomic folding through *in silico* genetic screening. Applying *in silico* genetic screening to T-cell acute lymphoblastic leukemia (T-ALL) cells and normal T cells, we identified a loss of insulation event at the upstream of *CHD4* in T-ALL, resulting in increased chromatin interaction between *CHD4* promoter and distal *cis*-elements. The high-throughput *in silico* genetic screening framework also makes it possible to identify a compendium of cell type-specific *trans*-regulators across multiple cell types. Additionally, we found that CDK7 plays a broader role in regulating 3D chromatin organization than that of NOTCH1, consistent with extensive experimental results by examining Hi-C contact matrices upon pharmacological inhibition of CDK7 and NOTCH1^30^. Together, our results demonstrate that the high performance of C.Origami enables systematic *in silico* genetic perturbation and screening experiments for identifying critical cell type-specific *cis*-elements and *trans*-acting regulators, thus empowering future studies of 3D chromatin regulation studies.

## RESULTS

### C.Origami: a multimodal architecture for predicting cell type-specific 3D chromatin organization

To achieve accurate and cell type-specific prediction of genomic features, we first developed Origami, a generic multimodal architecture, to integrate both nucleotide-level DNA sequence and cell type-specific genomic signal (Fig. 1a, excluding decoder). Specifically, the former enables recognition of informative sequence motifs, while the later provides cell type-specific features. Origami consists of two encoders, a transformer module, and a decoder (Fig. 1a, see Methods). The two encoders are 1D convolutional neural networks that condense DNA sequence and genomic features separately. The two streams of encoded features are then concatenated and further processed by a transformer module, which allows the encoded information to exchange between different genomic regions^31^. The decoder in Origami synthesizes the processed information to make predictions, and depending on the task, can be customized to specific downstream prediction targets. In this study, we deployed a 2D dilated convolutional network with broad receptive field as a decoder for predicting chromatin organization represented by Hi-C contact matrices (see Methods). We therefore named this chromatin organization predicting variant C.Origami.

**Figure 1:**
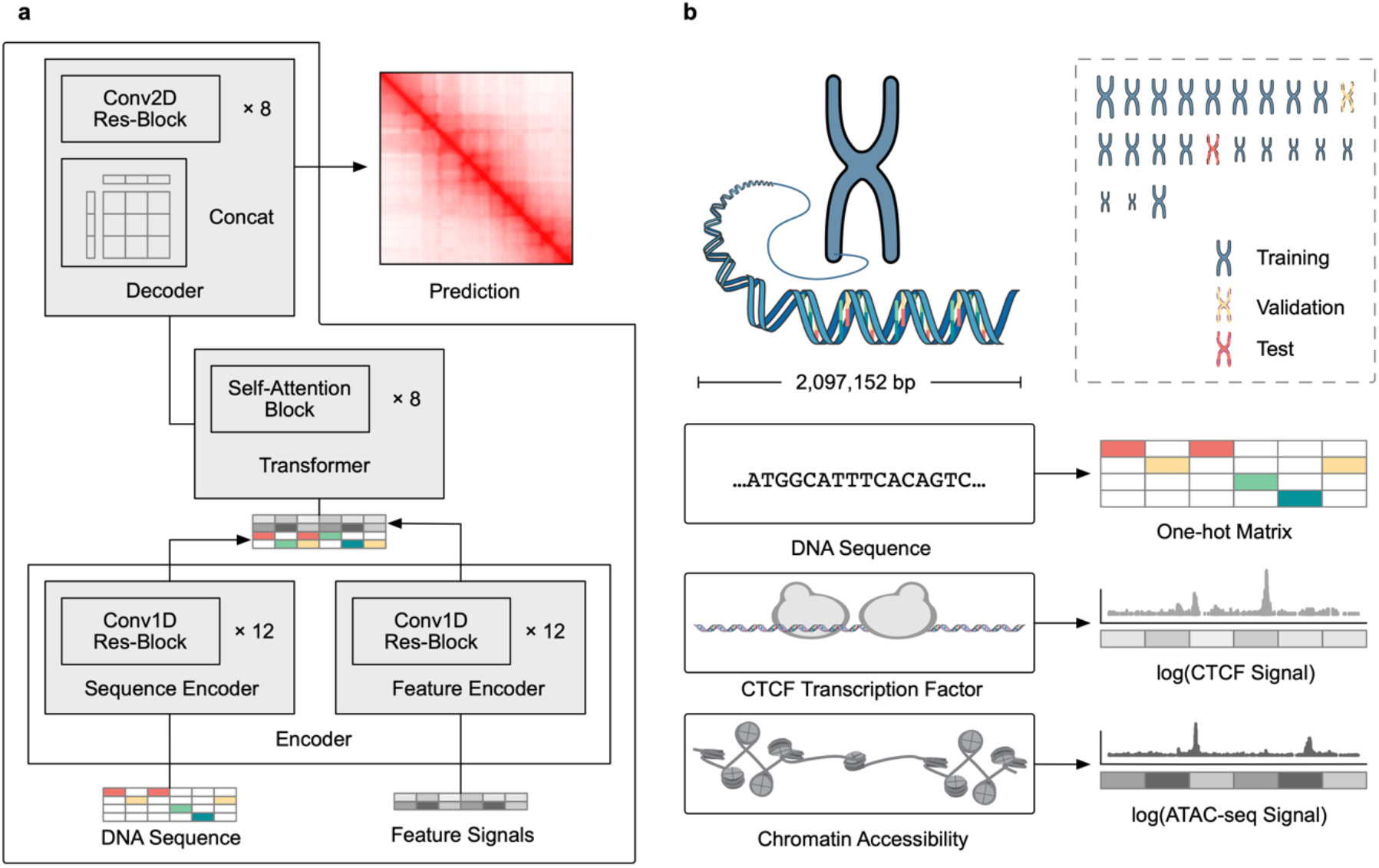
*De novo*, cell type-specific prediction of 3D chromatin organization with C.Origami. **a**, A schematic of C.Origami, a multimodal architecture for *de novo* prediction of chromatin organization. C.Origami adopts an encoder-decoder design, separately encoding DNA sequence features and cell type-specific genomic features. The two streams of encoded information are concatenated and processed by a transformer module. The decoder converts the processed 1D information to the final Hi-C interaction matrix. **b**, C.Origami predicts 3D chromatin organization by integrating DNA sequence, CTCF ChIP-seq signal and ATAC-seq signal as input features to predict Hi-C interaction matrix in 2 Mb windows.

C.Origami predicts chromatin organization within a 2 mega-base (2Mb) sized window to cover typical TADs in the genome while maximizing computation efficiency^1^. DNA sequence and genomic features within the window were separately encoded as nucleotide-level features (Fig. 1b, see Methods). The model reduces 2Mb wide genomic features down to 256 bins, and outputs a Hi-C contact matrix with a bin size of 8,192 bp. The target Hi-C matrix from the corresponding 2Mb genomic window was processed to have the same bin size. To train the model, we used data from IMR-90^32^, a fibroblast cell line isolated from normal lung tissue, and randomly split the chromosomes into training, validation (chromosome 10), and test set (chromosome 15) (Fig. 1b, top right).

When selecting genomic features as input for cell type-specific chromatin organization prediction, we considered three criteria: 1) representative for cell type specific chromatin organization; 2) widely available and experimentally robust; 3) minimized number of inputs to enable broad applicability of the model. CTCF binding is one of the most critical determinants of 3D genome organization, shaping the genome to organize into TADs^10^. Meanwhile, previous studies revealed widespread cell type-specific enhancer-promoter and promoter-promoter interactions which constitute a great portion of 3D chromatin organization at the accessible genomic regions^33–35^. In light of this knowledge, we envisioned C.Origami trained with CTCF ChIP-seq and ATAC-seq profiles, and together with nucleotide-level DNA sequence, would achieve high performance in predicting cell type-specific 3D chromatin organization (Fig. 1b).

To examine how different input features influence model performance, we first carried out an ablation study by training a set of prototype models with all seven combinations of the three input features, and then used validation loss to evaluate the model quality (Fig. 2a). We found that the model trained with DNA sequence alone has the highest validation loss – indicating lowest performance – due to its lack of cell type-specific genomic information. On the other hand, the model trained with a full set of input features – DNA sequence, CTCF ChIP-seq, and ATAC-seq profiles – consistently achieved the lowest validation loss. Moreover, replacing ATAC-seq profile with a key chromatin modification profile, H3K27ac, under-performs the original C.Origami model (Fig. 2a). Using only CTCF ChIP-seq or ATAC-seq profile as input give a mediocre performance. Notably, coupling genomic features with DNA sequence as training inputs always improves model performance (DNA + ATAC-seq > ATAC-seq; DNA + CTCF-binding > CTCF-binding; C.Origami > CTCF-binding + ATAC-seq), indicating that DNA sequence information contributes substantially for prediction quality.

**Figure 2:**
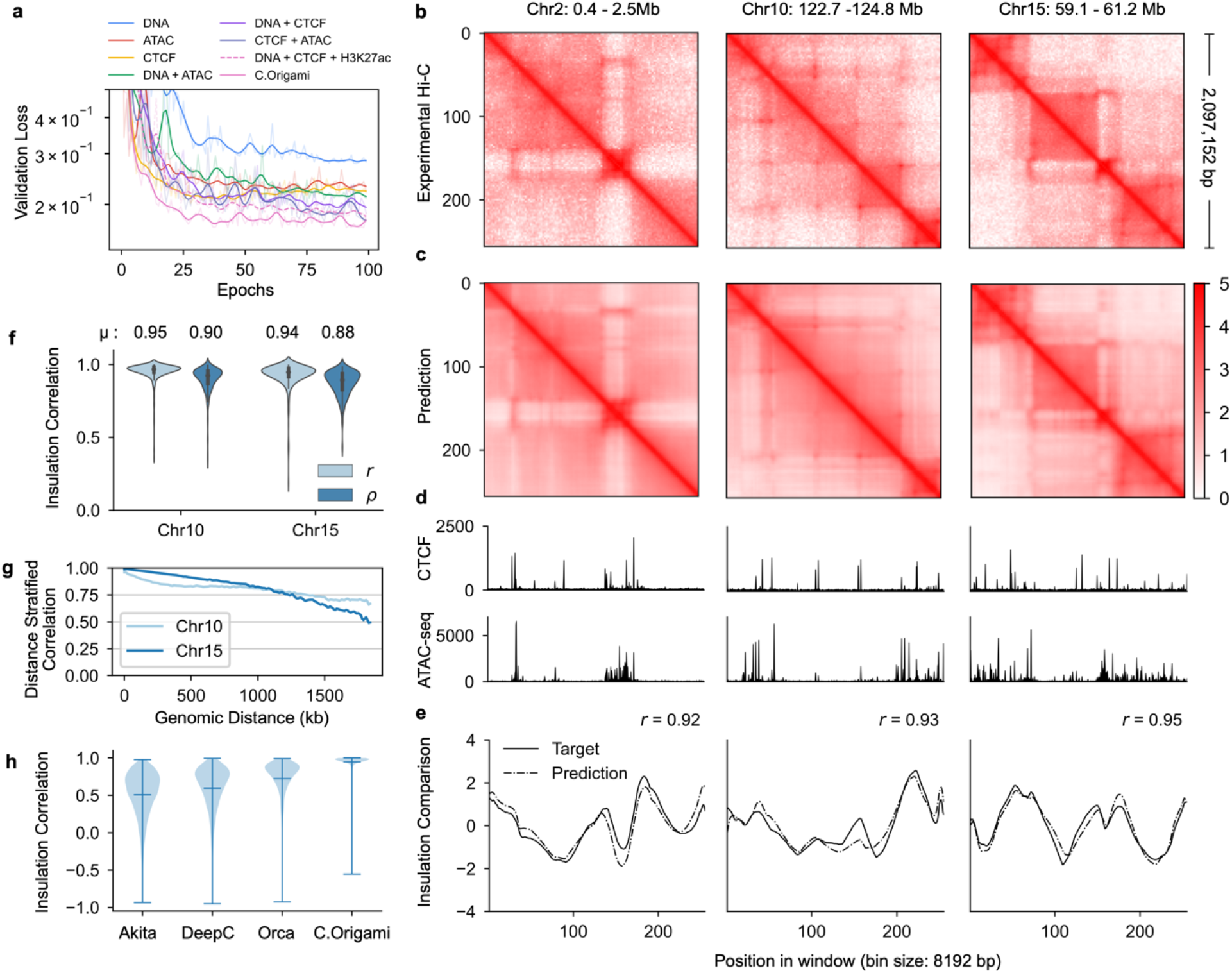
C.Origami accurately predicts 3D chromatin organization. **a**, Validation loss of prototype models trained from different combination of input features. **b-c**, Experimental Hi-C matrices (**b**) and C.Origami predicted Hi-C matrices (**c**) of IMR-90 cell line at chromosome 2 (left), chromosome 10 (middle), and chromosome 15 (right), representing training, validation and test chromosomes, respectively. **d**, Input CTCF binding and chromatin accessibility profiles. **e**, Insulation scores calculated from experimental Hi-C matrices (solid line) and C.Origami predicted Hi-C matrices (dotted line). Pearson correlation coefficients between prediction and target insulation scores is presented. **f**, Insulation score correlation between predicted and experimental Hi-C matrices across all windows in both validation and test chromosomes. Each group included both Pearson correlation (*r*) and Spearman correlation (*ρ*) coefficients. **g**, Chromosome-wide distance-stratified interaction correlation (Pearson) between prediction and experiment. **h**, Comparison of model performance across Akita, DeepC, Orca, and C.Origami using genome-wide insulation score correlation between prediction and experimental data from IMR-90 cells. Error bars in the violin plots indicate minimum, mean and maximum values within each group.

To further inspect the performance difference between C.Origami and models trained with incomplete inputs, we compared C.Origami with the model trained with DNA sequence and CTCF ChIP-seq signal. While the later model performed well in capturing the TAD structures and some chromatin loops, the model did not predict many fine-scale chromatin interaction features, especially in *de novo* prediction on a new cell type (Supplementary Fig. 2). These results indicate that integrating DNA sequence with CTCF binding signal alone is not sufficient for optimal prediction of cell type-specific 3D chromatin organization.

C.Origami trained with complete inputs achieved high-quality predictions for chromatin organization (Fig. 2, Supplementary Fig. 3). C.Origami predicted highly accurate contact matrices that emphasized both large topological domains and fine-scale chromatin looping events in samples from training, validation and test chromosomes (Fig. 2b-e and Supplementary Fig.3). Similar to the ablation study, compromising each of the input signals by random shuffling led to inferior performance, underscoring the necessity of including all input features for high-quality predictions (Supplementary Fig. 4). Last, we found that while it is possible to train the model using sparse input genomic features (ChIP-seq/ATAC-seq peaks) without significant performance penalty, the current C.Origami model trained with dense features (including peak profiles and sequencing background signals of ChIP-seq/ATAC-seq) achieved better performance, indicating that the model leveraged the nuanced genomic features to improve its prediction (Supplementary Fig. 5).

### Genome-wide evaluation of model performance

To systematically assess C.Origami, we calculated the insulation scores on validation and test chromosomes (see Methods). C.Origami achieved on average 0.95 and 0.94 insulation score correlation respectively (Fig. 2f). By plotting the insulation score correlation between prediction and experiment against Hi-C data intensity across the genome by chromosomes, we found that the prediction maintained uniform high performance, demonstrating the robustness of the model (Supplementary Fig. 6).

To evaluate the consistency of predicted Hi-C matrices, we calculated distance-stratified average intensity of Hi-C matrices from C.Origami prediction and experiment and found the same exponential decay pattern (Supplementary Fig. 7a). In addition, predicted chromatin structure from C.Origami were stable across neighboring regions. Therefore, consecutive predictions can be used to construct chromosome-wide prediction of Hi-C contact matrix by joining predictions across sliding windows (Supplementary Fig. 7b-d). Such genome-wide construction of Hi-C contact matrices allowed us to plot a distance-stratified correlation (Pearson) between the merged chromosome-wide prediction and experimental Hi-C (see Methods). C.Origami achieved correlation above 0.8 within 1Mb region and 0.6 within 1.5Mb (Fig. 2g, Supplementary Fig. 8).

Loop calling is a common analysis for identifying point-to-point interactions from Hi-C. As a third metric to evaluate C.Origami’s performance, we performed loop calling using global background as reference to capture significant chromatin interactions on both prediction and experimental Hi-C in IMR-90 cells (see Methods). We found that C.Origami achieved good performance in loop detection, with an AUROC of 0.92 for the top 5000 predicted loops (Supplementary Fig. 9). We further categorized loops by the chromatin background of loop anchors, resulting in three major categories: CTCF-CTCF loop, promoter-enhancer loop, and promoter-promoter loop (Supplementary Figure 9a-b). We found that C.Origami-predicted Hi-C maps can further predict chromatin loops comparable to the experimental results under each loop category (Supplementary Figure 10).

Last, we compared C.Origami against three recent sequence-based approaches, Akita^22^, DeepC^23^, and Orca^36^. Since the four models were trained with different scaling, resolution and prediction target customization, we included in the benchmark a set of preprocessing and normalization steps to standardize the results (See Methods, and Supplementary Fig. 11-13). To evaluate the performance of the four models, we compared their predicted results to experimental data by calculating: 1) insulation score correlation, 2) observed/expected Hi-C map correlation, 3) mean squared error (MSE), and 4) distance-stratified correlation using results from IMR-90 cells (see Methods). We found that C.Origami outperforms previous methods in all four comparison matrices (Fig. 2h, Supplementary Fig. 14).

### *De novo* prediction of cell type-specific chromatin organization

*De novo* prediction of cell type-specific 3D chromatin organization provides a valuable approach for studying genome regulation in new cell types. To assess C.Origami’s performance in *de novo* prediction of chromatin organization beyond the training cell type IMR-90, we applied the model to GM12878 cells using its corresponding CTCF ChIP-seq and ATAC-seq profiles. GM12878 is a lymphoblastoid cell line that differs substantially from IMR-90 in its chromatin organization^32^, as exemplified by locus Chr2:400,000-2,497,152 (Fig. 3a). Specifically, we highlighted a cell type-specific interaction related to chromatin accessibility changes (black arrowhead) and a distal interaction that associates with both CTCF and ATAC-seq signal changes (gray arrowhead, Fig. 3c). These cell type-specific features were demonstrated by differences in their signal intensity in Hi-C and genomic tracks (Fig. 3a and 3c, right).

**Figure 3:**
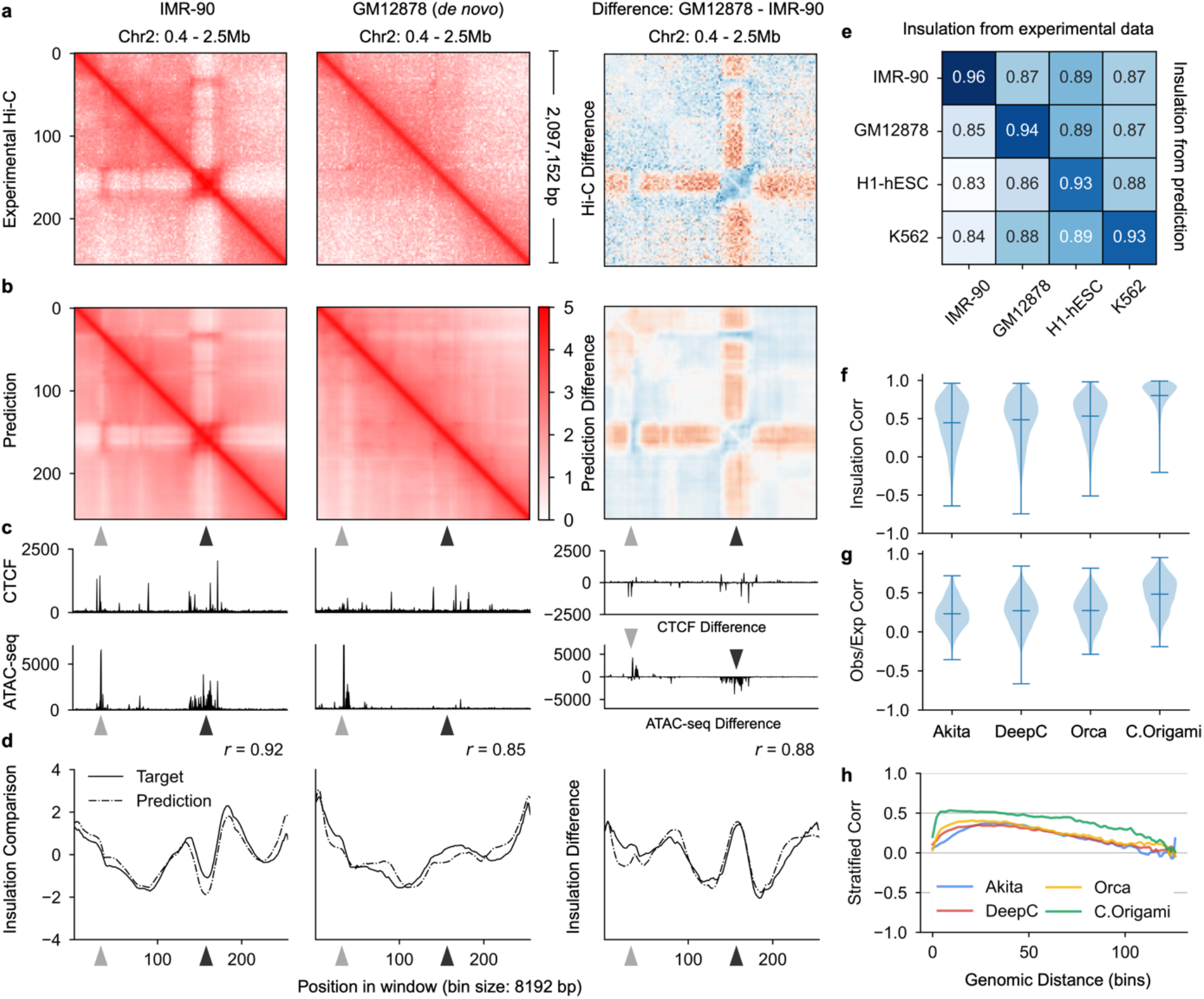
Cell type-specific *de novo* prediction of chromatin structure. **A**, Experimental Hi-C matrices from IMR-90 (left) and GM12878 (middle) cell lines at chromosome 2, and their differences (right). **B**, C.Origami-predicted Hi-C matrices of IMR-90 (left) and GM12878 (middle), precisely recapitulated the experimental Hi-C matrices (**a**). The arrow heads highlighted differential chromatin interactions between the two cell types. **C**, CTCF binding profiles and chromatin accessibility profiles of IMR-90 (left), GM12878 (middle) and their difference (right). **D**, Insulation scores calculated from experimental Hi-C matrices (solid line) and C.Origami predicted Hi-C matrices (dotted line) of IMR-90 (left), GM12878 (middle) and their difference (right). **E**, Pearson correlation between insulation scores calculated from predicted and experimental Hi-C matrices across cell types. **F-h**, Genome-wide evaluation of sequence-based models and C.Origami using *de novo* prediction results from GM12878 cells. Presented metrics include insulation score correlation (**f)**, observed vs expected matrix correlation (**g)**, and distance-stratified correlation (**h**). Error bars in violin plots of **f** and **g** indicate minimum, mean and maximum values within each group.

To demonstrate the capability of C.Origami in cell type-specific *de novo* prediction, we predicted Hi-C matrices in both IMR-90 and GM12878 cells at the same locus. Notably, C.Origami was trained on IMR-90 and was never exposed to GM12878-specific inputs and Hi-C data. Therefore, C.Origami needs to transfer its knowledge to the new cell type. We found that C.Origami accurately captured the cell type-specific chromatin interaction features in GM12878 *de novo* prediction (Fig. 3a-c, left and middle). The difference between IMR-90 and GM12878 experimental Hi-C matrices was also reflected between IMR-90 and GM12878 predictions (Fig. 3a-c, right). The calculated insulation scores from the predicted Hi-C matrix were also highly correlated with the scores of the experimental data from both cell types (Fig. 3d, left and middle). In addition, the difference between insulation scores of the two cell types were highly correlated, showing that C.Origami captured the chromatin architectural difference between two cell types (Fig. 3d, right). We further expanded the *de novo* chromatin organization prediction to two more cell lines, embryonic H1-hESC and erythroleukemia K562. Again, our model achieved accurate predictions of cell type-specific chromatin organization with high specificity, demonstrating the robustness of C.Origami in *de novo* prediction and its practical potential for a broader application (Supplementary Fig. 15).

To systematically evaluate the performance of C.Origami in *de novo* prediction, we next carried out an analysis of genome-wide predictions. Although we presented multiple loci that have cell type-specific chromatin structures, many TAD boundaries are conserved across cell types^12^. To test the model on structurally different regions, we first identified a subset of genomic loci with differential chromatin structures between IMR-90 and GM12878 experimental Hi-C matrices. Regions with normal intensity (> 10% intensity quantile) and low similarity (< 20% insulation difference) between the experimental Hi-C matrices of the two cell types were selected. In total, ∼15% of the entire genome (∼450Mb) were included for evaluating the performance of cell type-specific Hi-C prediction (Supplementary Fig. 16a).

We calculated the correlation coefficient between the insulation scores of the predicted and experimental Hi-C matrices across all four cell types in structurally different genomic regions (Fig. 3e, Supplementary Fig. 16). In line with observations from the single-locus results (Fig. 3a-d), we found that predictions using input features from one cell type have the highest correlation coefficients with the experimental Hi-C data of the same cell type (Fig. 3e, scores at the diagonal line). The correlation coefficients between mismatched prediction and experimental data were lower, consistent with the expectation that the model predicts cell type-specific chromatin interactions (Fig. 3e, off-diagonal scores). Similarly, these results were recapitulated by correlation analysis using pixel-level observed/expected contact matrices (Supplementary Fig. 16c-d). As a control, we performed a similar analysis using structurally conserved genomic regions, characterized by normal intensity (> 10% intensity quantile) and high similarity (> 20% insulation difference) between IMR-90 and GM12878 (Supplementary Fig. 16d). As expected, we found the prediction in these regions was highly correlated with the experimental data across all cell types (Supplementary Fig. 16e-f). We further compared the insulation score of IMR-90 to that of the three other cell lines and found such insulation score difference calculated from prediction and experimental data were highly correlated (Supplementary Fig. 16g).

As an orthogonal validation, we performed loop calling on IMR-90 and GM12878 prediction and experimental Hi-C to evaluate C.Origami’s ability to detect cell type-specific chromatin loops. We found that C.Origami can predict significant (log2fc > 1) IMR-90-specific and GM12878-specific loops with 0.88 and 0.87 AUROC, respectively (Supplementary Fig. 17). Cell type-specific loops under different categories also achieved similar performance (Supplementary Fig. 18).

Since DNA sequence-based models are unable to generalize to unseen cell types, we expect C.Origami to have an advantage in cell type-specific *de novo* prediction. This performance gap can be observed by comparing *de novo* predictions generated by sequence-based models and C.Origami in GM12878 cells (Supplementary Figure 19). Comparing genome-wide cell type-specific predictions in regions with cell type-specific chromatin organizations (see Methods), we again found that C.Origami outperformed sequence-based models by a large margin under all metrics, with higher insulation score correlation, higher observed/expected Hi-C matrix correlation, lower mean squared error (MSE), and higher distance-stratified correlation (Fig. 3f-h, Supplementary Figure 20).

The mouse genome differs from human in its genomic components but the two share similar mechanisms in 3D chromatin organization^1,34,37^. We sought to test whether C.Origami could perform *de novo* prediction across species. We found that C.Origami trained with human IMR-90 genomic features predicted mouse chromatin organization with good quality (Supplementary Figure 21). The overall performance in mouse was lower compared to that in human, possibly due to species-specific genomic features that were learned by the model during training. Notwithstanding its good performance, the accuracy of C.Origami could be further improved by training a model on mouse data to adapt to mouse sequence and genomic features. Together, these results indicate that C.Origami can extract and transfer the conserved genome organization principles learned across species.

Last, we tested whether C.Origami could predict the chromatin-organization changes upon removal of key *trans-*acting regulators, such as CTCF. Previous study found that acute degradation of CTCF protein led to the disappearance of TADs in mouse embryonic stem cells, and subsequent restoration of CTCF reestablished TAD structures^38^. We simulated such experiments by predicting chromatin organizations in pre-CTCF-depletion, CTCF-depleted, and CTCF-restored conditions (see Methods). We found that C.Origami accurately predicted the TAD-loss and restoration changes upon CTCF depletion and restoration, respectively (Supplementary Fig. 22).

### Accurate prediction of C.Origami enables cell type-specific *in silico* genetic experiments

Chromosomal translocations and other structural variants generate novel recombinant DNA sequences, subsequently inducing new chromatin interactions which may be critical in tumorigenesis and progression^8,40^. However, the allelic effect and high heterogeneity of translocation and structural variations frequently seen in cancer genomes make it challenging to study their custom genome organizations. As an example, CUTLL1, a T-cell acute lymphoblastic leukemia (T-ALL) cell line, incorporated a heterozygous t(7;9) translocation, a recombination of chromosome 7 and chromosome 9 (Fig. 4a)^39^. The translocation introduces new CTCF binding signals from chromosome 9 to chromosome 7, leading to the formation of a neo-TAD structure which can be observed in experimental Hi-C (Fig. 4b, see Methods)^30^.

**Figure 4:**
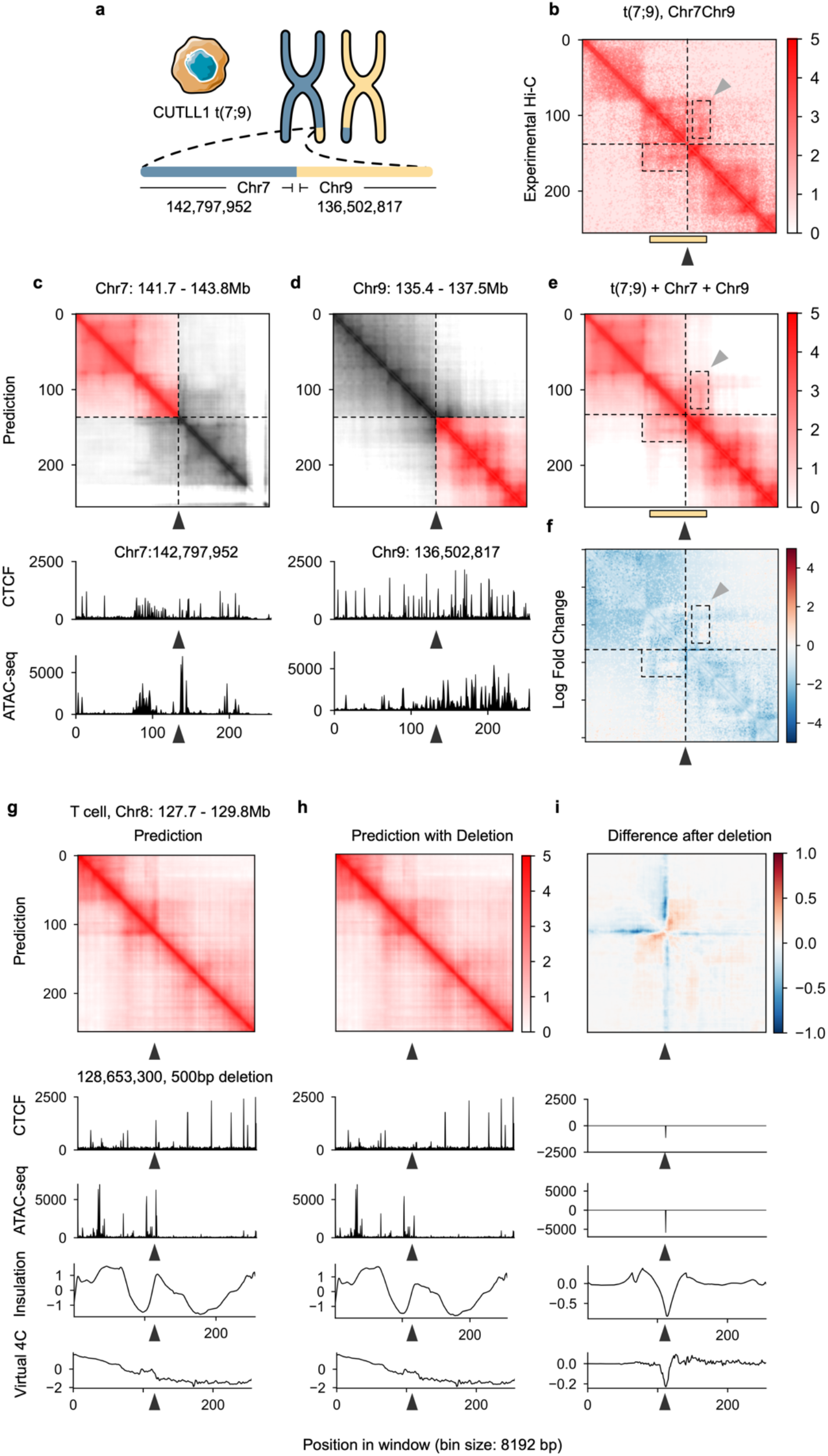
C.Origami enables prediction of 3D chromatin organization upon *in silico* genetic perturbations. **a**, Chromosomal translocation between chromosome 7 and chromosome 9 in CUTLL1 T cell leukemia cells^39^. **b**, Experimental Hi-C data mapped to a custom reference chromosome with t(7;9) translocation^30^. **c-d**, C.Origami prediction of chromatin organization of chromosome 7 (**c**) and chromosome 9 (**d**) in CUTLL1 cells. The windows represented intact chromosomal loci centered at the translocation sites in CUTLL1 cells. **e**, A simulated Hi-C contact matrix using prediction for mimicking of experimental mapping results. The simulated result was averaged from the prediction of both normal and translocated alleles, indicating heterozygous translocation. The yellow bar highlights the neo-TAD at the translocation locus. Black and gray arrowhead indicates the translocation site and a stripe in the neo-TAD, respectively. The predicted Hi-C matrix was aligned to the experimental Hi-C matrix in **d**. **f**, Log fold change between experiment and predicted Hi-C matrix at the t(7;9) translocation site in CUTLL1 cells. **g-i**, A 500bp deletion in chromosome 8 led to chromatin looping changes in T cells. The presented 2Mb window starts at the promoter region of *MYC*, and the experimental deletion perturbed a CTCF binding site at the arrowhead location^30^. The presented results include C.Origami prediction of the Hi-C contact matrices with (**g**) or without (**h**) the deletion, and their difference (**i**). Virtual 4C signals, calculated from the predicted Hi-C matrices, are shown at the bottom.

We highlight that C.Origami provides a high-performance alternative for discovering new chromatin interactions at rearranged genomic loci. To examine the performance of C.Origami in predicting chromatin organization from rearranged cancer genomes, we predicted Hi-C contact matrices from both the normal and translocated alleles, and then averaged the two matrices to mimic the allele-agnostic Hi-C mapping in the experimental data (Fig. 4c-e, see Methods). We found that the Hi-C map generated by C.Origami accurately predicted the neo-TAD structure covering the t(7;9) translocation site (Fig. 4e-f). Specifically, we found a stripe extending from translocated chromosome 9 to chromosome 7, indicating a novel regulation within the neo-TAD (Fig. 4b and 4e, dotted box and gray arrowhead). We additionally performed the same *in silico* experiments at three verified translocation loci in K562 cells and obtained similar results^41^ (Supplementary Fig. 23). The accuracy in detecting novel chromatin interaction at chromosomal translocation sites demonstrated C.Origami’s high performance and potential in future cancer genomics studies.

Moreover, we expect the high performance of C.Origami to enable cell type-specific *in silico* genetic perturbation experiments as a fast and cost-efficient approach for studying chromatin interaction mechanisms. As an example, while CTCF binding site has been found critical for organizing TADs via experimental perturbations^4–6^, not all perturbations at the CTCF binding sites led to the similar TAD changes due to motif redundancy and the complicated roles of CTCF in chromatin regulation^42–44^. Notably, experimental perturbation requires sophisticated genetic deletion followed by assessment through chromatin conformation capture techniques. Instead of experimentally performing such genetic studies, we modeled deletions of CTCF-binding at the TAD boundary sequences *in silico*, and subsequently predicted local chromatin interaction maps with C.Origami. We found that *in silico* deletion at TAD boundaries with CTCF-binding led to TAD merging events between the originally insulated TADs with a sharp drop in insulation score at the perturbed boundaries (Supplementary Fig. 24).

To further investigate the validity of *in silico* genetic perturbation, we applied C.Origami to predict chromatin interactions at loci with known experimental validations. Our previous study showed that disrupting a CTCF-binding site near *MYC* locus reduced the chromatin looping efficiency in human naive CD4+ T cells, resulting in a reduced chromatin insulation^30^. Applying C.Origami at the locus without perturbation, we found a stripe in the predicted Hi-C matrix (Fig. 4g, arrowhead). A 500bp *in silico* removal of the CTCF-binding region attenuated the stripe (Fig. 4h-i). Based on the two predicted Hi-C matrices, we calculated virtual 4C difference before and after perturbing the CTCF binding site and found them to be consistent with previous experimental data (Supplementary Fig. 7E in Kloetgen, *et al*)^30^. Another example is the *DXZ4* locus which is critical for determining the chromosomal organization in X chromosome inactivation (XCI)^45^. We tested *in silico* deletion of *DXZ4* locus in two female cell lines (IMR-90, GM12878) and two male cell lines (CUTLL1, Jurkat) to evaluate how *DXZ4* locus regulate X chromosome organization (Supplementary Fig. 25). Consistent with experimental knock-out results^45^, we found that deleting the *DXZ4* locus leads to substantial loss of insulation at the two flanking regions only in female cell lines (Supplementary Fig. 25), supporting the specific function of *DXZ4* locus in regulating XCI.

### Cell type-specific *in silico* genetic screening of *cis*-regulatory elements

Identifying *cis*-regulatory elements required for chromatin organization is one of the most important goals for 3D genome studies^46^. To determine whether C.Origami could be used to systematically identify such critical *cis*-elements, we propose using C.Origami to quantitatively assess how individual DNA elements contribute to the 3D chromatin organization (Fig. 5). Based on C.Origami’s model architecture, we developed two approaches for identifying critical *cis*-elements: a gradient-based saliency method named Gradient-weighted Regional Activation Mapping (GRAM), and attention scores derived from the transformer module (see Methods). As exemplified by the chr2:0-2.1Mb locus, both GRAM scores and attention scores captured important genomic regions that determine 3D genome structure, such as TAD boundaries and regions enriched with CTCF binding and ATAC-seq signals (Fig. 5c). In particular, GRAM can be positioned flexibly at different layers to obtain attribution maps at different resolutions up to nucleotide resolution (Supplementary Fig. 26a). The attention weights were averaged across all attention heads channels to obtain the layer-specific attention scores (Supplementary Fig. 26b). Visualization of all attention weights revealed that different attention heads attend to specific regions (Supplementary Fig. 27). Given that attention scores are robust to input shifts (Supplementary Fig. 26d), it is possible that the attention heads respond to specific categories of regulatory elements consistently. Additionally, we found although GRAM is more flexible, it is less robust compared to attention scores, susceptible to input window shifts and random seeds (Supplementary Fig. 26c-e). While both approaches are able to estimate the contribution of *cis*-elements, neither of them could quantitatively assess how much a specific DNA element influence the local 3D chromatin organization.

**Figure 5:**
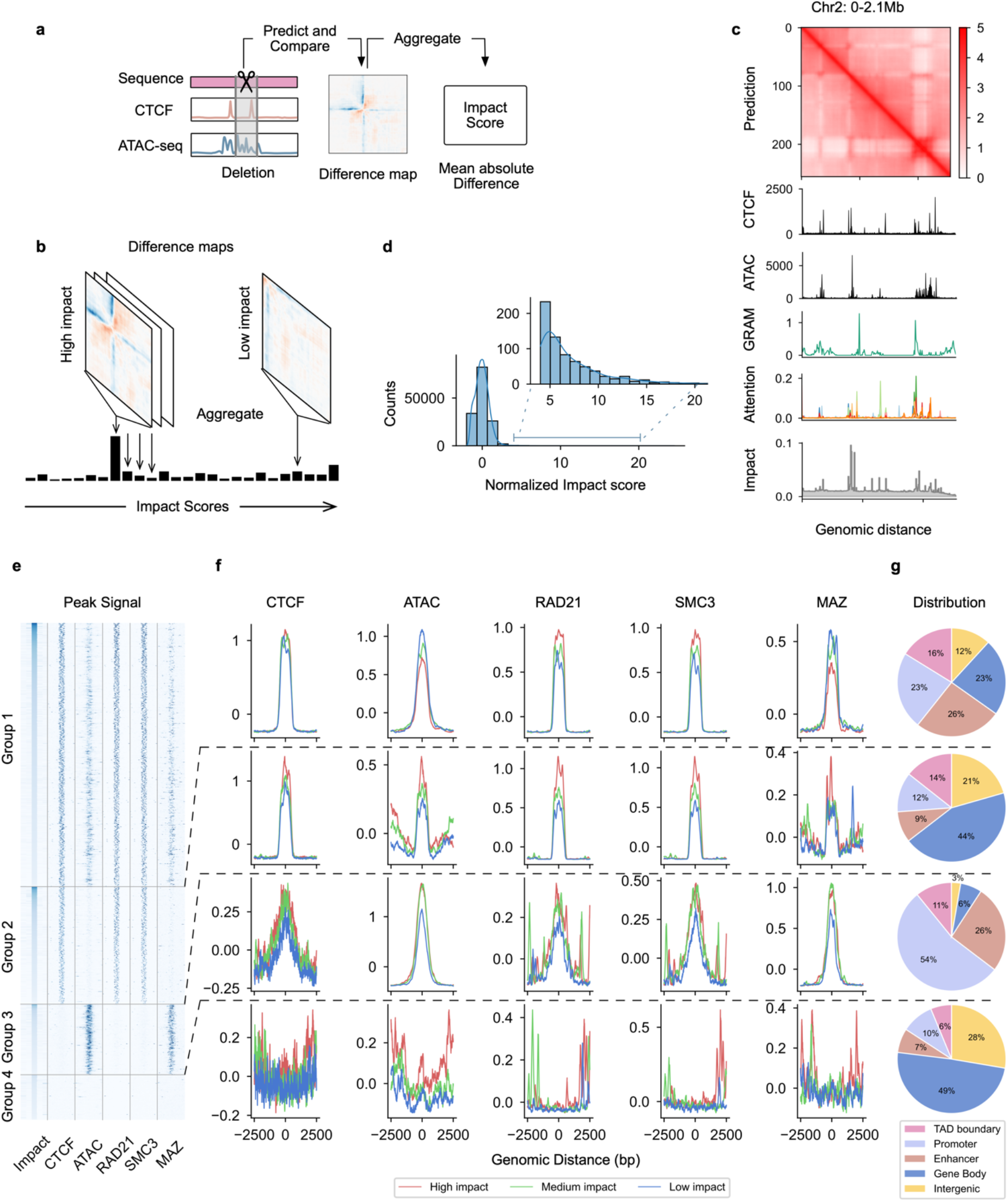
High-throughput *in silico* genetic screening identifies *cis*-regulatory elements determining chromatin organization. **a-b**, Schematic of *in silico* genetic screen for identifying impactful *cis-*regulatory elements. For each perturbed DNA element, an impact score is calculated to indicate how perturbation of the locus affected local chromatin organization. **c**, Visualization of different attribution method. GRAM, attention score, and impact score tracks are aligned to the predicted Hi-C and input genomic signals. **d**, Distribution of chromosome-wide-normalized impact scores in *in silico* deletion screening. **e**, Heatmap of *in silico* deletion screening-identified impactful *cis-*elements which contribute to the 3D chromatin organization. Each row shows an 5Kb locus centered by an impactful 1Kb *cis*-element. The loci in each group were ranked by their impact scores. **f,** The relative enrichment (z-score normalized) of ATAC-seq signal and multiple ChIP-seq signals at the four groups of impactful elements. According to the impact score values, *cis*-elements of each group were further grouped into high, medium, and low-impact quantile groups when plotting the ChIP-seq/ATAC-seq signals. **g**, Characterization of *in silico* screening-identified *cis*-elements by their genomic annotations.

Systematic DNA sequence perturbation was widely used in reverse genetic screening experiments for identifying functional genes or *cis-*regulatory elements. Inspired by the mechanism of genetic screening, we developed an *in silico* genetic screening (ISGS) approach based on C.Origami. Differ from qualitative GRAM and attention score, ISGS quantifies the difference in C.Origami predictions upon systematic perturbation (deletion) of DNA elements (see Methods). As an example, we first carried out ISGS in a 2Mb window (chr2:0-2.1Mb) by sequentially perturbing 256 loci of ∼8kb lengths, followed by Hi-C contact map prediction. We quantify the impact of a perturbation via a metric termed impact score, calculated by taking the mean absolute difference between predictions before and after perturbation (Fig. 5a, see Methods). We found that perturbations at TAD boundaries with enriched CTCF ChIP-seq and ATAC-seq signals had higher impact on chromatin folding, consistent with the GRAM and attention scores (Fig. 5c).

To systematically locate the impactful *cis*-elements that are required for 3D chromatin organization across the genome, we conducted high-resolution *in silico* genetic screening by sequentially deleting 1Kb DNA elements, followed by C.Origami prediction and impact score computation (see Methods). As expected, deletion of most of the DNA elements across the genome have low impact scores and does not significantly alter the 3D chromatin organization (Fig. 5d). We performed a peak calling by comparing each impact score to its surrounding signals, and isolated a set of impactful *cis-*elements representing ∼1% of the screened genome (see Methods).

Further characterization of the impactful *cis*-elements led to the identification of differential genomic features regulating chromatin organizations. According to the presence or absence of CTCF binding and ATAC-seq signals, the impactful *cis*-elements were characterized into four groups (Fig. 5e). More than half of the impactful *cis*-elements are open chromatin and simultaneously bound by CTCF (Group 1, Fig. 5e). Plotting CTCF binding signals and ATAC-seq signals across *cis*-elements in three quantiles separated by impact score group intensity, we found that CTCF-bound *cis-*elements intensity stays overall the same across Group 1 and Group 2 quantiles, while the ATAC-seq signals are negatively correlated with the impact scores (Fig. 5f, top). This result indicates that CTCF has a high impact on chromatin organization, regardless of the intensity of chromatin accessibility. Meanwhile, Group 1 and Group 2 *cis*-elements are enriched with RAD21 and SMC3 binding signals, supporting their function in defining boundaries during chromatin loop extrusion (Fig. 5f). Consistently, Group 1 and Group 2 elements enriched higher at TAD boundaries and enhancer-promoter regions (Fig. 5g). Notably, we identified a substantial fraction of *cis*-elements enriched in open chromatin, but are not bound by CTCF (Group 3). As expected, the ATAC-seq signal intensity is positively correlated with impact scores in Group 3 elements (Fig. 5f). Group 3 *cis*-elements are highly enriched in promoter and enhancer regions, indicating possible enhancer-promoter or promoter-promoter interactions^35^. We also found a small set of elements that are not related to CTCF and ATAC-seq signals (Group 4, Fig. 5e-g). Despite relatively lower impact scores, these elements may indicate alternative mechanisms which shape local 3D chromatin organization.

In addition, we sought to test whether additional factors could be enriched in the impactful elements for chromatin organization. Recently, Myc-associated zinc finger protein (MAZ) has been shown to co-localize with CTCF, thus may act as an additional architectural protein to organize chromatin structure^47,48^. To test this observation, we performed a similar enrichment analysis of MAZ ChIP-seq profile across the four groups of impactful elements (Fig. 5f). We found that MAZ is enriched in CTCF and ATAC-seq co-overlapped elements (Group 1), but not in the Group 2 elements where there is no open chromatin signal. Surprisingly, we found that MAZ is much more enriched in the open chromatin region where there is no CTCF binding (Group 3, Fig. 5e-f). This observation indicates that MAZ may function as a chromatin architectural protein independent of CTCF, acting at the active promoter-enhancer interaction regions.

### *In silico* genetic screening identified new T-ALL-specific chromatin organizations

Owing to C.Origami’s accurate cell type-specific prediction, we envisioned that the subsequent ISGS framework could empower systematic discovery of disease-specific chromatin organization. To systematically identify T-ALL-specific *cis*-elements, we performed ISGS and calculated impact scores across the genomes in CUTLL1 and Jurkat cells, in parallel with T cells (Fig. 6a, see Methods). Analyzing the impactful *cis*-elements between cell types, we identified both T-ALL-specific and T cell-specific elements (Fig. 6a).

**Figure 6:**
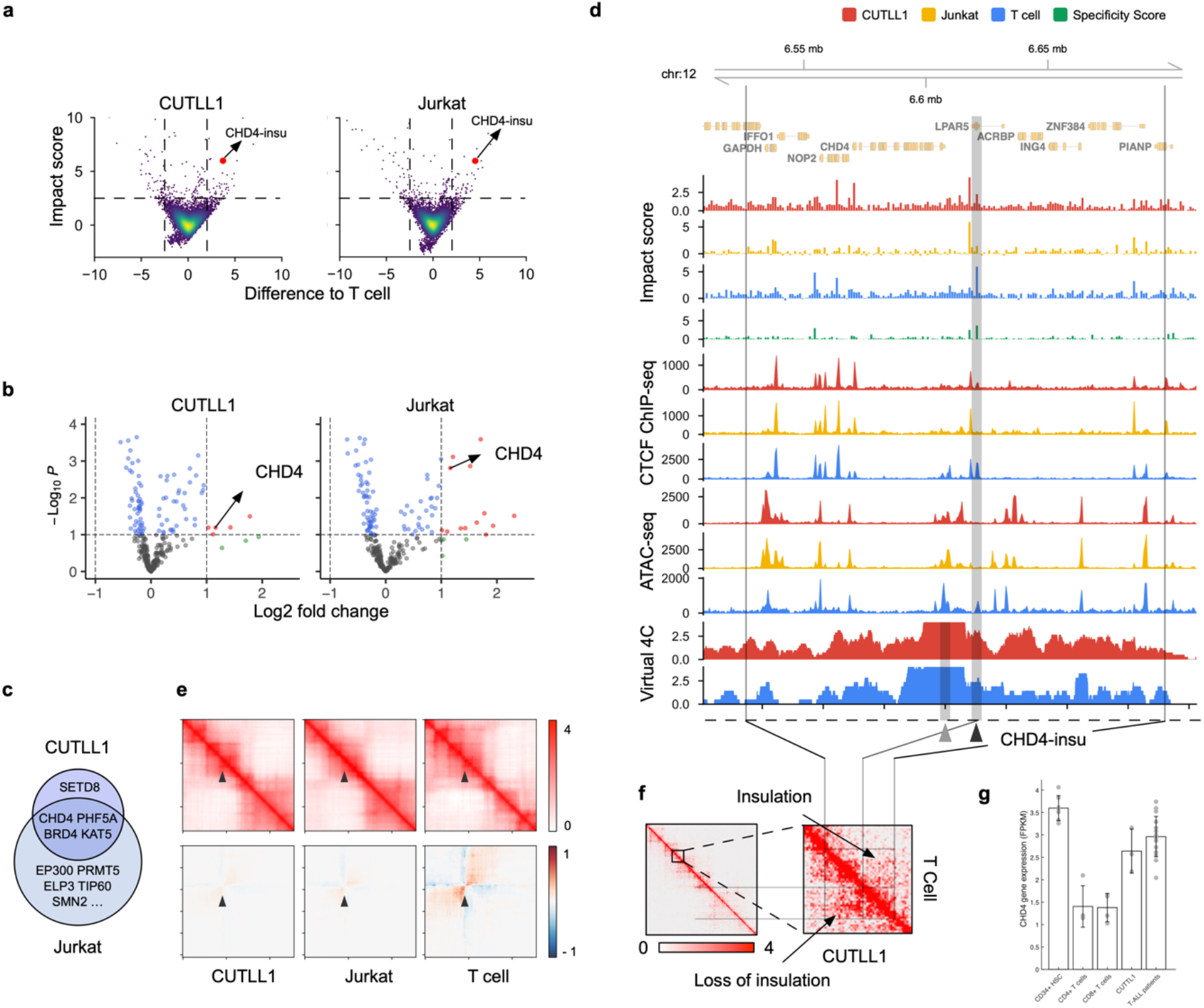
CRISPR and *in silico* genetic screen reveals T-ALL-specific chromatin interaction. **a**, A scatter plot showing impact scores of a sample of screened regions (n = 10000). The impact score difference between target cell type and T cell are shown on the x axis, and the higher impact scores between the corresponding cell type and the T cells are shown on y axis. The *CHD4-insu* locus is marked in red. **b**, Volcano plot of pooled CRISPR screening results on chromatin remodeling genes in CUTLL1 (left) and Jurkat (right) cell lines. The log2 fold changes indicate the normalized gRNA abundance in Day 4 versus Day 20 post-transfection, which reflect cell proliferation rate upon CRISPR targeting. Significant factors with log2 fold changes > 1 are marked in red. **c**, Overlap between CRISPR screening-identified significant chromatin-remodeling genes from CUTLL1 and Jurkat cells (**b**). **d**, Genomic tracks of 170Kb length around the *CHD4* locus. Presented genomic tracks include impact scores, CTCF ChIP-seq and ATAC-seq profiles, and virtual 4C signal using *CHD4* promoter as viewing point (highlighted by a gray band with gray arrowhead). The T cell-specific impactful *cis-*element, *CHD4-insu*, is highlighted with a gray band with black arrowhead. **e,** C.Origami prediction of *CHD4* locus (top row) and the difference (bottom row) upon deleting the *CHD4-insu* locus across cell types. The presented window represents chromosome 12: 6,620,219-6,621,219, with black arrowhead pointing to the *CHD4-insu* locus. **f,** Experimental Hi-C matrices of CUTLL1 (lower triangular region) and T cells (upper triangular region) at the *CHD4* locus. The presented region is aligned from the genomic track shown in **c**, highlighting T-ALL-specific interactions between *CHD4* promoter region and distal *cis-*elements. **g,** RNA-seq expression levels of *CHD4* in CUTLL1 cells, T-ALL primary patient samples, normal T cells and CD34+ hematopoietic stem cells. Error bars indicate one standard deviation.

Dysregulation of chromatin remodeling factors is frequently found in cancer cells^49,50^. We hypothesized that the dysregulation of local *cis*-regulatory elements around chromatin remodeling factors can lead to their abnormal expression in cancer. To connect the impactful *cis*-elements with critical chromatin remodeling genes in T-ALL, we first performed a pooled CRISPR knock-out screening in CUTLL1 and Jurkat cells, targeting chromatin remodeling factors that are required for T-ALL proliferation. This screening identified a set of genes, including *CHD4, PHF5A, BRD4* and *KAT5* as top hits important for T-ALL cell proliferation (Fig. 6b-c). By associating the ISGS-identified impactful elements with these four genes (Supplementary Fig. 28), we found an insulator element in the upstream region of *CHD4* gene, thereafter termed *CHD4-insu*, with a high impact score in T cells but a low score in T-ALL cells (Fig. 6d, black arrowhead. Also see Methods). Specifically, we found that the loss of CTCF binding at the *CHD4-insu* element might be responsible for the reduction of impact scores in T-ALL cells (Fig. 6d). Consistent with this observation, *in silico* deletion of the *CHD4-insu* element followed by C.Origami prediction in T cells led to loss of insulation and stronger interaction gain between the flanking regions compared to the effect in T-ALL cells (Fig. 6e).

CHD4 is the helicase component of NuRD complex, which functions to deacetylate H3K27ac^51^. Perturbation of CHD4 causes an arrest of cell cycle at G0 phase in childhood acute myeloid leukemia cells, indicating potential therapeutic target^52^. According to the *in silico* deletion experiment, we hypothesized that the loss of CTCF binding signal at the *CHD4-insu* locus leads to insulation loss. To test this hypothesis, we compared the experimental Virtual4C and Hi-C contact matrices of CUTLL1 and T cells (see Methods). As expected, we found that, compared to T cells, CUTLL1 cells have a higher interaction signal between the flanking regions of the *CHD4-insu* sequence, signifying higher interactions between *CHD4* promoter region and *cis-*regulatory elements in T-ALL cells (Fig. 6d virtual 4C tracks, and Fig. 6f). We further hypothesized that such increase of interaction affects CHD4 expression which is important for T-ALL proliferation. Supporting this hypothesis, RNA-seq experiment showed that *CHD4* expression is significantly upregulated in CUTLL1 cells and T-ALL patient samples compared to that in normal T cells (Fig 6g). These results indicate that the loss-of-insulation at the *CHD4-insu* element in T-ALL cells may have increased the expression of *CHD4* gene through establishing new chromatin interactions, consequently promoting leukemia cell proliferation. Together, our results demonstrated that the C.Origami-enabled ISGS framework is capable of identifying novel chromatin regulation mechanisms.

### Genome-wide *in silico* screening uncovers *trans*-acting regulators of chromatin folding

We next asked whether C.Origami-enabled *in silico* genetic screening could be leveraged for identifying cell type-specific *trans*-acting regulators determining the 3D chromatin organization. We first conducted chromosome-wide *in silico* deletion screening to identify cell type-specific impactful loci that were critical for predicting chromatin organization (see Methods). High-impact 1Kb regions were then annotated and tested for enrichment in transcription factor binding profiles from the ReMap database^53^. Odds ratio for binding potential was calculated for each factor, followed by normalization within each cell type. The normalized odds ratio scores enable characterization of differential *trans*-acting regulators across cell types (see Methods).

Applying this framework to the two T-ALL cell lines and T cells, we found differential compendia of transcription factors contributing to the cell type-specific 3D genome organization (Fig. 7a, Supplementary Fig. 29). Scoring these *trans*-acting regulators across cell types, we identified different categories. Notably, our analysis consistently identified known 3D chromatin regulators, such as CTCF, RAD21 and SMC1/SMC3, as top candidates across cell types (Category 1, Fig. 7a). In addition, we found differential sets of *trans*-acting regulators enriched in T cells and T-ALL cell lines, respectively. Several known factors critical for T cell function (Category 2), such as RCOR1, SMAD3 and ZEB2, are enriched in the T cell-specific group of *trans*-acting factors (Fig. 7a). Consistently, CUTLL1 and Jurkat cells enriched similar groups of *trans*-acting factors (Category 3), represented by MAZ, BRD2, and NOTCH1 (Fig. 7a).

**Figure 7:**
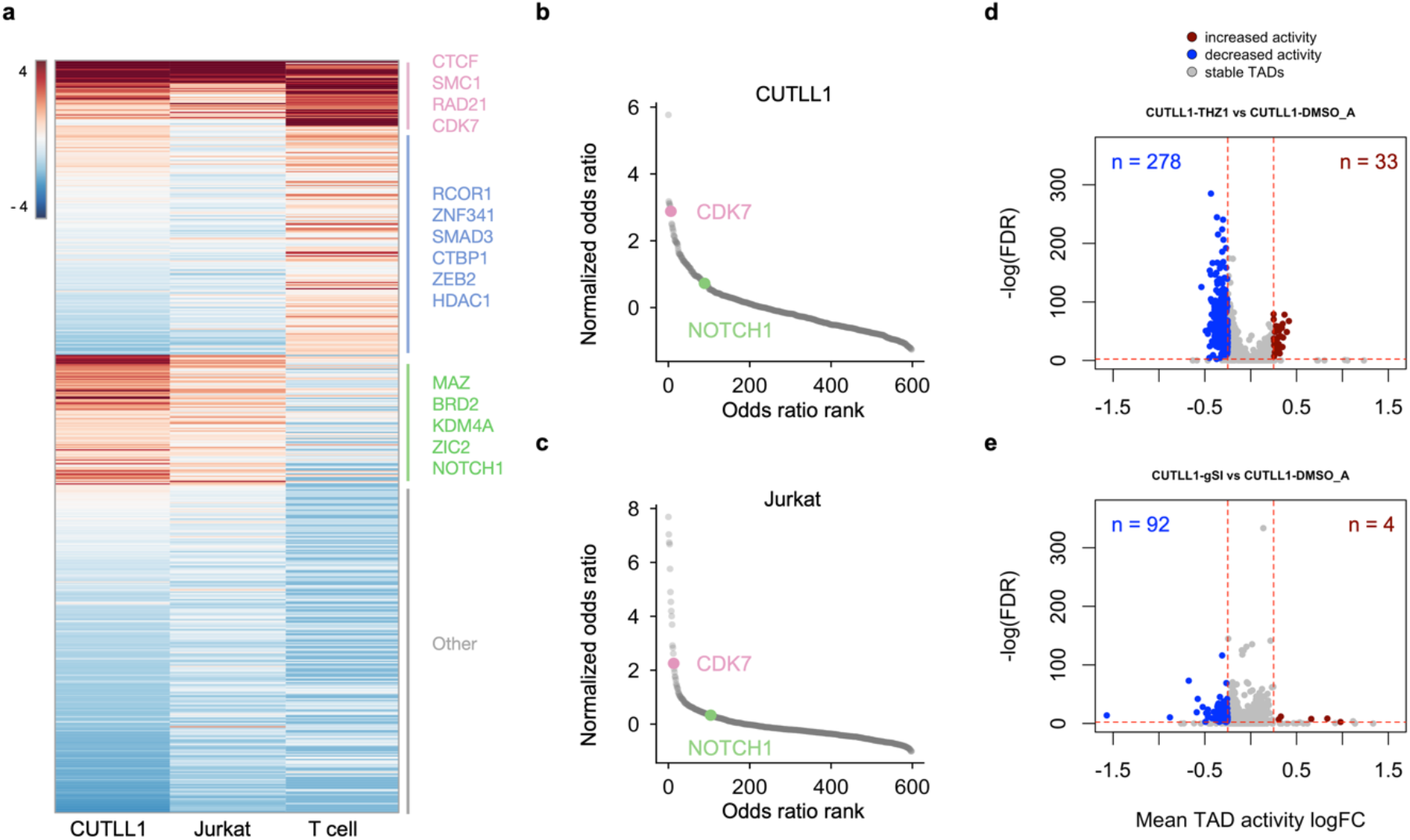
*In silico* genetic screening uncovers *trans-*acting regulators of chromatin folding. **a,** A heatmap of normalized odds ratio scores of the enrichment of *trans*-acting regulators across cell types. Representative factors are listed next to the three major categories. **b-c**, Elbow plots of *in silico* genetic screening-identified *trans*-acting regulators in CUTLL1 cells (**b**) and Jurkat cells (**c**). CDK7 and NOTCH1 are highlighted in both plots. **d-e,** Volcano plots showing chromatin organization changes of individual TADs upon pharmacological inhibition of CDK7 (**d**) or NOTCH1 (**e**) in CUTLL1 cells. Each dot represents a TAD varied from ∼200Kb to 3Mb.

Previously, we found that both CDK7 and NOTCH1 regulate enhancer-promoter interactions in T-ALL cells^30^. Pharmacological inhibiting NOTCH1 leads to H3K27ac alterations in a subset of NOTCH1-associated chromatin interactions, while inhibiting CDK7 leads to broader changes in H3K27ac, indicating that CDK7 may have a broader impact on 3D chromatin organization^30^. To further test the hypothesis that pharmacological inhibiting CDK7 leads to broader chromatin organization changes for controlling T-ALL cell proliferation, we systematically assessed the relative contribution of *trans*-acting factors through *in silico* genetic screening in CUTLL1 and Jurkat T-ALL cells. Consistent with our prior results, we found that CDK7 ranked among top factors in regulating 3D chromatin organization, whereas NOTCH1’s predicted contribution was ranked much lower (Fig. 7b-c). Supporting the results inferred from ISGS analysis for *trans*-acting regulators, we found that pharmacological inhibition of CDK7 (+THZ1) leads to more TADs with chromatin organization changes than the effect from inhibiting NOTCH1 (+γSI) in CUTLL1 cells (Fig. 6d-e). Moreover, we found that impactful elements are more enriched in TADs with significant intensity changes upon CDK7 inhibition (Supplementary Fig. 30).

## Discussion

Cell type-specific gene expression requires specific chromatin folding patterns. In this study, we developed a multimodal deep neural network architecture, C.Origami, that incorporates both DNA sequence and genomic features for *de novo* prediction of cell type-specific 3D genome organization (Fig. 1). We found that DNA sequence information together with CTCF binding signal alone was not sufficient for accurate *de novo* prediction of cell type-specific chromatin organization, whereas incorporating chromatin accessibility data into C.Origami provided the model with sufficient information to achieve prediction results comparable to high-quality Hi-C experiments (Fig. 2-3). These results are consistent with the observation of widespread transcription-associated chromatin interactions at the accessible chromatin regions^33,34^. Systematic ablation study further showed that the specific input combination of DNA sequence, CTCF binding, and open chromatin features enables the best prediction result (Fig. 2a).

The rules governing 3D chromatin organization is consistent across different cell types, even between human and mouse. Although C.Origami was trained only using IMR-90 cell data, its ability to learn from one cell type and extrapolate prediction to other unseen cell types implies that the 3D chromatin organization rules learned by C.Origami is applicable to the general mammalian genome. We found that C.Origami achieved a general high performance in predicting cell type-specific TAD structures. In addition, the predicted results can further be applied for detecting various types of chromatin loops across cell types. Due to its sensitivity to input data noise and quality difference in the public datasets, we expect future development of the C.Origami model would further improve chromatin loop detection performance from prediction by incorporating a customized normalization of input information.

The high performance and minimal requirement on cell type-specific input data make C.Origami feasible for studies requiring frequent *de novo* analysis of 3D chromatin organizations without performing Hi-C experiments (Fig. 4). Similar to high-resolution Hi-C data, the predicted chromatin contact matrices can be directly analyzed by other downstream computational tools for inferring TADs, chromatin loops, and enhancer-promoter interactions^54–56^. C.Origami can be useful in fields such as cancer genomics involving widespread genome rearrangement and synthetic regulatory genomics with *de novo* regulatory circuit construction^8,40,57,58^.

With highly accurate prediction of chromatin organization, our model enables *in silico* genetic perturbation as a tool to study how *cis-*elements determine 3D chromatin organization in a cell type-specific manner. Given data from genomic features and Hi-C map, it is challenging to establish the causal relationship between differential genomic features and chromatin organization changes. C.Origami can accurately simulate the changes in chromatin organization upon *in silico* genetic perturbation, providing an effective way to map the causal relationship between genomic regions and chromatin organizations. *In silico* perturbation can be performed within seconds and is much more efficient compared to traditional experiments. Expanding the throughput of *in silico* genetic perturbations, we demonstrated the efficacy of *in silico* genetic screening framework for identifying critical DNA elements determining 3D chromatin organization (Fig. 5). While multiple previous methods, such as Expecto^59^, BPNet^60^ and Enformer^61^, have been developed to identify functional *cis-*regulatory elements, none of these methods could identify the cell type-specific chromatin interactions between those functional DNA elements. The *in silico* genetic screening allowed us to categorize different groups of *cis-*regulatory elements that are importance for 3D chromatin organization, including those only bound by CTCF or CTCF-free open chromatins. These differential genomic features may indicate distinct types of chromatin interactions, ranging from CTCF-dependent structural organization though loop extrusion to transcription-associated chromatin looping bound by MAZ^10,47,48^.

We demonstrated the power of *in silico* genetic studies of 3D chromatin organization in leukemia. Screening for differential impactful *cis*-elements between T-ALL cells and normal T cells, we found a loss of insulation event at the upstream of *CHD4* gene in T-ALL cell lines (Fig. 6). Such loss of insulation induced new chromatin interactions between *CHD4* promoter and distal *cis*-elements, correlating with gene expression level changes in T-ALL cells (Fig. 6). Notably, CHD4 has been found critical for cell growth in childhood acute myeloid leukemia^52^. The discovery of a T-ALL-specific *CHD4* gene expression regulation hints a potential anti-leukemia target by perturbing *CHD4* gene expression. Moving beyond, disruption of chromatin organization insulations has been identified through extensive experimental studies^30,62,63^. We envision that *in silico* genetic screening framework could be generally applicable for identifying critical *cis*-regulatory elements across biological systems.

Last, through systematic *in silico* screening followed by integrative analysis with TF-binding databases, we could compile a compendium of potential *trans*-acting regulators determining the chromatin organization in a cell type-specific manner. Analyzing *trans*-acting regulators in T-ALL samples, we provide direct evidence that CDK7 plays a broader role in modulating 3D chromatin organization than NOTCH1, consistent with our previous results^30^. As the number of CTCF ChIP-seq and ATAC-seq grows for new cell types, we expect the model to be capable of identifying cell type-specific features through their predicted chromatin structure and trans-acting regulators. Application of *in silico* screening across normal and disease conditions could lead to the identification of novel targets for therapeutics.

By integrating cell type-specific genomic features and DNA sequence information, we demonstrated that C.Origami can predict complex genomic features and enables *in silico* genetic perturbation and screening with high accuracy. We expect the underlying architecture of our model, Origami, is generalizable for applications across a broader range of genomic features, such as epigenetic modifications and gene expression. We expect future genomics study to shift towards using tools that leverage high-capacity machine learning models like Origami to perform *in silico* experiments for discovering cell type-specific genomic regulations.

## Acknowledgements

A.T. is supported by the NCI/NIH P01CA229086, NCI/NIH R01CA252239, NCI/NIH R01CA260028 and NIH/NCI R01CA140729. B.X. is supported by NIH DP5OD033430. I.A. is supported by the NIH R01CA266212, R01CA242020, R01CA228135 and P01CA229086. JS is supported by the R35GM122515, P01 CA229086 and P30CA016087. We would like to thank the Genome Technology Center (GTC) for expert library preparation and sequencing, and the Applied Bioinformatics Laboratories (ABL) for providing bioinformatics support and helping with the analysis and interpretation of the data. GTC and ABL are shared resources partially supported by the Cancer Center Support Grant P30CA016087 at the Laura and Isaac Perlmutter Cancer Center. This work has used computing resources at the NYU School of Medicine High Performance Computing (HPC) Facility. We would like to thank Sudarshan Pinglay, Jef Boeke, Huiyuan Zhang, and the members of the Tsirigos lab for suggestions and discussion.

## Author contribution

J.T. and B.X. conceived the project. J.T., B.X. and A.T. designed the experiments and interpreted the results. J.T. designed, implemented and optimized the neural network, and performed all the downstream computational analysis with help from J.R. N.S. and T.S. contributed to the public code repository. E.W. performed the CRISPR screening experiments. F.B. generated ATAC-seq for CUTLL1. J.T. prepared figures with inputs from B.X., A.T. and D.F. T.S., P.T., J.S., I.A. and D.F. contributed to discussion. B.X., J.T. and A.T. wrote the manuscript with input from all authors.

## Methods

### Hi-C data and processing

We used seven human and mouse Hi-C profiles in this study: IMR-90, GM12878, H1-hESC, K562, CUTLL1, T cell, Mouse Patski (Supplementary Table 1). All the data are available on GEO (www.ncbi.nlm.nih.gov/geo) and/or 4D Nucleome Data Portal (https://data.4dnucleome.org). To minimize bias in Hi-C data preprocessing, we obtained counts data in raw fastq format. The reads from human cell lines were aligned to GRCh38 human reference genome and mouse cell lines are aligned to mm10 mouse genome. The alignments were filtered at 10kb resolution and iteratively corrected with HiC-bench^64^. To ensure the compatibility of prediction result with downstream analytical tools, we only used a reversible natural log transform to process the Hi-C prediction targets. Prediction from C.Origami with exponential transformation can be directly used as Hi-C chromatin contact matrix data for any downstream analysis.

**Supplementary Table 1.** Hi-C data used for training and validation.

| Cell Type | Enzyme | Accession Number | Reference |
| --- | --- | --- | --- |
| IMR-90 | MboI | GSE63525 | Rao et al.<sup>32</sup> |
| GM12878 | MboI | GSE63525 | Rao et al.<sup>32</sup> |
| H1-hESC | Arima | 4DNESFSCP5L8 | Calandrelli et al.<sup>65</sup> |
| K562 | MboI | GSE63525 | Rao et al.<sup>32</sup> |
| CUTLL1 | Arima | GSE115896 | Kloetgen et al.<sup>30</sup> |
| T cell | Arima | GSE115896 | Kloetgen et al.<sup>30</sup> |
| Mouse Patski | Arima | GSE71831 | Darrow et al.<sup>45</sup> |
| Mouse ESC | HindIII | GSE98671 | Nora et al.<sup>38</sup> |

### CTCF ChIP-seq and ATAC-seq data

CTCF ChIP-seq and ATAC-seq data for all cell-types are publicly available online from GEO (www.ncbi.nlm.nih.gov/geo) and ENCODE data portal (www.encodeproject.org/). CUTLL1 ATAC-seq was sequenced according to standard method^66^. Details on accession number are listed in Supplementary Table 2. To maintain signal consistency across different cell lines, we aggregated fastq data from different replicates and subsampled them down to 40 million reads. The reads were processed by Seq-N-Slide to generate bigWig files (https://doi.org/10.5281/zenodo.6308846). The bigWig was used as regular, dense inputs to our model. To prepare an alternative sparse input format, we used MACS2 to perform peak calling on the intermediate bam files to obtain sparse peaks for CTCF and ATAC-seq^67^. The sparse narrowPeak file was converted back to bigWig with ucscutils. We took the natural log transformation of both dense and sparse bigWig files and used them as inputs to the model.

**Supplementary Table 2.** CTCF ChIP-seq and ATAC-seq used for training and validation.

| Cell Type | CTCF ChIP-seq | ATAC-seq |
| --- | --- | --- |
| IMR-90 | ENCSR000EFI | ENCSR200OML |
| GM12878 | ENCSR000AKB | ENCSR095QNB |
| H1-hESC | ENCSR000AMF | GSE85330 |
| K562 | ENCSR000AKO | ENCSR483RKN |
| CUTLL1 | GSE115893 | see Methods CUTLL1 |
| Jurkat | GSE115893 | GSE90718 |
| T cell | GSE115893 | GSE168880 |
| Mouse Patski | ENCSR419OOD | ENCSR351QUO |
| Mouse ESC | GSE98671 | N.A. |

### DNA sequence

We used the reference genome sequence (hg38 and mm10) from UCSC genome browser database. The original fasta file includes four types of nucleotides and “n” for unknown type. We retained the ‘n’ category and encoded it as the unknown fifth ‘nucleotide’. After encoding, each nucleotide is a 5 channel one-hot vector representing ATCGN. The same reference genome sequence was used for all cell types.

### Training data

The training data consists of DNA sequence, CTCF binding signal, ATAC-seq signal and Hi-C matrix from IMR-90 cell line. The input data to the model includes DNA sequence, CTCF ChIP-seq signal, and ATAC-seq signal at a 2,097,152 bp region. The output target is the Hi-C matrix at the corresponding region. The Hi-C matrix was originally called at 10Kb resolution and downscaled 8,192 bp to match the model output resolution. To generate batches of training data, we defined 2Mb sliding windows across the genome with 40Kb steps. Windows that have overlap with telomere or centromere regions were removed. We split the genome into training, validation and test chromosomes. Chromosome 10 and 15 were used as the validation set and the test set respectively. The rest of the chromosomes were used as the training set.

### Model architecture

C.Origami is implemented with the PyTorch framework. The model consists of two 1D convolutional encoders, a transformer module and a 2D convolutional decoder to adapt to input channels of sequence and genomic features. The sequence encoder has five input channels, and the genomic feature encoder has two input channels. The two encoders have similar structures otherwise. To reduce memory cost, each encoder starts with a 1D convolution header with stride 2 to half the size of the 2Mb bp input before it goes to convolution blocks. To reduce the input length down to 256, we deployed twelve convolution modules, each of which consists of a residual block and a scaling block. The residual block has two sets of convolution layers with kernel width 5 and same padding. Batch normalization and ReLU nonlinearity follows each convolutional layer, and the start and end position of the residual block is connected by a residual connection. The residual blocks do not alter dimension of inputs. The skip-connections within the residual block help promote information propagation. The scaling block consists of a 1D convolutional layer with kernel size 5 and stride 2 followed by batch normalization and ReLU activation. The scaling block reduces input length by a factor of 2 and increases the number of hidden layers. We increase the hidden size according to this schedule: 32, 32, 32, 32, 64, 64, 128, 128, 128, 128, 256, 256. The output from the last scaling module has a length of 256 with 256 channels.

The transformer module is crucial for the model to encode dependencies across input elements at different positions. The module is built with eight customized attention layers similar to a BERT model^68^. Specifically, we set the number of hidden layers to 256, ReLU as the activation function and used eight attention heads. We used relative key query as positional embedding and set the maximum length to be 256.

After the transformer module, the model concatenates each position in the 256 bins to every other position to form a 256 by 256 interaction map. The concatenation function takes the 256-bin sequence from the feature extraction module and outputs a 256-by-256 grid where location (I, j) is a concatenation of the features at i and j position. Then a 1-dimensional distance matrix is calculated and appended to the grid. The distance matrix value at location (i, j) is the Manhattan Distance between point (i, i) and (j, j) on the grid divided by 2. Since each bin has 256 channels, after concatenation and addition of the distance matrix, we arrived at an output of 256-by-256 grid with 513 channels.

The decoder consists of five dilated residual networks. We designed the dilation at the corresponding layer to be 2, 4, 8, 16, 32 so that the receptive field of each pixel at the last layer covers the input space, reinforcing interactions between different elements. At the end of the decoder, we use a Conv2D layer with 1×1 kernel to combine 256 channels down to one channel and the output is a 256×256 matrix with one channel.

The 256×256 output from the model was compared with experimental Hi-C map (ground truth) via a mean squared error (MSE) loss. The loss was back propagated through the whole network for gradient updates.

### Data augmentation

To avoid overfitting, we implemented three types of data augmentations. First, during training, we dynamically selected the 2Mb window with random shifts between plus and minus 0.36 Mb range. Second, we reverse-complemented the sequence and flipped the target Hi-C matrix with 0.5 probability. Third, we added Gaussian noise to sequence, CTCF ChIP-seq and ATAC-seq signals with zero mean and 0.1 standard deviation.

### Model training and prediction

To train the model, we used a training batch size of 8 and Adam optimizer with learning rate 0.002. The cosine learning rate scheduler with 200 epoch period was used for stabilizing training. The model achieved minimal validation loss when trained for 54 epochs. The model training time was 18 hours on a GPU cluster with 4 NVIDIA Tesla V100 GPUs with 320GB RAM to store training data. To prevent bottlenecking from data loading process, we used 8 CPU workers to load data and assigned 10 CPU cores in total for the training procedure. Model inference with a mobile NVIDIA RTX 2060 GPU can be achieved in under 1 second, and inference on an Intel i7-8750H CPU is around 3 seconds. To run prediction in IMR-90, the reference DNA sequence, CTCF ChIP-seq and ATAC-seq from IMR-90 in a 2Mb region are taken as input. For *de novo* prediction in a target cell type, we replaced IMR-90 CTCF ChIP-seq and ATAC-seq with corresponding CTCF and ATAC-seq from the specific target. The reference sequence is kept the same.

### Insulation score

Insulation score is implemented as the ratio of maximum left and right region average intensity and the middle region intensity^64^. We also added a pseudo-count calculated from chromosome-wide average intensity to prevent division by zero in unmappable regions. Given that all the regions contain *n* interactions, the insulation score can be formulated as follows:

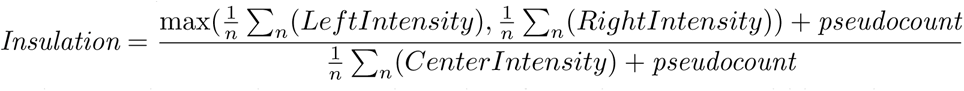

where pseudo-count is set to the average intensity of one chromosome within 2mb.

### Loop calling

We used the Hi-C valid pairs with the FitHiC software^69,70^ to identify significant interactions. We used a resolution of 10kb, minimum and maximum distance of 30kb and 1Mb. For loop calling on predicted matrices, we converted the predicted matrix back to valid pairs by merging predictions to chromosomes and counting the discretized intensity value. FitHiC generated a list of significant interactions with corresponding FDR corrected q-values. For loop analysis on IMR-90, we computed AUROC and overlap between loops called from experimental Hi-C and loops called from predicted Hi-C. To calculate AUROC, we used predicted loops as target. Q-value cutoffs ranging from 1e-5 to 1e-13 are selected to filter significant loops called from the predicted Hi-C. Then, the q-values from loop called from experimental Hi-C were compared to significant loops called from prediction to calculate an AUROC. For overlap analysis, we chose a fixed 1e-5 cutoff for loops called from predicted and experimental Hi-C and compared the overlap of significant loops. For loop analysis on specific type of interaction, we overlapped the two anchors of each loop and obtain the categories for each loop called. The loops were then filtered by different categories and the same AUROC and overlap analysis was performed on each category of loops.

For cell type-specific loop analysis between IMR-90 and GM12878, we first used a more stringent cutoff of 1e-7 as a threshold for significant loops. Then we further categorize specific loops into IMR-90 specific or GM12878 specific according to the log2 fold change (log2fc) of loop interactions counts. To calculate AUROC, we used log2 fold change in place of the q-value cutoff from previous analysis. We compared two log2fc. The first log2fc is between predicted loops in cell type 1 and predicted loops in cell type 2 (e.g. IMR-90 predicted loop / GM12878 predicted loop). The second log2fc is between experimental loops in cell type 1 and predicted loops in cell type 2 (e.g. IMR-90 experimental loop / GM12878 predicted loop). Then the same AUROC and overlap analysis was performed for each of the two cell type-specific groups. For loop analysis on specific type of interaction in a cell type-specific way, the same anchor overlap was performed with corresponding AUROC and overlap analysis.

### Chromosome-scale Hi-C contact matrix prediction

To bridge adjacent 2Mb-window predictions into chromosome-wide Hi-C contact matrices, we ran the prediction in a sliding window of step side 262,144 bp, which is 1/8 of the 2Mb prediction window. All predictions were in-painted to their corresponding location on the contact map. Most regions were covered by prediction for 8 times except for regions at the beginning or end of the chromosome. To correct for different levels of overlap, we counted the total times of overlap for every pixel and applied corresponding scaling factors. The resulting chromosome-wide prediction can be directly used for downstream analysis such as TAD calling and insulation score calculation.

### Distance-stratified intensity and correlation

Distance-stratified intensity and correlation calculation were based on fused chromosome prediction. Stratified intensity at distance i was calculated by aggregating the line that is parallel to the diagonal with offset of i. Stratified correlation was calculated as Pearson’s *r* between the shifted diagonal line of prediction and ground truth.

### Performance comparison with previous methods

We compared performance of C.Origami against three previously published methods: Akita^22^, DeepC^23^, and Orca^36^. We compared the performance using four metrics: insulation score correlation, observed vs expected Hi-C metrices correlation, mean squared error (MSE), and distance-stratified correlation. We calculated the four metrics separately for the four models by their prediction to the experimental data as ground truth. The comparison were carried out in two different cell types: 1) in the training cell type, IMR-90 cell, which most models were trained on, and 2) in a new cell type, GM12878 cells, aiming to quantify the performance of *de novo* prediction of chromatin organization of the four models.

We generated a set of sliding windows that covers the whole genome and can be predicted by each model. Since Akita and DeepC are only able to predict interaction within a 1Mb window, we restricted the test regions to 1Mb blocks. To generate a genome-wide testing dataset, we selected all 1Mb regions in a sliding window with 0.5Mb overlap between neighboring regions. To ensure compatibility with all models’ prediction windows, the first 1.5Mb and last 1.5Mb of chromosomes were used as buffer regions for models requiring 2Mb windows as inputs. In total 5935 regions were generated. The Hi-C experimental data was extracted from these regions as targets. We used all 4 models to predict the interaction for the corresponding regions.

The most relevant versions of the previous models were selected for comparison. For Akita, the IMR-90 output channel was selected. For DeepC, we used their model trained with IMR-90 data. Orca was only trained on HFF and H1-hESC. We used the HFF model because HFF is also a fibroblast cell line similar to IMR-90. The comparison turned out to be valid because even though Orca was trained on HFF, it outperformed both Akita and DeepC on IMR-90 in many benchmarks. For C.Origami, we used the IMR-90-trained model.

It is necessary to perform scaling and normalization to each models’ outputs due to their varied prediction target customizations. Akita predicts a 1048576bp region with 512 bins. We removed the extra 48576bp on the sides to make the prediction 1Mb, followed by rescaling bins into 128. Orca can predict interactions at multiple scales. Since C.Origami used a 2Mb window as prediction target, we selected the 2Mb window in Orca for consistency. The prediction was then cropped to 1Mb and rescaled to 128 by 128. For C.Origami, the prediction is a 2097152 bp window. We cropped the prediction to leave the center 1Mb regions and rescaled the bins to 128.

DeepC’s prediction target is different from other models with predictions of 45-degree rotated version of the Hi-C matrices. DeepC also produces predicted Hi-C maps in different scales compared with other methods. Thus, we performed a series of transformations (Supplementary Figure 11) including mirroring, rotating and cropping to make a comparable contact matrix to outputs produced by other models. We used a 1Mb prediction window for DeepC and rescaled the output to 128 by 128.

The first step to make the models comparable is selecting a common ground truth Hi-C as the evaluation target. Since each model used a different ground truth with different transformations (e.g. obs/exp, log, gaussian smoothing), they cannot be compared directly. We defined the evaluation target as logged Hi-C intensity (log(ICE normalized counts + 1)). Logged intensity has a few advantages over observed vs expected map. First, it allows for computing insulation scores. Second, it can be converted to observed vs expected while the reverse is not straightforward. It can also be converted to raw counts by taking the exponent. Third and most importantly, it is used as the default Hi-C format for most downstream analysis pipelines like loop calling, and visualization.

The second step to make the models comparable is to normalize model outputs to the evaluation Hi-C target. Since each model used a different original prediction target, we want to measure the difference between the original target and the evaluation target. We plotted the mean/std of intensity over distance between prediction and evaluation target and found a large discrepancy between models. Specifically, DeepC results stood out with a unique pattern that might be a result of their custom stratified binning method (Supplementary Figure 12). We also observed that the raw predicted matrix intensities were too different to compare (Supplementary Figure 12).

We performed distance-stratified normalization (DSN) to align all predictions to the target prediction (Supplementary Figure 13). We computed the mean and std for each diagonal and then normalized the prediction to target experimental Hi-C. Formally, let *T̂* be the normalized matrix, *T* be the target ground truth matrix, and *M* be the unnormalized matrix. Let *m_d,i_* be the corresponding element in *M* and *µ, σ* denote the mean and std at diagonal *d* in matrix *T* and *M*. Then, every *i^th^* entry on *d^th^* diagonal *t_d,i_* can be normalized as follows.

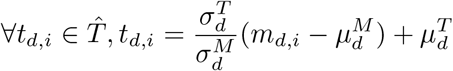

The normalized prediction were compared to the target Hi-C using four metrics: insulation score correlation, obs/exp Hi-C matrix correlation, MSE (mean squared error), and distance-stratified correlation. Each metric was calculated per chromosome for every tested model using their corresponding prediction and the experimental data as ground truth.

We also performed GM12878 *de novo* prediction comparison. For C.Origami, we used the same IMR-90 trained model but GM12878 CTCF ChIP-seq and ATAC-seq profiles as inputs to predict Hi-C. For sequence only models, we used the same DNA sequence setup because they could not provide cell type-specific *de novo* prediction. Though ideally input DNA sequence should be cell type-specific, such procedure is not realistic for general applications.

### C.Origami prediction at the CUTLL1 t(7;9) translocation site

To generate experimental Hi-C data, we defined a custom chromosome in Hi-C bench^64^. The custom genome in Hi-C bench is defined at the matrix-filtered step where the pipeline assign reads to chromosomes. For CUTLL1 experiment, we defined a custom chromosome chr7chr9 with chr7: 0-142800000 as the starting chromosome and chr9: 136500000-138394717 as the ending chromosome.

CUTLL1 t(7;9) translocation is heterozygous, leading to allele-specific complexity to its corresponding Hi-C matrix. Since only one allele is translocation, the experimental Hi-C data mapped to either normal reference genome or the t(7;9) translocated reference genome would be a mixture of chromatin interactions from both translocated and normal chromosomes. To align with this hybrid effect of Hi-C contact map, we first separately predicted three sets of Hi-C maps: t(7;9) translocated chromosome, normal chromosome 7, and normal chromosome 9. The predicted Hi-C matrix at the t(7;9) locus is an average of the predicted Hi-C maps of t(7;9) translocation chromosome and a fused prediction map ranging from normal chr7 to the breakpoint chr7:142,797,952 and extending from chr9:136,502,817 to the rest of normal chr9. We did not count the inter-chromosomal interactions at these loci due to their much weaker intensity compared to the intra-chromosomal interaction at the translocation site.

### Mouse prediction

For the mouse Patski cell type prediction^45^, the CTCF ChIP-seq and ATAC-seq inputs were processed using the same pipeline with mm10 as the assembly number. The original C.Origami model trained with IMR-90 dense input features was used for prediction. For genome-wide evaluation of predicting mouse chromatin organization, we adopted the same procedure from the “Performance comparison with previous methods” section.

### CTCF depletion prediction in mESC

We preprocessed CTCF ChIP-seq and Hi-C on mouse ESC cells from Nora et al^38^ following the same pipeline for ChIP-seq and Hi-C. In total, three sets of data with conditions: untreated, auxin-induced CTCF depletion, and wash-off are processed. Since this study did not measure ATAC-seq signal, C.Origami model was re-trained using only DNA sequence and CTCF ChIP-seq on the untreated condition. The re-trained model was then used for predicting chromatin organization in the CTCF depletion (auxin treatment) and restoration (auxin wash-off) conditions. Genome-wide performance benchmark followed the same procedure as in the “Performance comparison with previous methods” section.

### Reducing impact score from 3D voxels

Screening by deletion produces a 3D voxel with coordinates (i, j, k) where the first two dimensions (i, j) correspond to the Hi-C matrix difference and the third dimension k denotes deletion locus. Under this framework, the impact score can be defined as reducing the first two dimensions (i, j) with mean or sum, denoting the overall intensity shift with respect to deletion. The sensitivity score can be defined as the result of reducing either of the first two dimensions (i or j) and the third deletion dimension k. From another perspective, sensitivity score of a locus denotes average intensity shift over all deletions with respect to its location.

### GRAM (Gradient-weighted Regional Activation Mapping)

The GRAM scoring system is a generalized version of Grad-CAM on 2D outputs^71^. Instead of taking a single output, GRAM operates on a region *r* in the output space *y* and runs backpropagation on all pixels within *r*. GRAM on region *r* in network layer *m* is defined as follows:

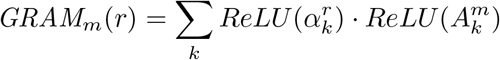

where 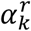 is the activation weight for channel *k* and region *r*. Formally, 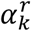 is defined as:

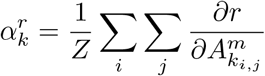

where *Z* is the number of activations in the layer and the quotient is the gradient at position *i*, *j* in the activation layer *m* with respect to output *r*. 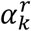 can be interpreted as the average gradient across the *i, j* (width and height) dimension at the layer *m*. 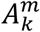 is the activation in channel k at layer *m*. In this study, we choose *r* to be the full output space. During forward propagations, activation (*A^m^*) at the target layer *m* is recorded. This activation map is a 3D tensor, or an image with k channels. Then, the *r* region of the output is selected for backpropagation and gradients are calculated for every layer. The gradients (used for calculating weights 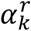) at the target layer *m* are collected. The set of collected gradients is also an image-like 3D tensor with k channels. To obtain 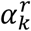, we averaged the gradients across width and height dimension, resulting in a k-dimensional array. The goal of GRAM is to visualize a gradient-weighted activation map that maximizes the output signal. To obtain this weighted activation, 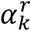 is used as weights to average the k channels activation image (*A^m^*). The final averaged activation is defined as the GRAM output.

### Attention score

In the transformer module, we implemented the vanilla multi-head attention:

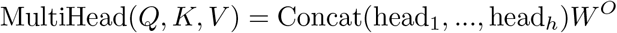

where Q, K, V are query, key, and values. *W^O^* is the out projection of dimension (number of heads h times value dimension *d_v_* by model dimension *d_m_*). In our implementation *d_v_* and *d_v_* are set to 128. *head_i_* is a single attention head and is calculated by:

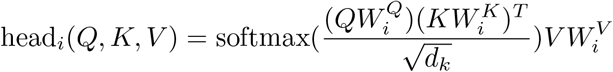

where *W^Q^*, *W^K^*, *W^V^* are projection weights for query, key and value. *d_k_* is the embedding dimension of key, also implemented as 128. During forward propagation, we extract attention weights for head *i* which is defined as the alignment between query and key:

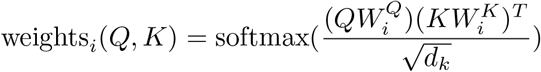

The attention score can be calculated by averaging attention weights across different heads:

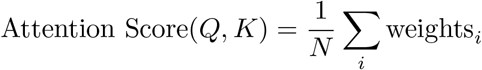

where N = 8 because each layer has eight attention heads. Since the transformer module consists of eight attention layers, for each prediction, we obtained a set of eight attention scores. The attention score is visualized with the BertViz package^72^.

### Impact score

The impact score in the screening experiment is defined as the pixel-wise mean absolute difference between two predictions. Formally, given that we have a prediction S, a 2D contact matrix from the original input and S’ from the input perturbed at location x, and let s{i, j} be the individual pixel in S at position i and j, the impact score of location x is defined as:

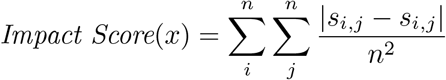

### *In silico* genetic screening

Typical ChIP-seq profiles have peak widths ranging from a few hundred base pairs to 1Kb. To capture fine-regulation elements, we performed genome-wide *in silico* genetic screening at 1Kb resolution. The screening starts from individual chromosomes with a window size of 2Mb, denoted as (i, i + 2097152). Inside this window, a 1Kb perturbation region centered at the 2Mb window, or location (i + 2097152 / 2 - 500, i + 2097152 / 2 + 500), was deleted followed by C.Origami prediction. After deleting the 1Kb segment, we appended a 1Kb empty input at the end to keep a complete 2Mb window size for C.Origami prediction. For each window, the original input and perturbed input were predicted by C.Origami, resulting to two outputs, S_i and S_i’, which were collected for downstream impact score calculation. Once the output acquisition was completed for the window at (i, i + 2097152), the screening moves to a downstream overlapping window that has 1Kb offset from the current window with range (i + 1000, i + 2097152 + 1000). The mean absolute average of difference between the original and perturbed output S_i and S_i’ were computed and attributed to the perturbation region (i + 2097152 / 2 - 500, i + 2097152 / 2 + 500). Since the *in silico* screening offset is equal to the length of perturbation size, this procedure produces a continuous impact score that covers all genomic regions with a resolution of 1Kb.

It is worth noting that screening at 1Kb resolution could be computationally intensive. For instance, screening on chromosome 8, a medium-size chromosome which has a length of 146Mb, requires the model to make 146Mb / 1Kb * 2 predictions = 292,000 separate predictions. In our optimized framework that predicts 600 windows per minute, and screening chromosome 8 takes 8 hours. To reduce computational load, we randomly sampled 10 chromosomes (chr 5, 7, 8, 11, 12, 14, 15, 19, 20, 22) to represent the whole genome and performed 1Kb-resolution screening on the selected chromosomes.

In order to obtain the most impactful elements from the screening result, we designed a custom peak calling algorithm. We defined the peak score *p* of a locus as the difference between maximum and minimum signal within the range of 3 bins including the locus. We then selected the top 1% of the total screened regions as a cutoff for impactful elements based on the peak score.

To annotate the *in silico* genetic screen-identified impactful *cis*-elements, we compiled a set of genomic annotations including TAD boundary regions, enhancers, promoters, intragenic regions and intergenic regions. The boundary region was generated by calling TAD boundaries at 10Kb resolution with HiC-bench^64^, using its TopDom module and connecting adjacent TADs. To increase robustness of TAD boundary calling, we expanded the boundary width to 5 bins, or 50Kb. The promoter region was defined as 5.5Kb fragments, spanning 5Kb upstream and 500bp downstream of gene transcription start site. Enhancers were defined as by the H3K4me1 modification, which marks both active and inactive enhancers^73^. The H3K4me1 ChIP-seq peaks for IMR-90 was downloaded from ENOCDE with accession number: ENCFF611UWF. To increase robustness, we expanded peaks to have at least 1Kb width.

### *In silico* genetic screen at 2Mb windows

We conducted *in silico* genetic screen at a fixed 2Mb window without centering the deletion element. We systematically removing segments of 8,192 bp, or 1 bin, from model inputs. To scan through the entire 2Mb region, we performed 256 deletion experiments at each bin and calculated the prediction difference map before and after deletion. Deletion reduces the input length from 2,097,152 bp to 2,088,960 bp. To maintain input shape, we appended 8,192 bp of empty input features to facilitate subsequent prediction.

### CRISPR screening for chromatin remodeling genes in T-ALL cell lines

Pooled CRISPR screening across 313 chromatin remodeling genes in CUTLL1 and Jurkat cells were carried out in parallel with our previous studies for pooled screening of RNA binding protein in T-ALL cells^74^. Briefly, for each chromatin remodeling gene, we designed on average 6-8 sgRNA, for a total of ∼2,500 sgRNAs. The gRNA sequences were synthesized from Twist Bioscience, and cloned into a lentivirus-based sgRNA vector tagged with GFP (Addgene plasmid no. 65656). Cas9-expressing T-ALL cell lines were transduced with sgRNA library virus at a low MOI (∼0.3), followed by infection efficiency assessment through GFP percentage on Day 4 post-transduction. Remaining cells were placed back into culture until 20 days post-transduction.

Cell proliferation was measured by comparing the sgRNA frequencies between Day 4 and Day 20 cells. Genomic DNA was harvested on Day 4 and Day 20 cells using Qiagen DNA Purification kit based on the manufacturer’s protocol. The gRNA frequencies in the genomic DNA were amplified and quantified following our previous procedure^75^. For pooled CRISPR screening analysis, samples of each time-point were normalized as sgRNA read count / total read count x 100,000. Subsequently, normalized reads were then used to calculate log2 fold change as (normalized read count Day 4 / normalized read count Day 20) for each gRNA. The fold changes between Day 4 and Day 20 for each gene were averaged from all CRISPR gRNA targets. P values were calculated via a two-sided t-test comparing the fold changes of all gRNA targets of the same gene to fold change of 1.

### Virtual 4C

HiC-Bench “virtual4C” pipeline^64^ was used to compute the interactions of each selected viewpoint in a roll-window fashion. We summed the valid read pairs in a 5 kb area centered at 100 bp bins that covered the area of +-2.5 Mb from the viewpoint (50k bins per viewpoint). The interactions were normalized by the total number of valid pairs of the sample.

### *Trans*-acting regulator identification in T-ALL cell lines

Different cell types have a unique set of impactful *cis*-elements, which constitutes the cell type-specific chromatin interaction map. To connect the differential patterns of *cis*-elements with *trans*-acting regulators, we compared selected the cell type-specific impactful regions by a custom peak calling method, followed by a transcription factor enrichment test for identifying potential *trans-*acting regulators. We used the transcription factor database from ReMap2022^53^. To reduce low quality signals from the ReMap database, we filtered out transcription factors profiles that have less than 7000 hits, or profiles that only have one experiment. Together, we collected 612 transcription factor binding profiles for downstream analysis. We used Fisher’s exact test to evaluate the overlap between impactful *cis*-elements from *in silico* genetic screening with each transcription factor from the database. The test was conducted using the LOLA package (Locus Overlap Analysis)^75^. For common transcription factors with hit counts larger than 20K, we down sampled profiles to 20K. We calculated the q value with FDR correction based on the 612 TF profiles tested and used odds ratio as the main metric to determine enrichment of each factor in impactful *cis*-elements.

To compare the contributing *trans-*acting regulator profiles between different cell types, we first normalized the odds ratio within each cell type. We performed k-means clustering of transcription factors based on their normalized odds ratio in CUTLL1, Jurkat and T cells. K-means clustering was performed with standard Euclidean distance with 6 centroids. The clusters were further grouped and visualized using a heatmap.

### Intra-TAD activity analysis

Iteratively corrected matrices were re-normalized by dividing each bin value by the sum of all the values in the same distance bin in the same chromosome (distance-normalization). All the TADs identified in the control sample were used as the reference TADs to compute the intra-TAD activity changes. The set of reference TADs between the two samples S1 (control) and S2 (treatment) were denoted as set T. A paired two-sided t-test was performed on each single interaction bin within each reference TAD between the two samples. We also calculated the difference between the average scores of all interaction intensities within such TADs and the TAD interaction log fold-change. Finally, a multiple testing correction by calculating the false-discovery rate on the total number of TAD pairs tested. The TAD interaction change for each <u>t</u> in T is define as follow:

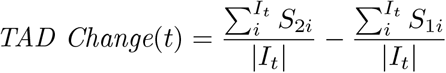

We classified the reference TADs in terms of Loss, Gain or Stable intra-TAD changes by using the following thresholds: FDR < 0.01, absolute TAD interaction log fold change > 0.25, and absolute TAD interaction change > 0.1.

## Data availability

Most of the Hi-C, CTCF ChIP-seq, and ATAC-seq datasets used in the study were public data from ENCODE portal and/or NCBI GEO database, with the accession codes listed in the corresponding Methods section. The generated data (CUTLL1 ATAC-seq) is uploaded to GEO with accession number GSE216430.

## Code availability

The code for C.Origami is available at https://github.com/tanjimin/C.Origami.

## Supplementary Figures

**Supplementary Figure 1:**
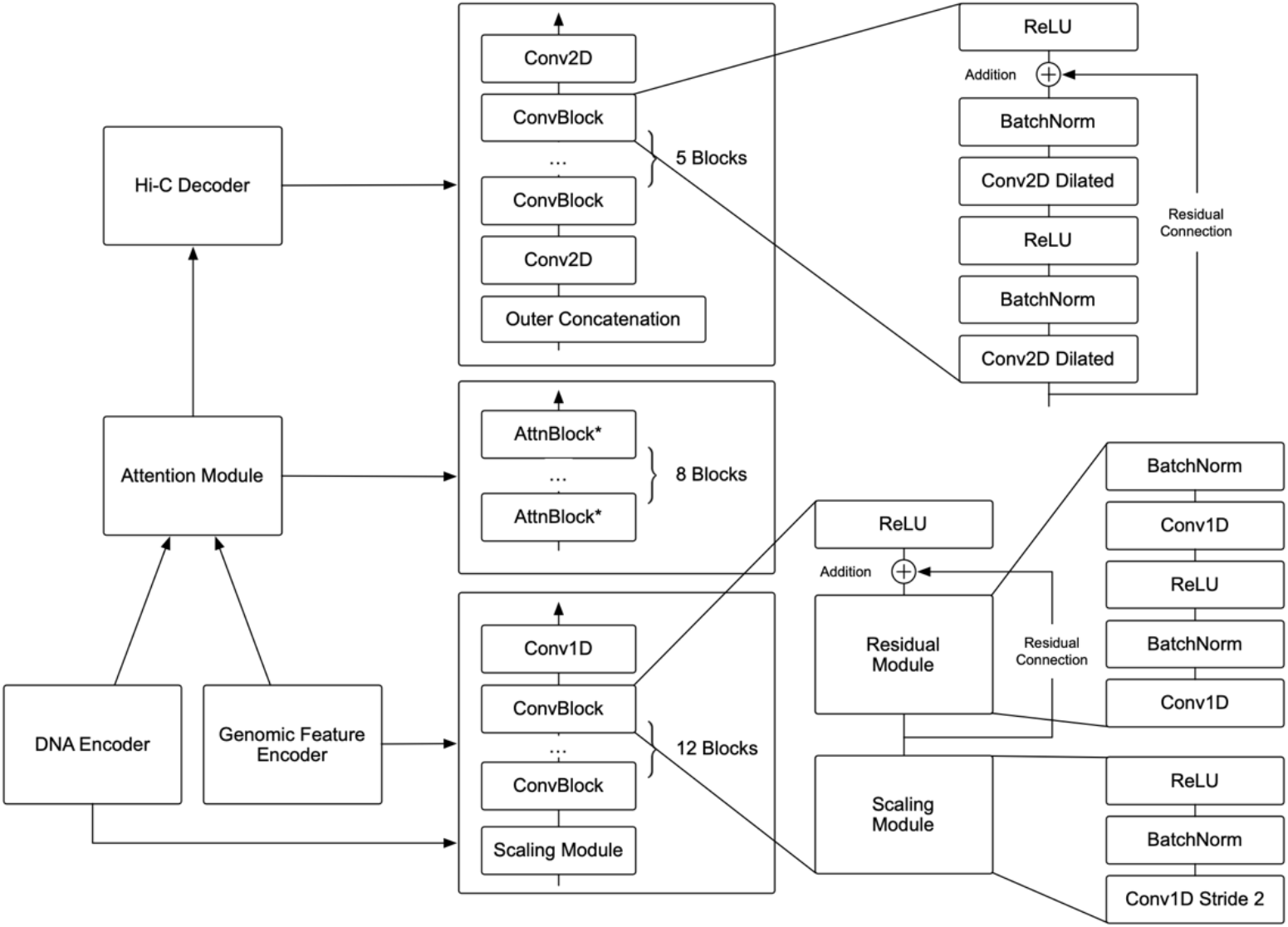
C.Origami model structure and module components. A schematic of C.Origami model architecture. The DNA encoder and Genomic Feature encoder have similar architectures and only different in input channels where DNA encoder has 5 channels and feature encoder has 2 channels. We built the encoder with 12 convolution blocks. Each block consists of a scaling module and a residual module. The scaling module downscales input features by a factor of two with a stride-2 1D convolution layer. The residual module promotes information propagation in very deep networks^76^. The number of modules was carefully chosen so that the 2,097,152 input are scaled down to 256 bins at the end of the encoder. To enhance interactions within the 2Mb window, we used an attention module consisting of eight attention blocks. Each position of the output is concatenated with every other position to form a 2D matrix, resembling a vector outer product process. To refine the final prediction, we used a 5-layer dilated 2D convolutional network as decoder. We deliberately chose the dilation parameters to ensure that every position at the last layer has a receptive field covering the input range.

**Supplementary Figure 2:**
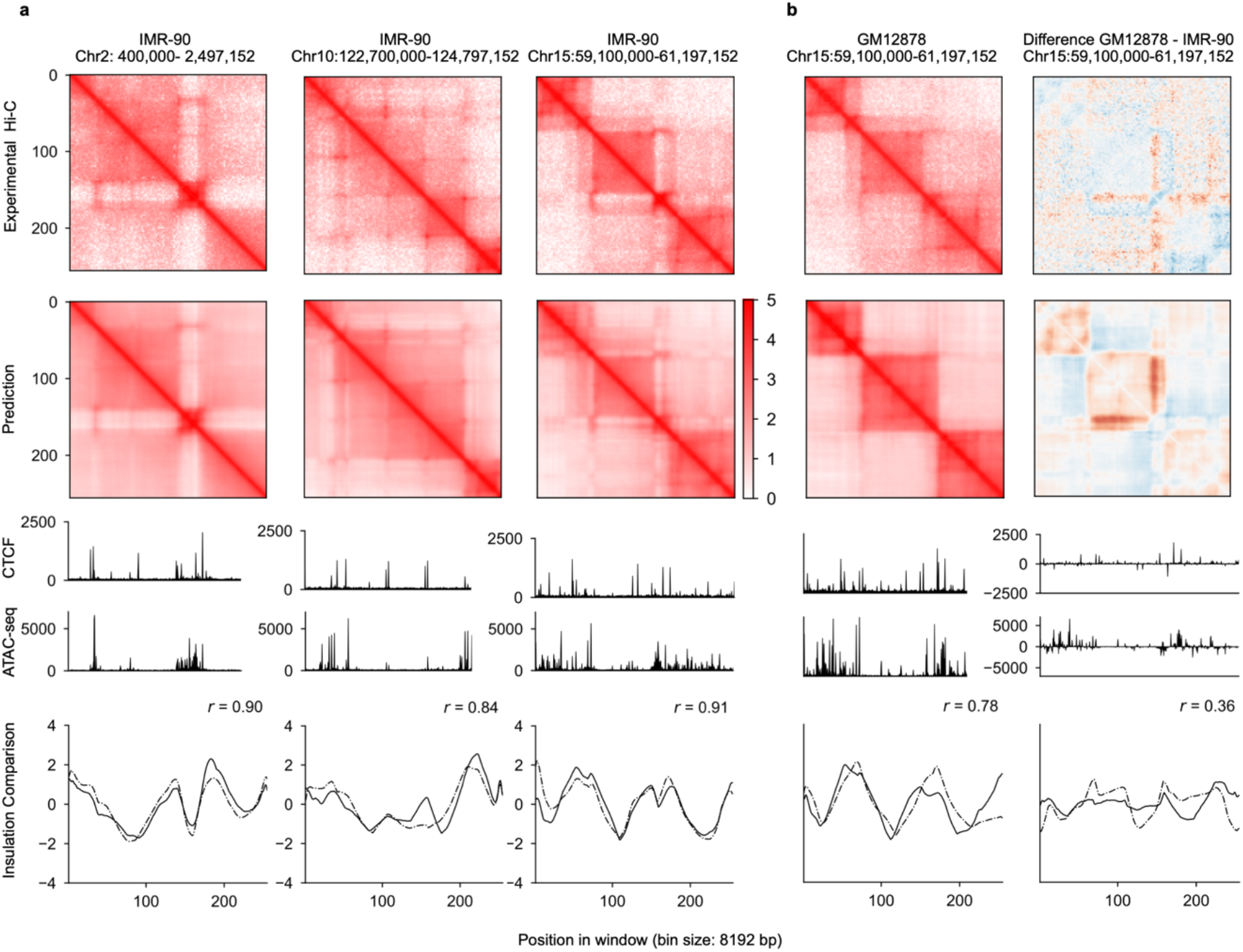
Performance of C.Origami trained with DNA sequence and CTCF ChIP-seq. **a**, Prediction from a model trained with DNA sequence and CTCF ChIP-seq. The plots were organized the same way as Fig. 2. **b**, *De novo* predicting chromatin organization of the chromosome 15 locus in GM12878 using the model trained with DNA sequence and CTCF binding profiles. The difference between IMR-90 and GM12878 is presented on the right. While C.Origami trained with DNA sequence and CTCF profile achieved good performance in validation and test set in IMR-90 (**a**), it missed predicting some fine-scale chromatin structures in GM12878.

**Supplementary Figure 3:**
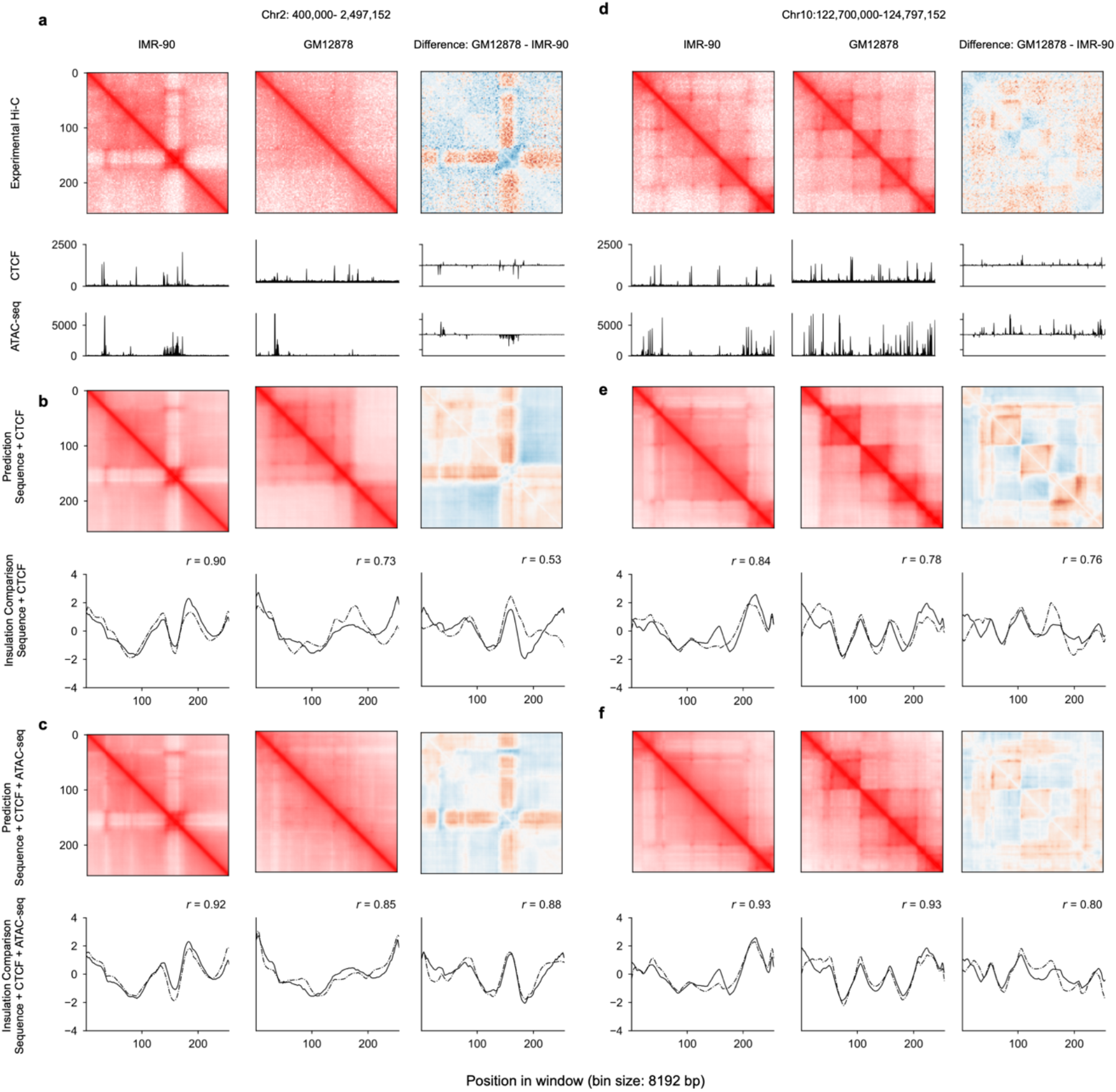
C.Origami trained with DNA sequence, CTCF binding, and chromatin accessibility profiles performed optimally. **a**, Experimental Hi-C matrices, and genomic profiles of IMR-90 and GM12878 cells at chr2:400,000-2,497,152. The difference between the two cell lines were presented on the right. **b-c**, Cell type-specific prediction of the chromatin organization at the same locus using C.Origami (**b**) or model trained with DNA sequence and CTCF binding (**c**). **d-e**, Same as **a-c** at a difference locus, chr10:122,700,000-122,797,152.

**Supplementary Figure 4:**
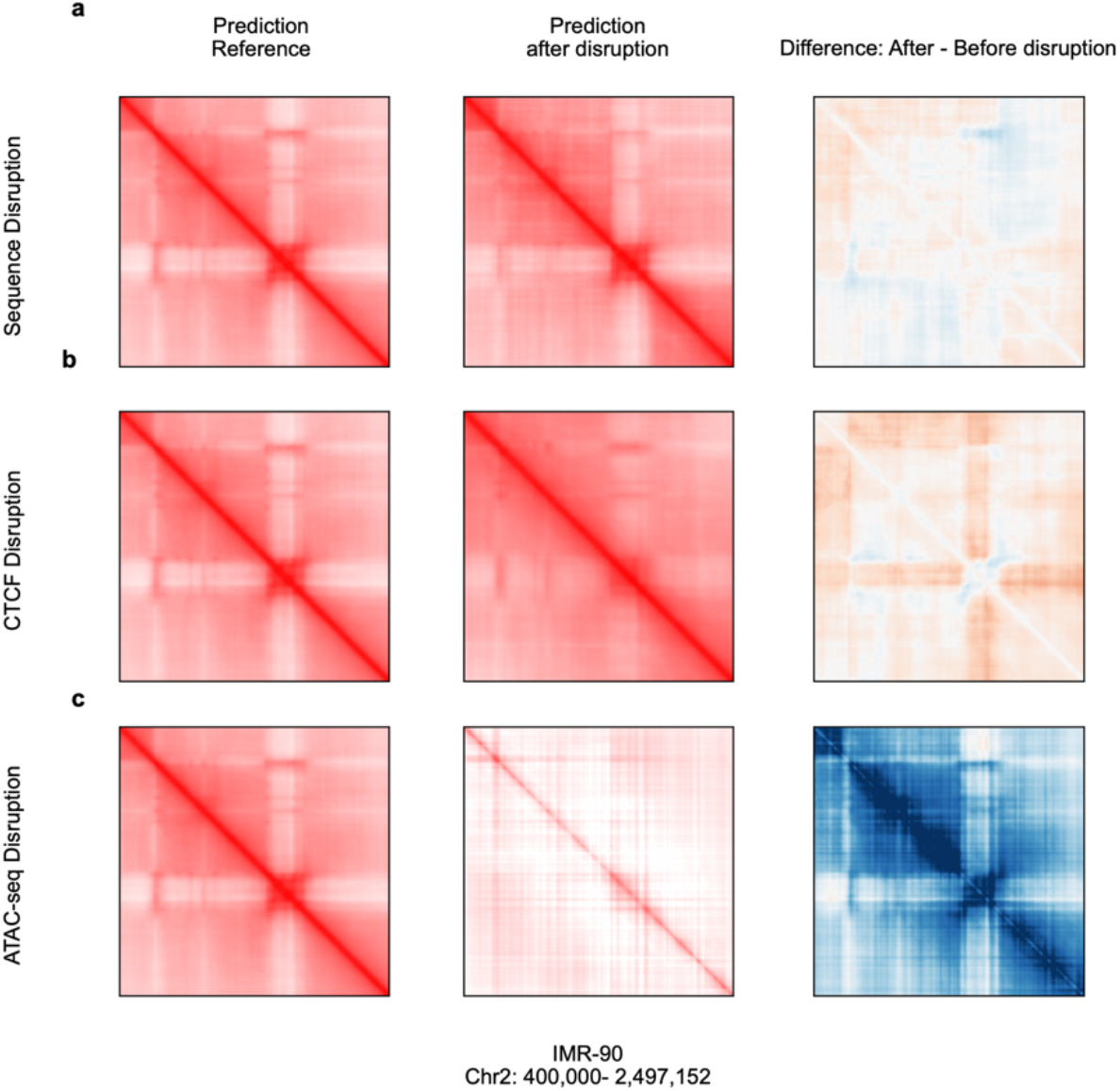
Ablation study on different input features. Using C.Origami trained with DNA sequence, CTCF binding, and chromatin accessibility profiles, the experiments were performed by random shuffling DNA sequences at base pair level (**a**), random shuffling CTCF signal (**b**), and random shuffling ATAC-seq signal (**c**). From left to right, reference prediction with all inputs (left), prediction with sequence shuffled (middle), difference between perturbed prediction and reference prediction (right).

**Supplementary Figure 5:**
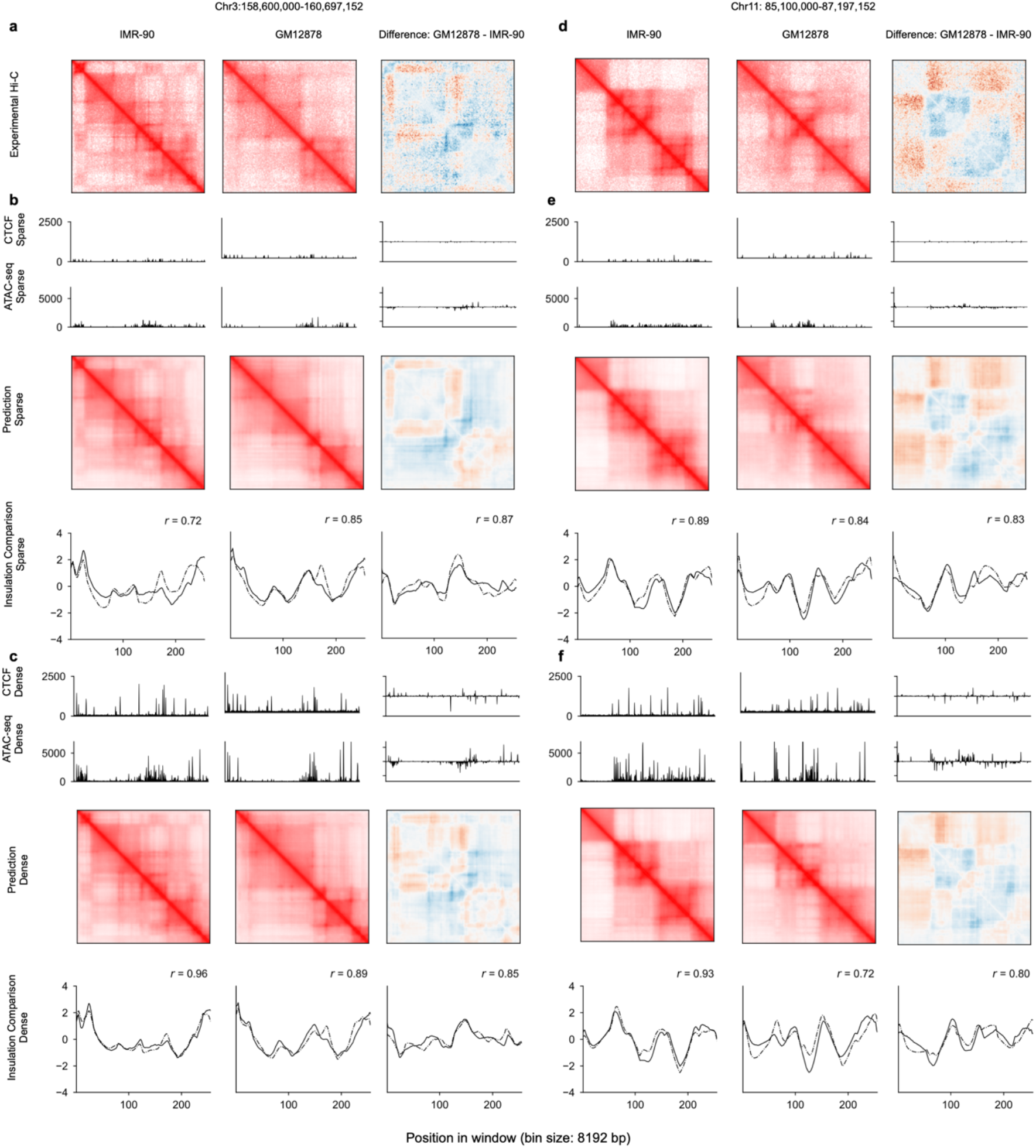
Performance comparison of C.Origami models trained with sparse information and dense information. **a**, Experimental Hi-C matrices of IMR-90 and GM12878 cells at chr3: 158,600,000-160,697,152. The difference between the two cell lines were presented on the right. **b-c**, Cell type-specific prediction of the chromatin organization at the same locus using C.Origami models trains with sparse genomic information (**b**) or dense genomic information (**c**). For each set of plots in **b** and **c**, the input CTCF ChIP-seq and ATAC-seq profiles were aligned with the predicted Hi-C matrices and the insulation score results. **d-f**, Same as **a-c** at a difference locus, chr10: 85,100,000-87,197,152.

**Supplementary Figure 6:**
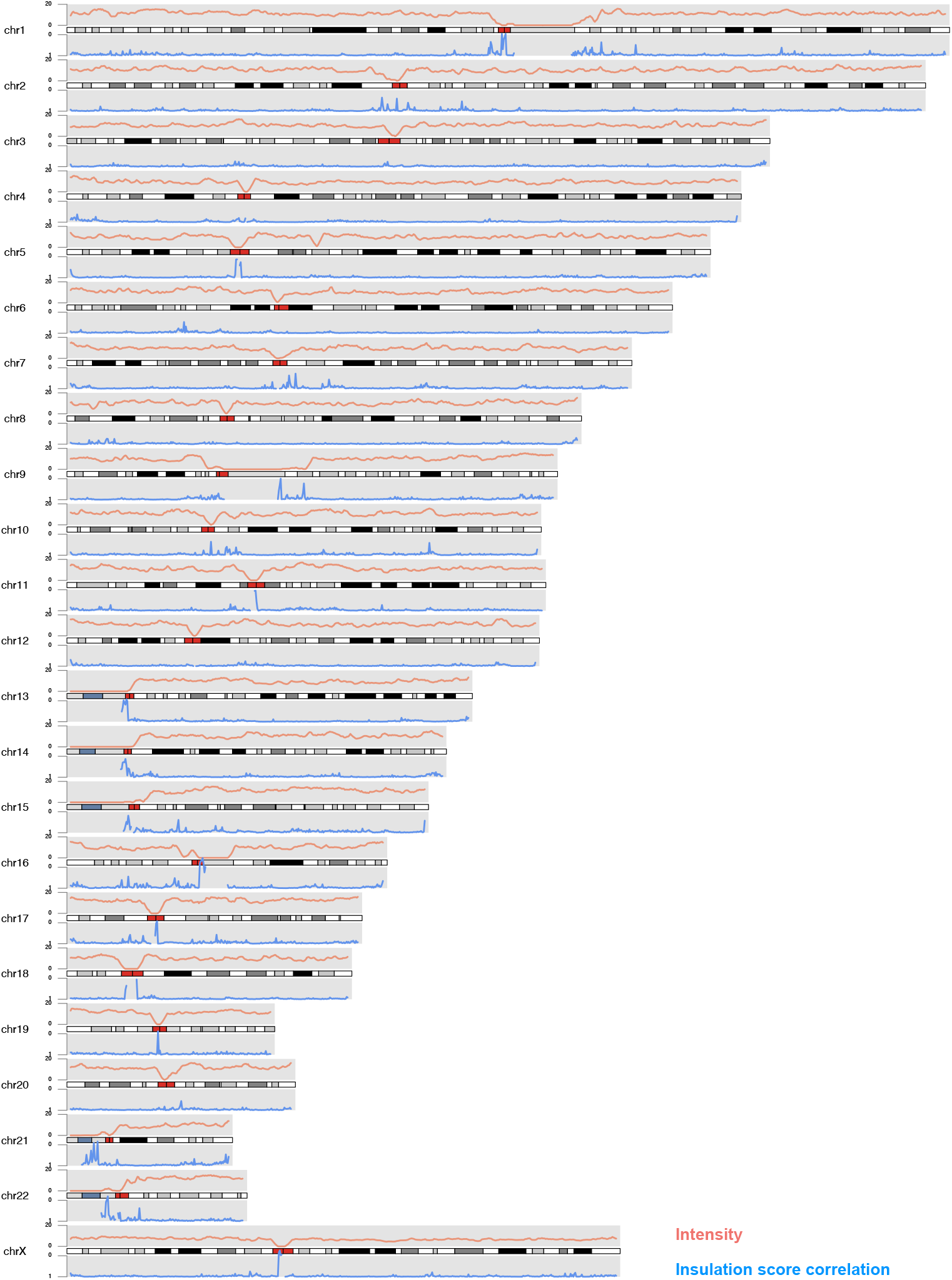
Chromosome karyotype visualization along with chromosome-wide Hi-C intensity and correlation of insulation scores. The results were visualized using karyoploteR^77^. Chromosome 1 to chromosome X were plotted to visualize the Pearson correlation coefficients of insulation scores calculated from prediction and that from experimental Hi-C. Average intensity of 2Mb windows were plotted in red. Centromere regions were denoted with red segments on the genome. The few data points with low intensity are regions corresponding to unmappable or repeat sequences such as centromeres and telomeres.

**Supplementary Figure 7:**
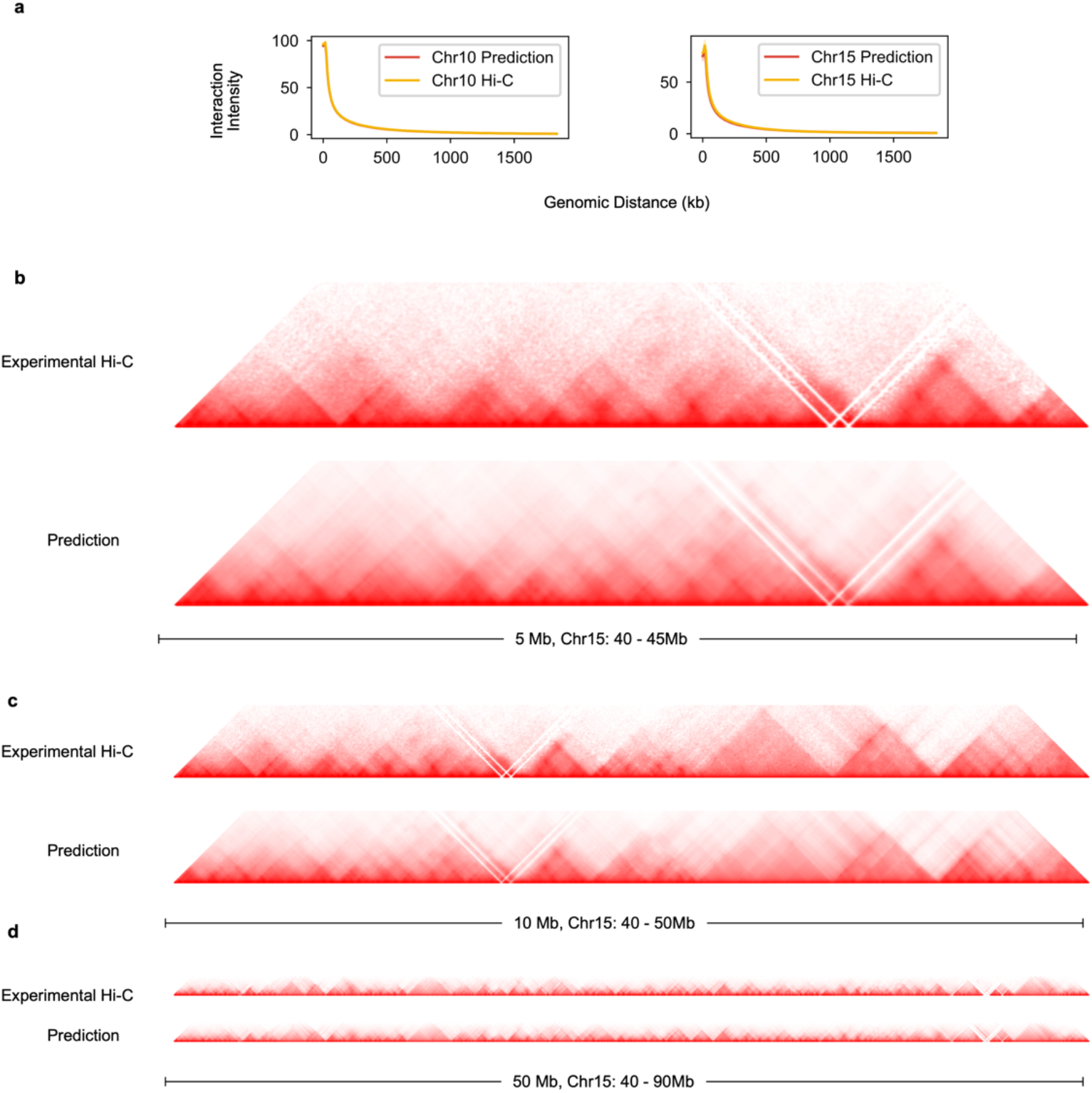
C.Origami-predicted 2Mb Hi-C maps can be fused into larger interaction maps. **a,** Interaction intensity distribution of prediction and experimental Hi-C on validation (chromosome 10) and test chromosomes (chromosome 15). **b-d,** The predicted 2Mb Hi-C maps were fused to 5Mb (**b**), 10Mb (**c**), and 50Mb (**d**) on chromosome 15, all with the same starting site at 40 Mb.

**Supplementary Figure 8:**
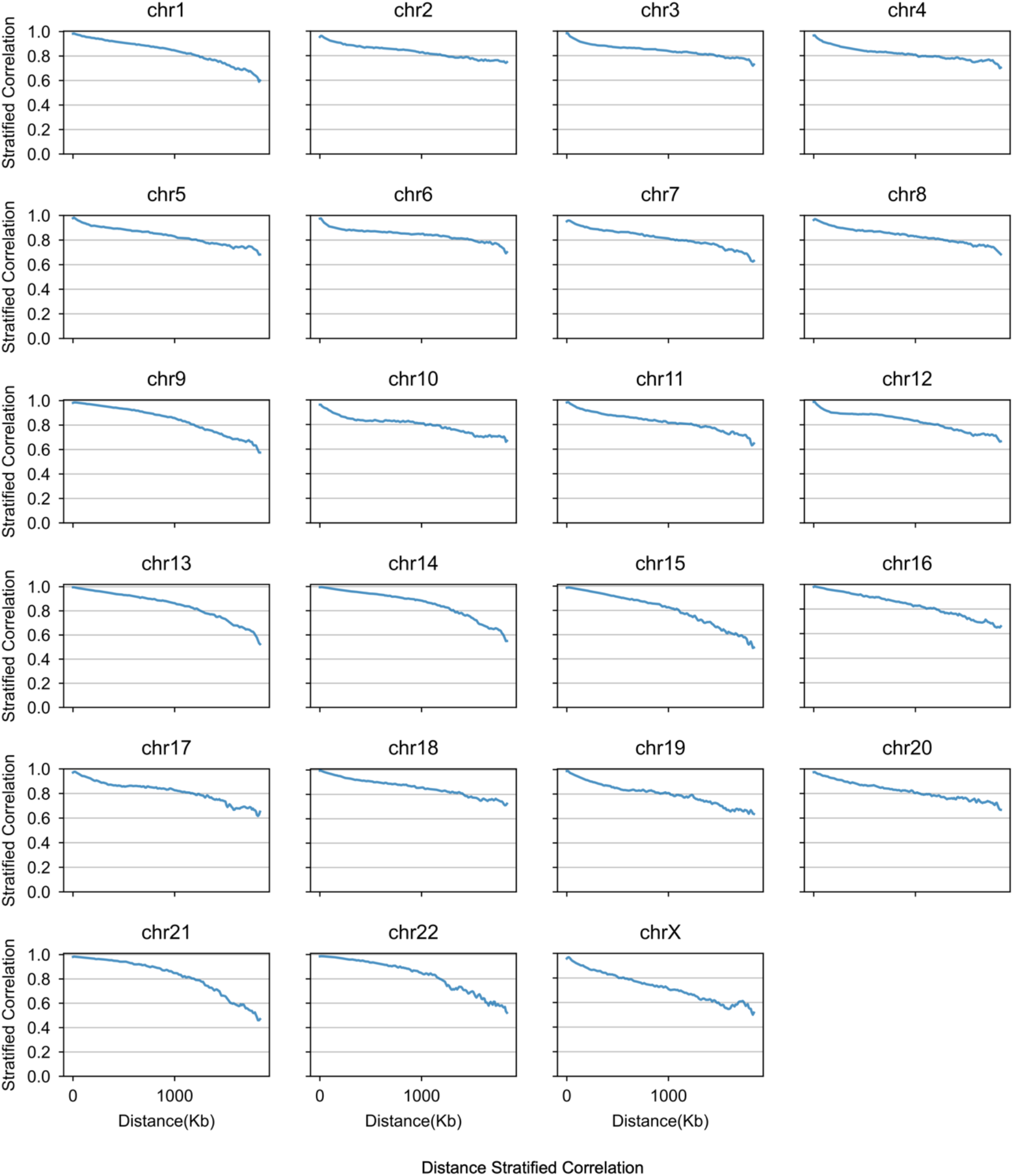
Chromosome-level distance-stratified intensity correlation. **a**, Interaction intensity distribution of prediction and experimental Hi-C on validation (chromosome 10) and test chromosome (chromosome 15). Chromosome-level distance-stratified correlation between prediction and experimental Hi-C were calculated on each chromosome of IMR-90 cells.

**Supplementary Figure 9:**
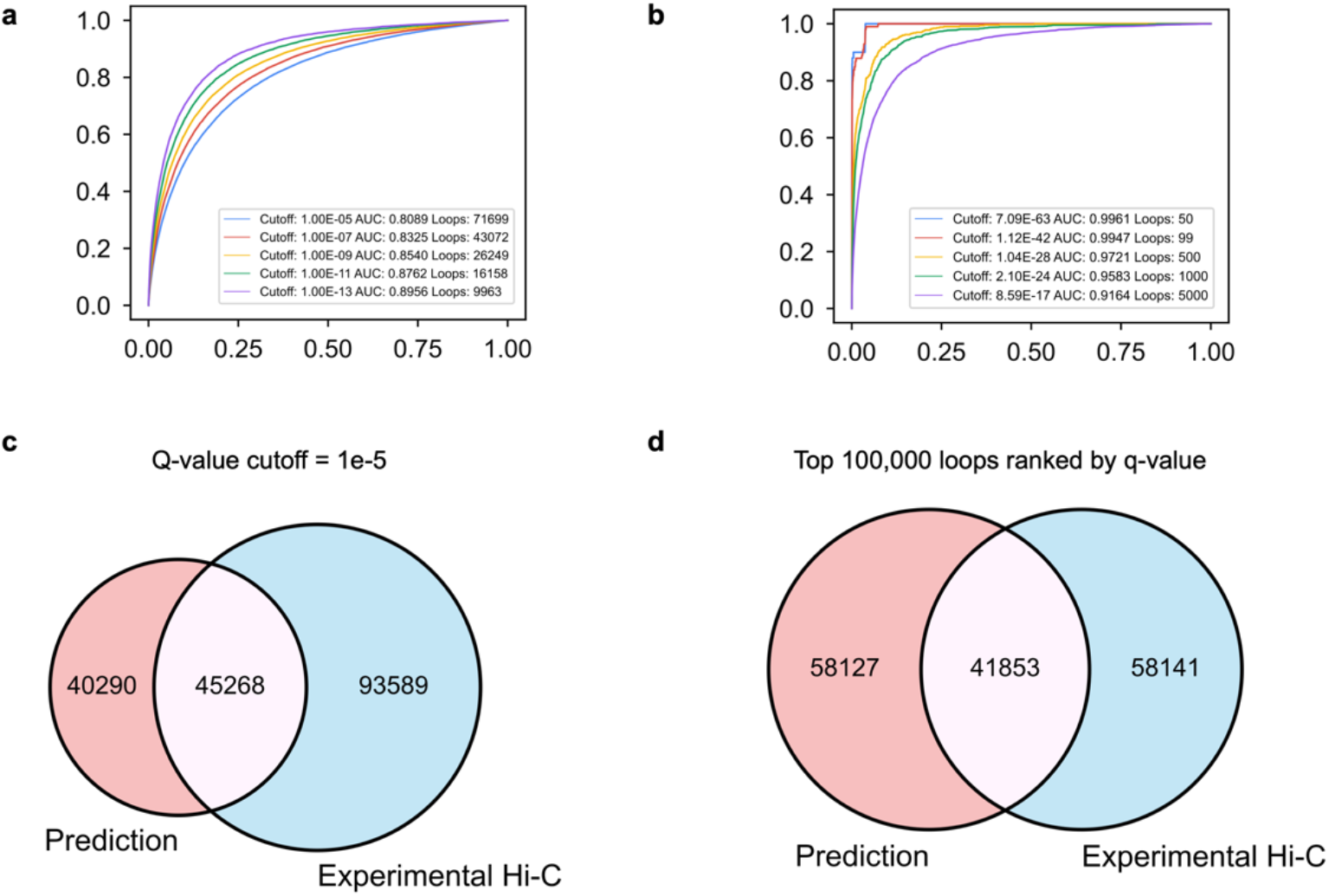
Evaluating C.Origami’s performance on detecting significant chromatin loops in IMR-90 cells. **a**, ROC curves of significant chromatin loops called in experimental Hi-C and prediction. Significant chromatin loop referring to global background were called at different q-value ranging from 1e-5 to 1e-13 from predicted Hi-C matrices. Q-value of experimental Hi-C was ranked against predicted loops to calculate AUROC. Each curve represents an ROC curve comparing experimental Hi-C q-value to predicted loops with specific cutoffs. **b**, ROC curves of top 50 to top 5000 loops with corresponding q-value cutoffs. AUROC under each criterion is indicated in legends of **a** and **b**. **c-d**, Venn diagram of chromatin loop overlapping between experiment and prediction with q-value cutoff of at 1e-5 (**c**) or between the top 100,000 loops (**d**). All loop calling was carried out with global background as reference to increase sensitivity to all significant chromatin interactions.

**Supplementary Figure 10:**
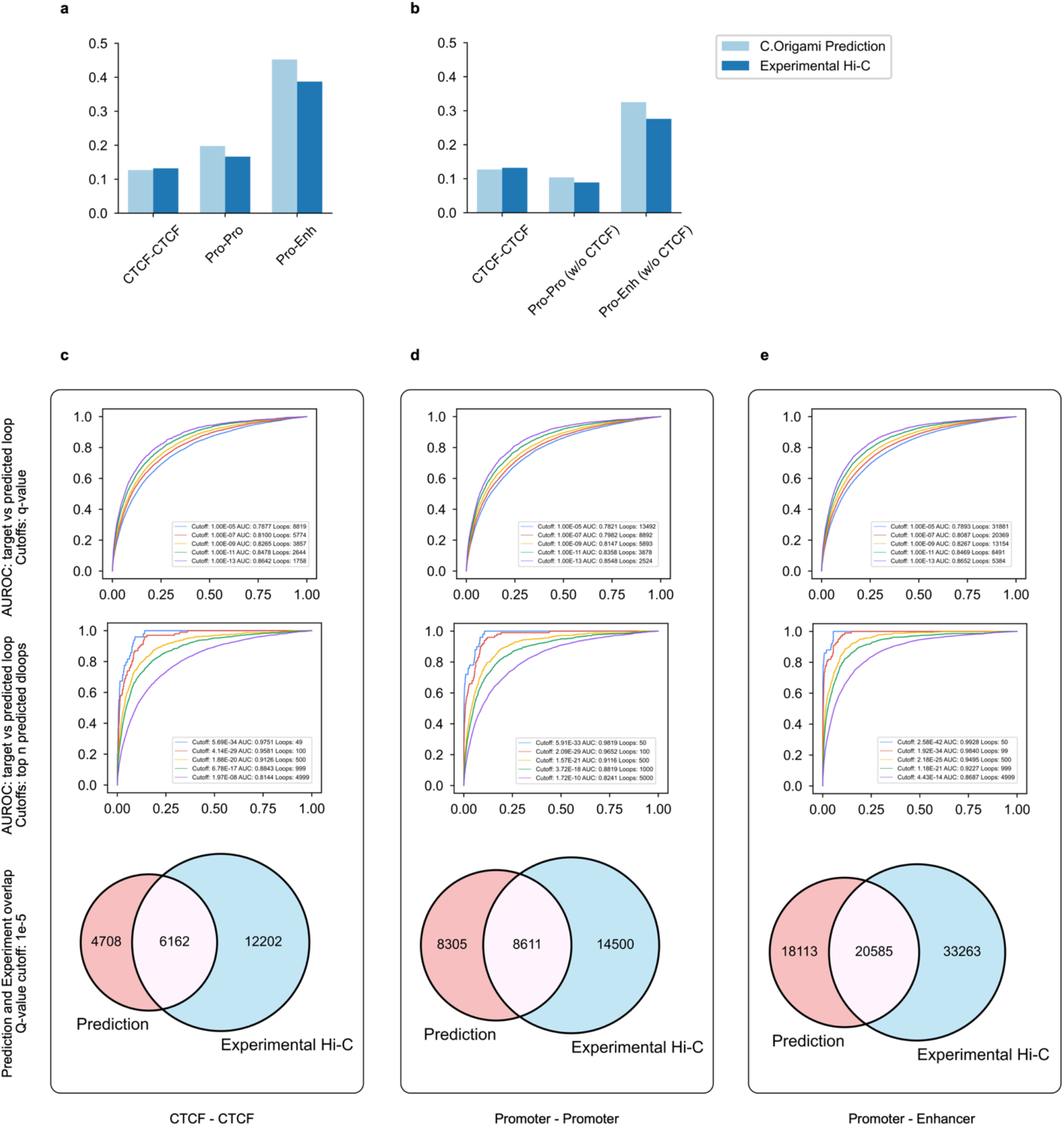
Performance of detecting loop interactions under different chromatin backgrounds. **a-b**, Percentages of loop counts in three different categories, including CTCF-CTCF loop, promoter-promoter loop, and promoter-enhancer loop. Significant chromatin loop referring to global background were called at different q-value in IMR-90 cells and then categorized according to their anchor content. Within each panel, AUROC between loops from experiment and prediction was calculated with q-value cutoffs ranging from 1e-5 to 1e-13, similar to the previous loop analysis. Category counts were divided by the total number of loops called. **c-e,** ROC curves and the Venn diagrams of the significant chromatin loops called in experimental Hi-C and prediction categorized by anchor content: CTCF-CTCF loop (**c**), promoter-promoter loop (**d**), and promoter-enhancer loop (**e**). AUROC from top 50 to top 5000 loops were also plotted.

**Supplementary Figure 11:**
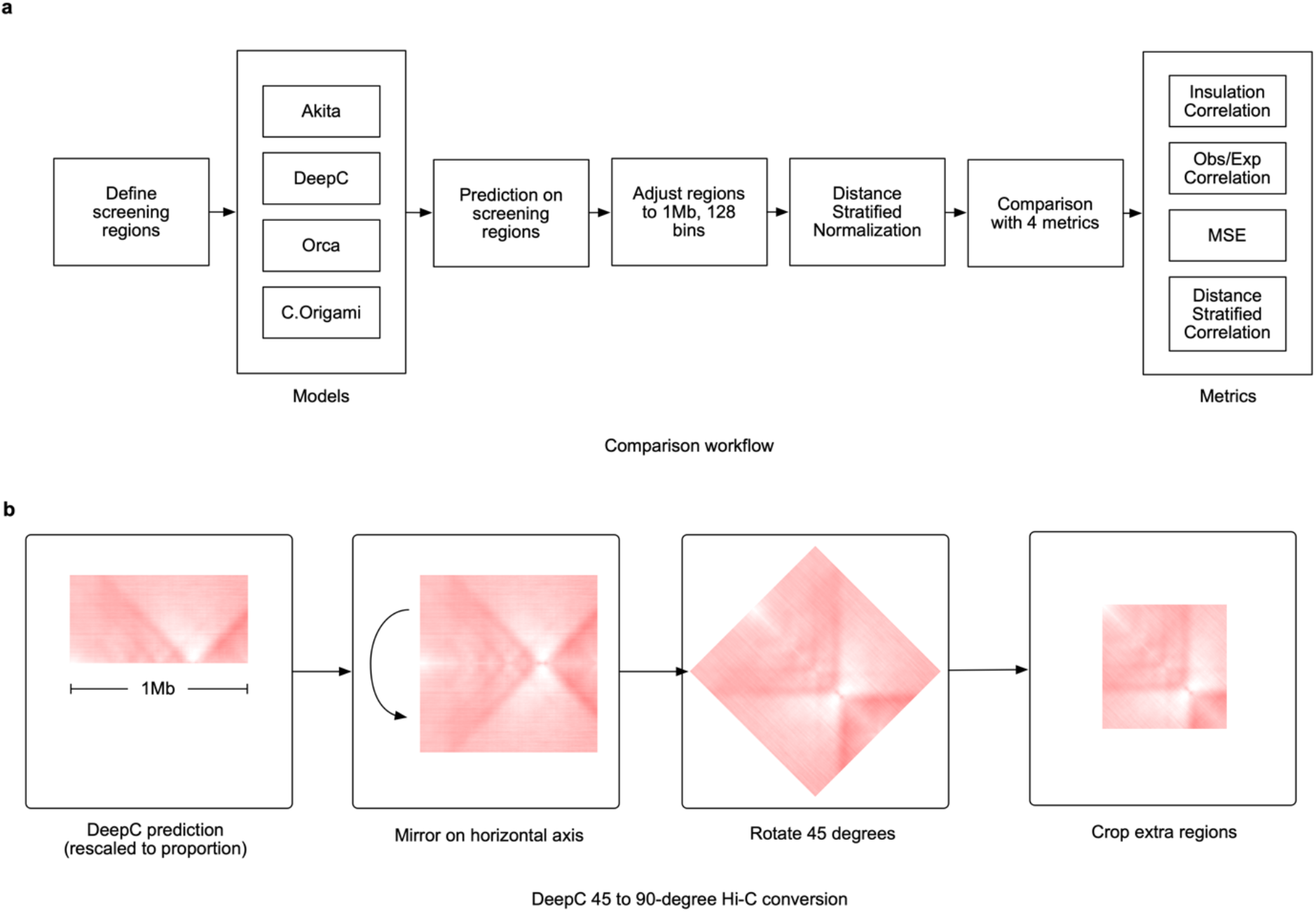
Workflow of comparing performance of models predicting 3D chromatin organization. **a**, Workflow of the comparison procedures to standardize and evaluate the predictions from Akita, DeepC, Orca, and C.Origami. **b**, Post-processing of DeepC prediction results. DeepC method by default produces a 45 degree Hi-C map, thus requiring mirroring, rotation and cropping steps to make the results comparable to Hi-C targets.

**Supplementary Figure 12:**
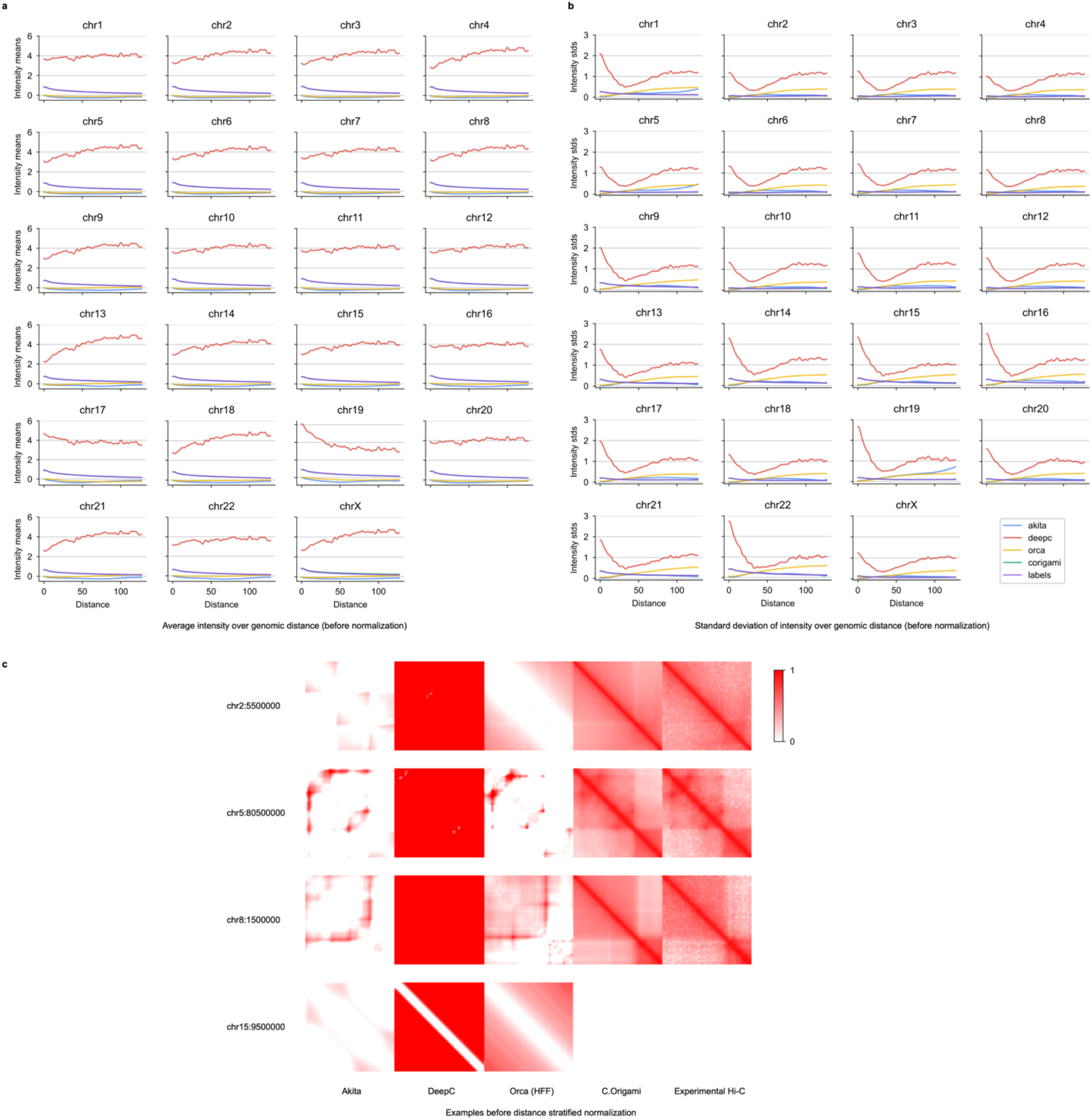
Distance-stratified statistics of raw predictions results from the four models in comparison. **a-b**, Distance-stratified mean intensity (**a**) and standard deviation (**b**) of predicted Hi-C results from the four models. The horizontal axis denotes the rescaled 128 bins representing a 1Mb region. DeepC has a different distribution of intensities compared to the rest of the models. The abnormality could be a result of its custom percentile normalization on the training target. **c.** Raw prediction results from four models together with experimental Hi-C. Intensity values was set to be from 0 to 1 according to experimental Hi-C data.

**Supplementary Figure 13:**
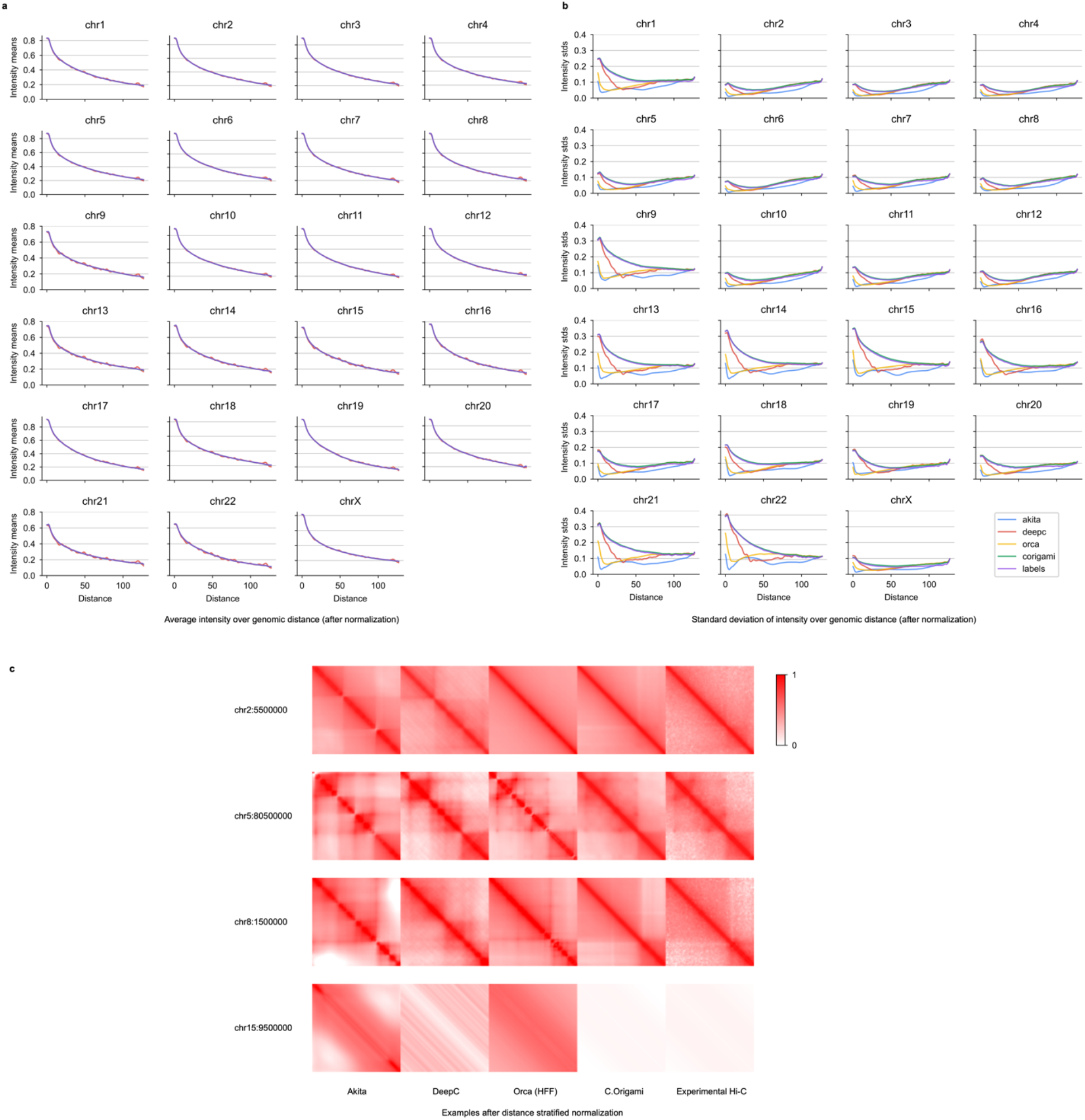
Distance-stratified statistics of prediction results after standardization. **a-b**, Distance-stratified mean intensity (**a**) and standard deviation (**b**) of predicted Hi-C results from the four models after distance-stratified normalization. After normalization, the differences between all model predictions are comparable to experimental Hi-C. **c.** Normalized prediction results from four models together with experimental Hi-C. Intensity values was set to be from 0 to 1 according to experimental Hi-C data. Presented loci are from the same regions as in Supplementary Figure 12. In comparison, normalized predictions are more comparable in between and closer to the experimental Hi-C.

**Supplementary Figure 14:**
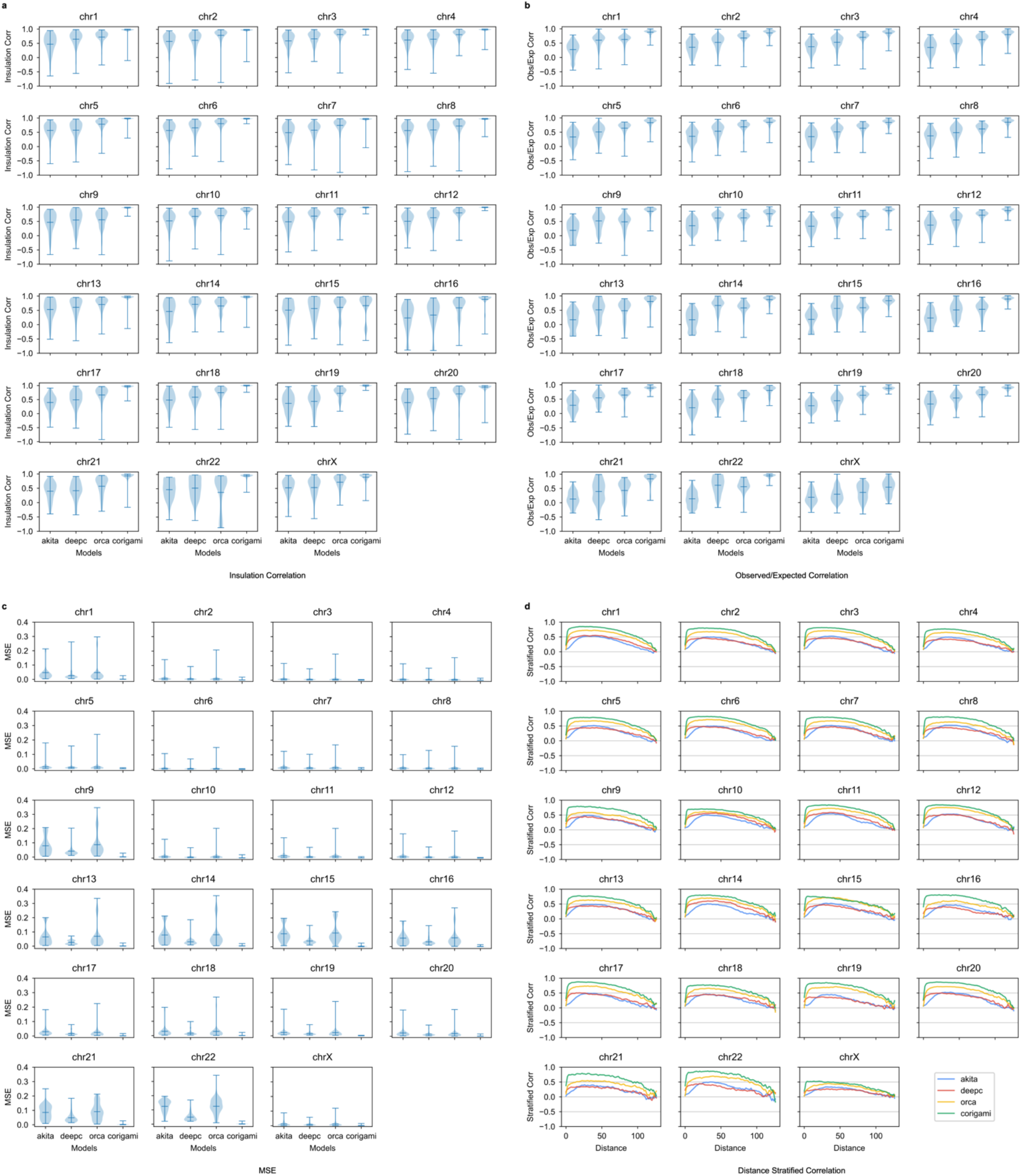
Genome-wide comparison of model performance in IMR-90 cells. For predictions from each model (Akita, DeepC, Orca and C.Origami), we measured insulation score correlation (**a**), observed vs expected Hi-C matrices correlation (**b**), mean squared error (MSE, **c**), and distance-stratified correlation (**d**).

**Supplementary Figure 15:**
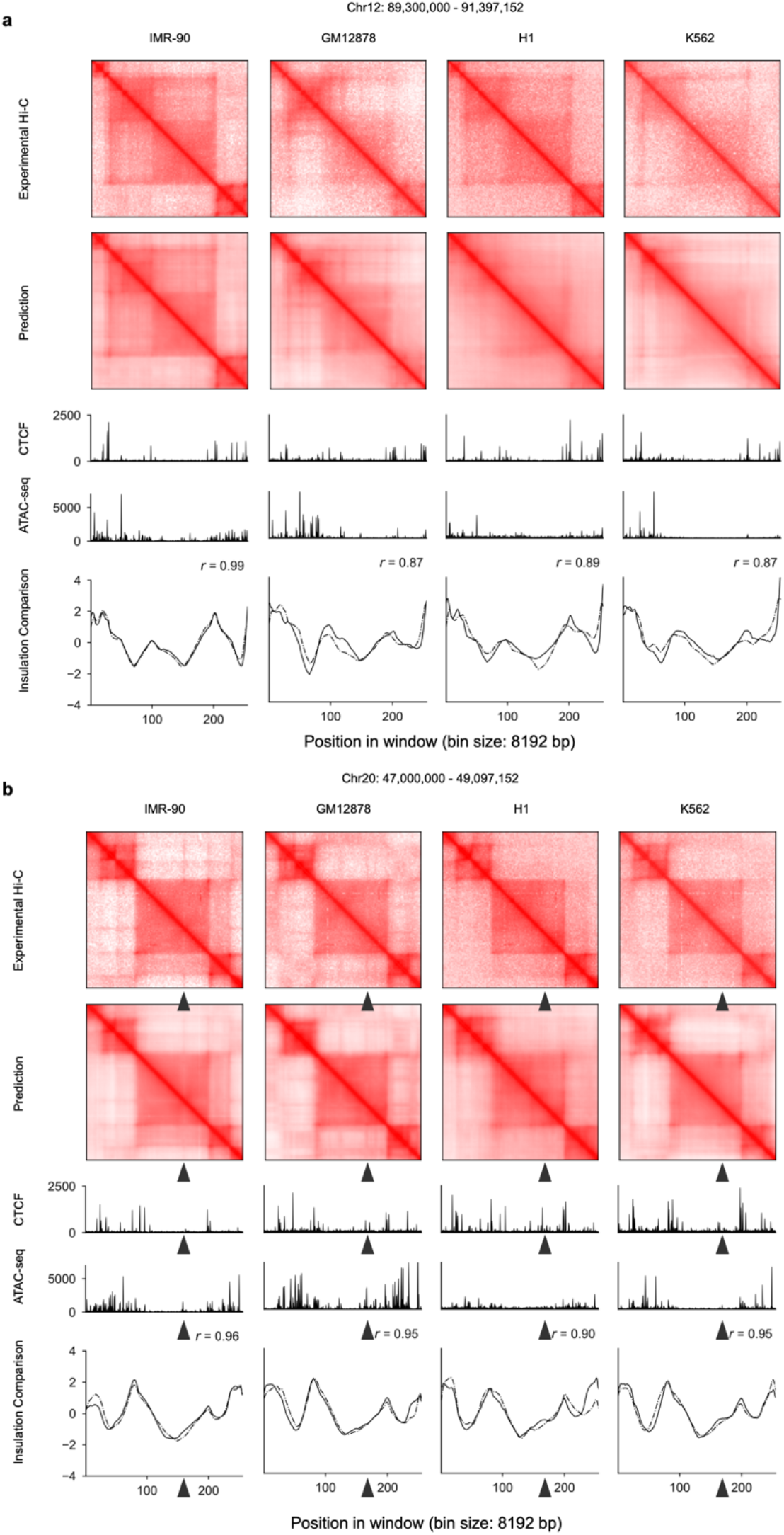
C.Origami predicts chromatin organizations across multiple cell types. Two representative loci were separately presented across IMR-90, GM12878, H1-hESCs, and K562 in **a** and **b**. From top to bottom, each panel included experimental Hi-C matrix, predicted Hi-C matrix, CTCF and ATAC-seq signals, and insulation scores calculated from experimental and predicted Hi-C data.

**Supplementary Figure 16:**
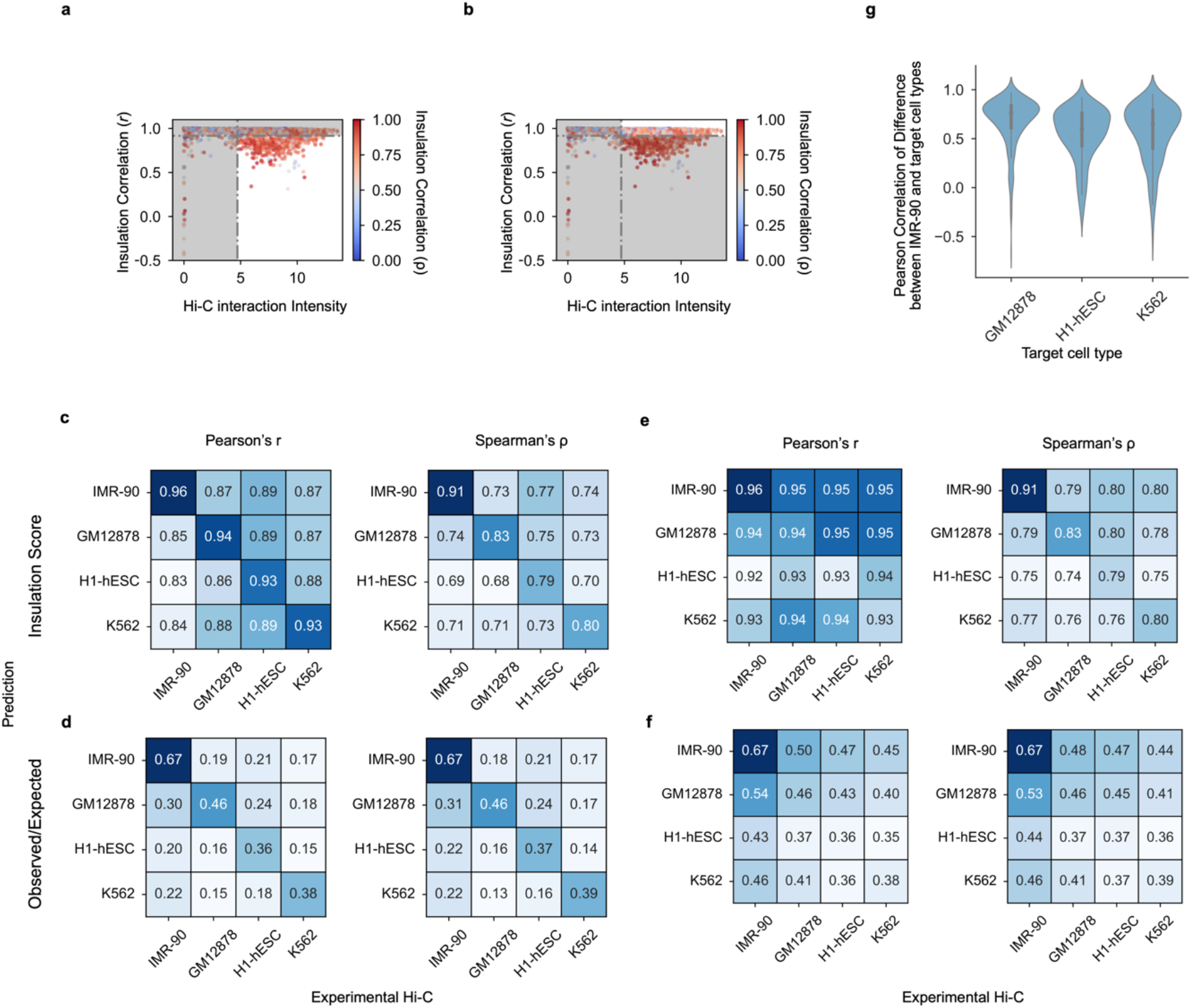
Genome-wide statistics on cell type-specific prediction. **a-b**, The distribution of interaction intensity by insulation correlation (Pearson) between the experimental Hi-C matrices of IMR-90 and GM12878. Dotted lines denote the filtering criteria in selecting representative loci with cell-type specificity (**a**) or structurally conserved regions between two cell types (**b**). Colormap indicates the corresponding Spearman correlation coefficient (*ρ*). **c-d**, Pearson’s *r* (left) and Spearman’s *ρ* (right) between prediction (row) and experimental data (column) for different cell types with insulation score (**c**) and observed/expected score (**d**) as metrics. Diagonal entries denote the metrics of prediction and Hi-C in the same cell type without filtering for cell type specific regions. The scores were calculated based on the differentially structured loci defined in Fig. 3. **e-f**, Same as **c-d** but for the structurally conserved loci across different cell types. **g**, Pearson’s *r* of predicted insulation difference and experimental insulation difference between IMR-90 and other cell types. The correlation was calculated as: Pearson(*Insu*(IMR-90_pred) - *Insu*(Target_pred), *Insu*(IMR-90_data) - *Insu*(Target_data)). High correlation indicates that our model detected cell types-specific features applicable across different cell types.

**Supplementary Figure 17:**
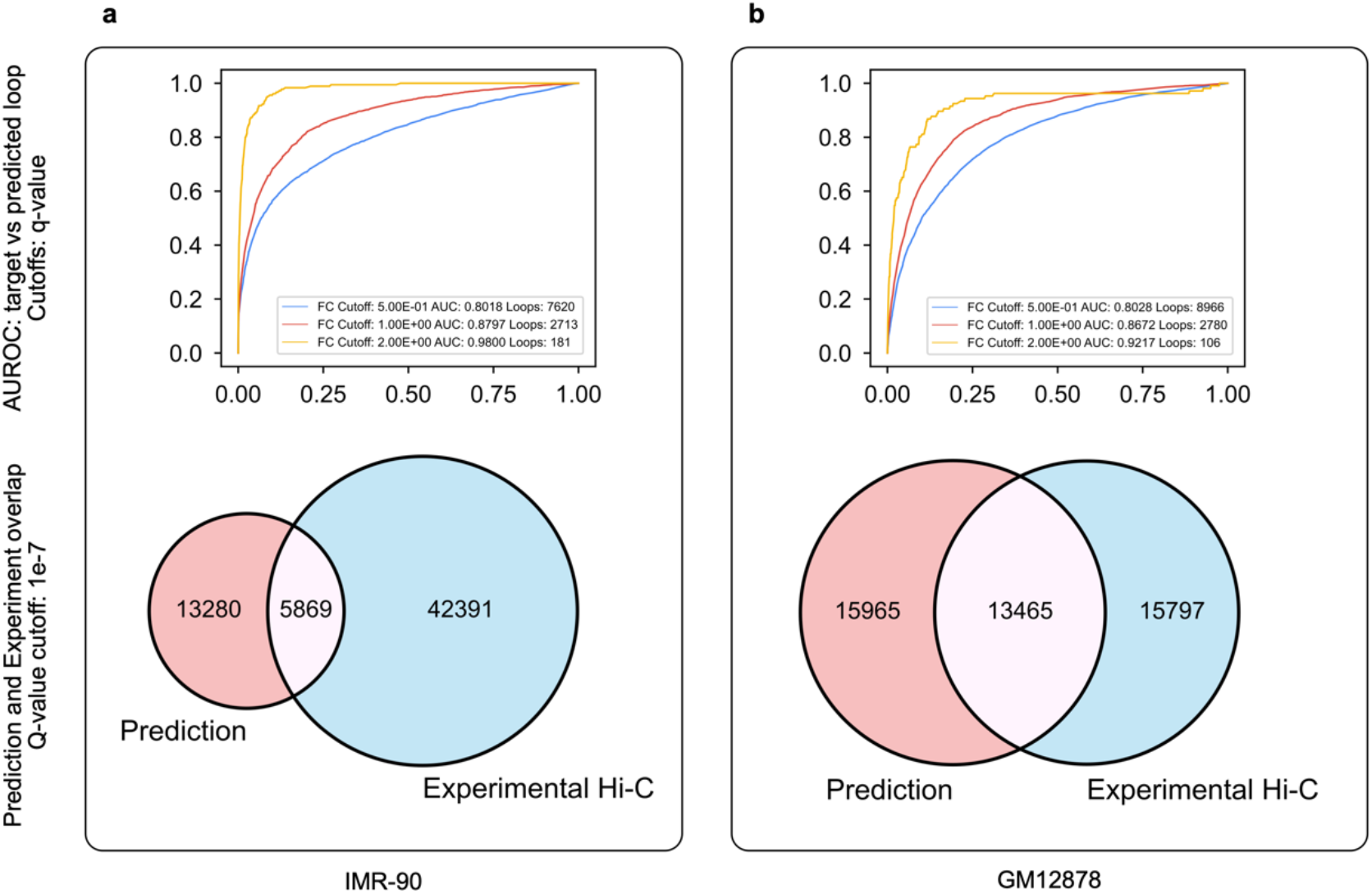
Performance of detecting cell type-specific loop interactions between IMR-90 and GM12878. **a-b**, Comparing cell type-specific loops between prediction and experiment in IMP-90 (**a**) and GM12878 cells (**b**). Loops detected from prediction were first filtered with a more stringent q-value cutoff of 1e-7 in both cell types. We then calculated cell type-specific loops according to signal value fold change. Within each panel, AUROC between loops from experiment and prediction was calculated with log2 fold change cutoffs ranging from 0.5 to 2. Overlap between loops called from prediction and experimental data is presented in a Venn diagram with a q-value cutoff of 1e-7.

**Supplementary Figure 18:**
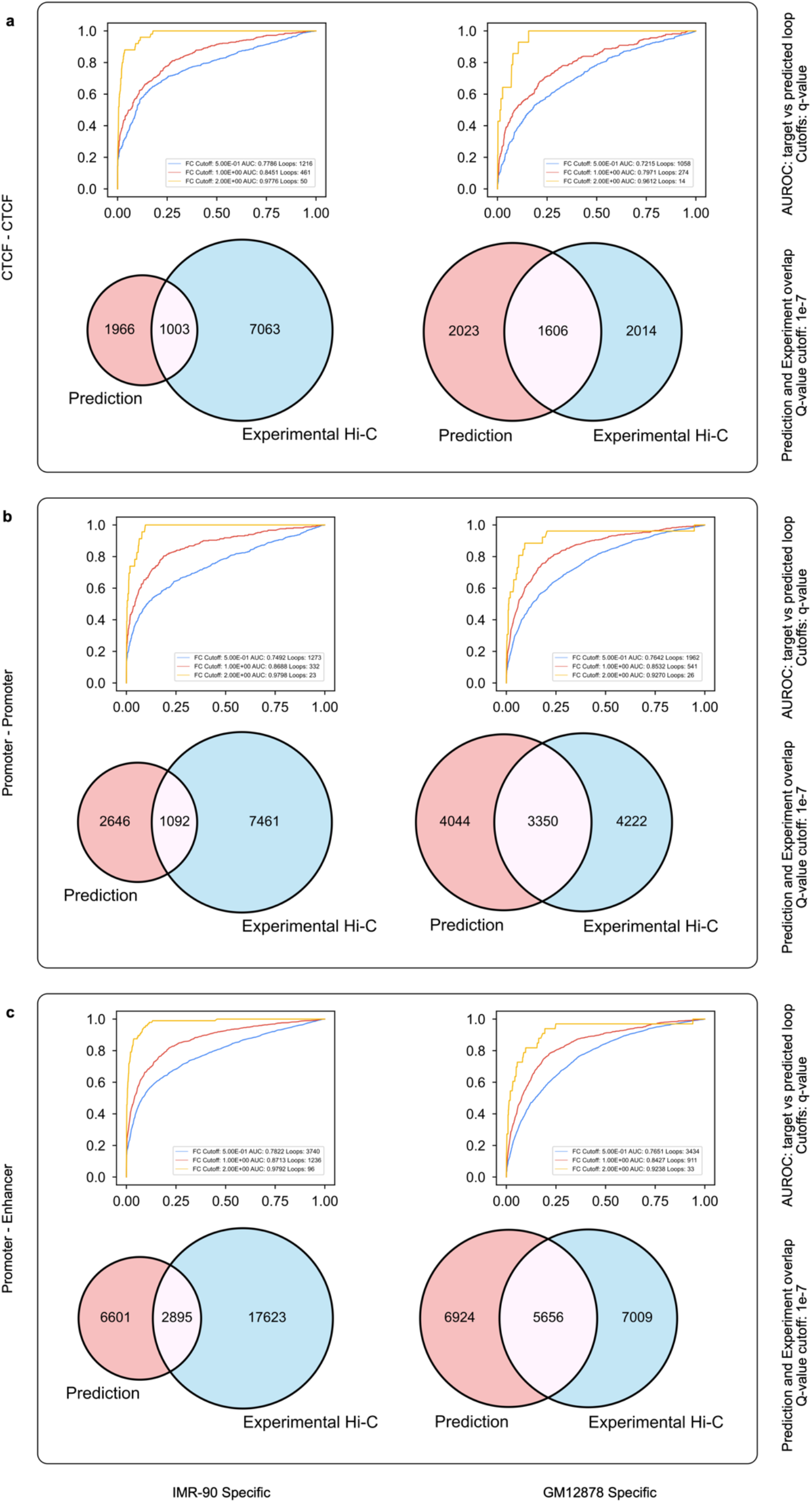
Performance of detecting cell type-specific loop interactions between IMR-90 and GM12878 under different chromatin backgrounds. **a-c**, Evaluating cell type-specific loop detection performance in three types of loops: CTCF-CTCF loop (**a**), promoter-promoter loop (**b**), and promoter-enhancer loop (**c**). Loops were first filtered with a stringent q-value cutoff of 1e-7. We then calculated cell type-specific loops according to signal value fold change. Within each panel, AUROC between loops from experiment and prediction was calculated with log2 fold change cutoffs ranging from 0.5 to 2. Overlap between loops called from prediction and experimental data is presented in a Venn diagram with a q-value cutoff of 1e-7.

**Supplementary Figure 19:**
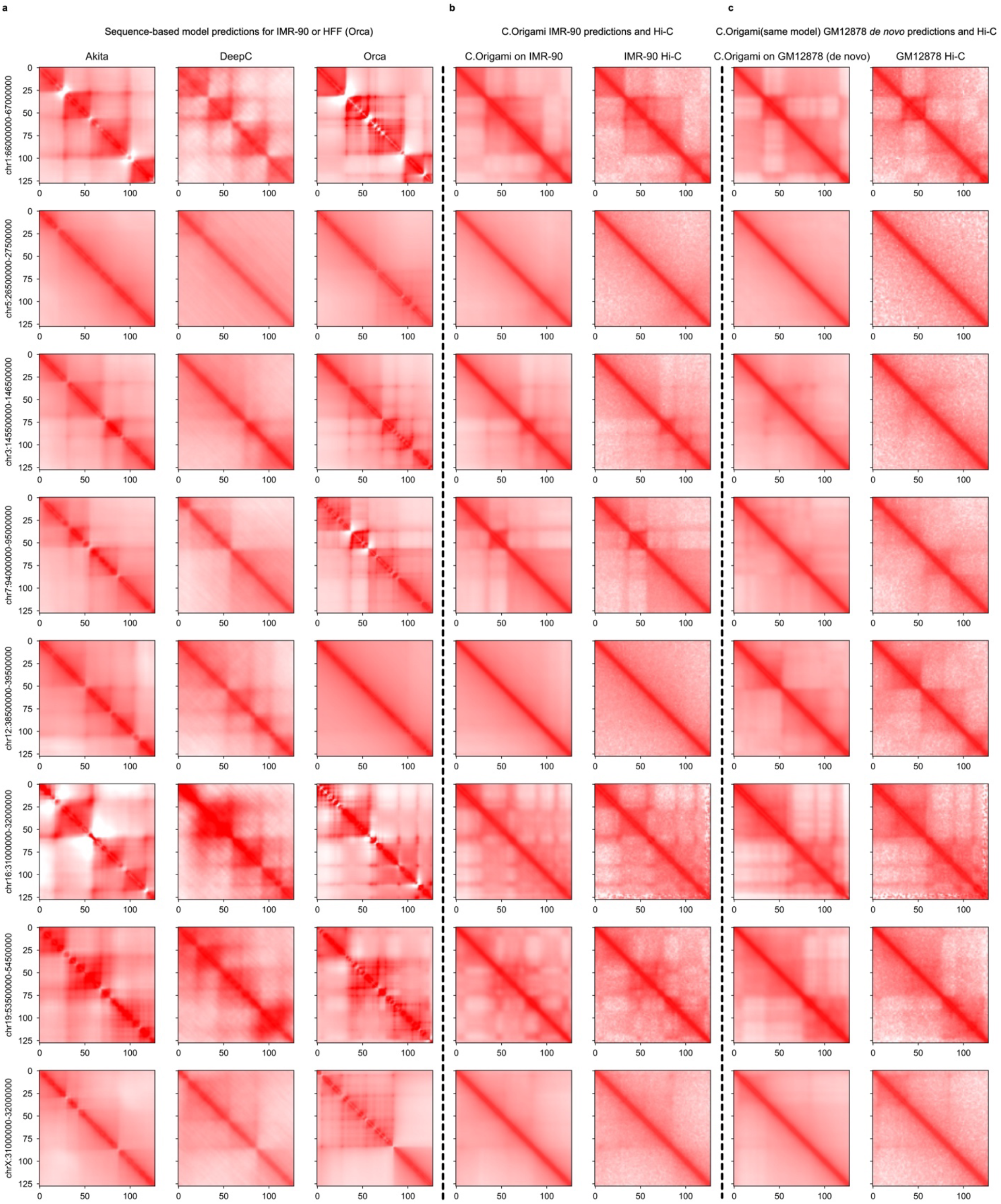
Randomly selected examples of cell type-specific predictions from Akita, DeepC, Orca, and C.Origami. **a**, Sequence-based model predictions, **b**, C.Origami prediction with IMR-90-specific genomic features (CTCF ChIP-seq and ATAC-seq) and IMR-90 experimental Hi-C, **c**, C.Origami *de novo* prediction with GM12878 specific genomic features and GM12878 experimental Hi-C. All presented results were aligned at randomly selected regions from different chromosomes. The full set of prediction results across all cell type-specific chromatin regions between IMR-90 and GM12878 cells were included in the Supplementary material under Cell type-specific predictions.

**Supplementary Figure 20:**
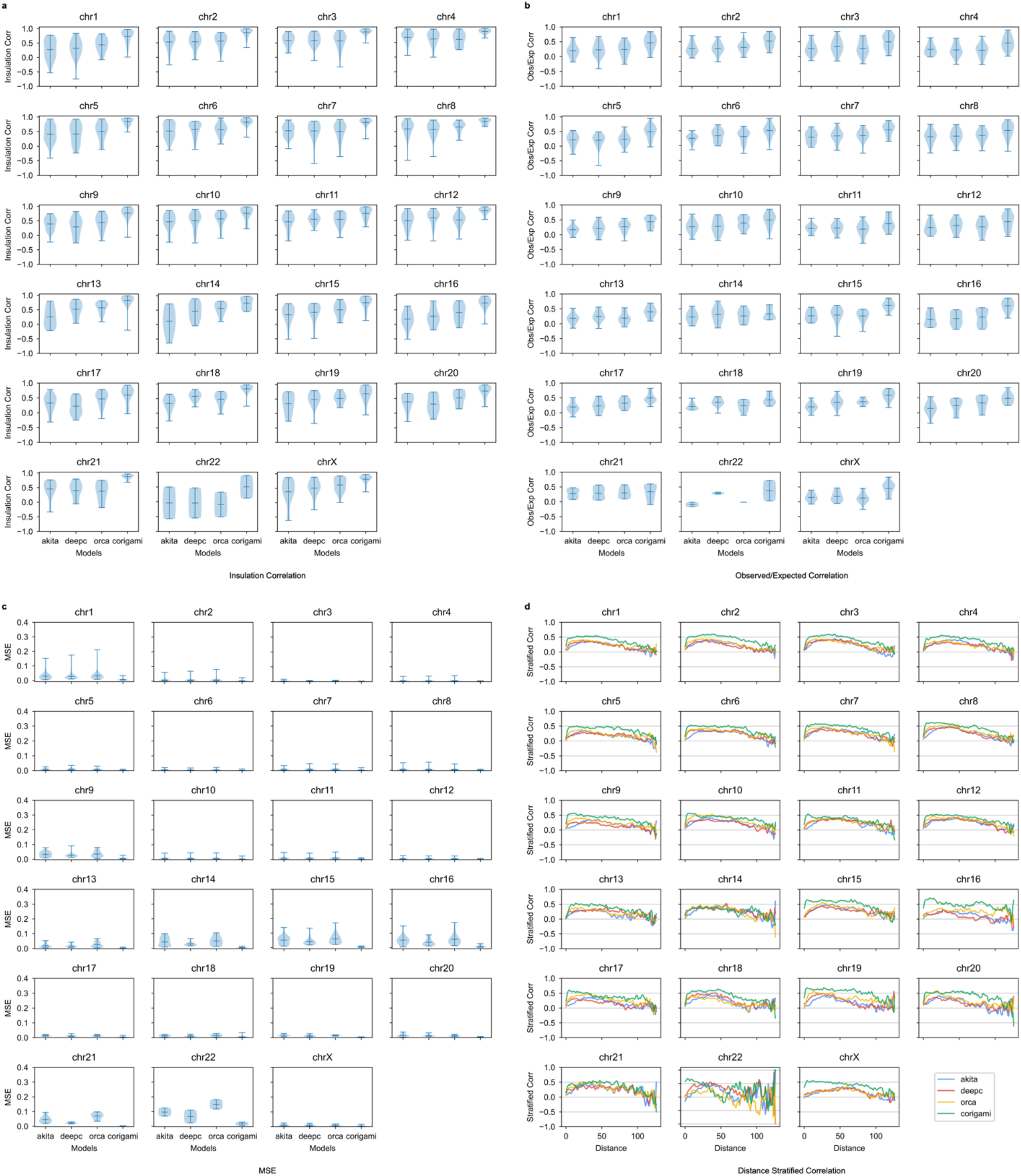
Genome-wide comparison of *de novo* prediction quality in GM12878. For *de novo* prediction results from each model (Akita, DeepC, Orca and C.Origami), we measured insulation score correlation (**a**), observed vs expected Hi-C matrices correlation (**b**), mean squared error (MSE, **c**), and distance-stratified correlation (**d**). Prediction results at cell type-specific regions between IMR-90 and GM12878 cells were selected for this analysis.

**Supplementary Figure 21:**
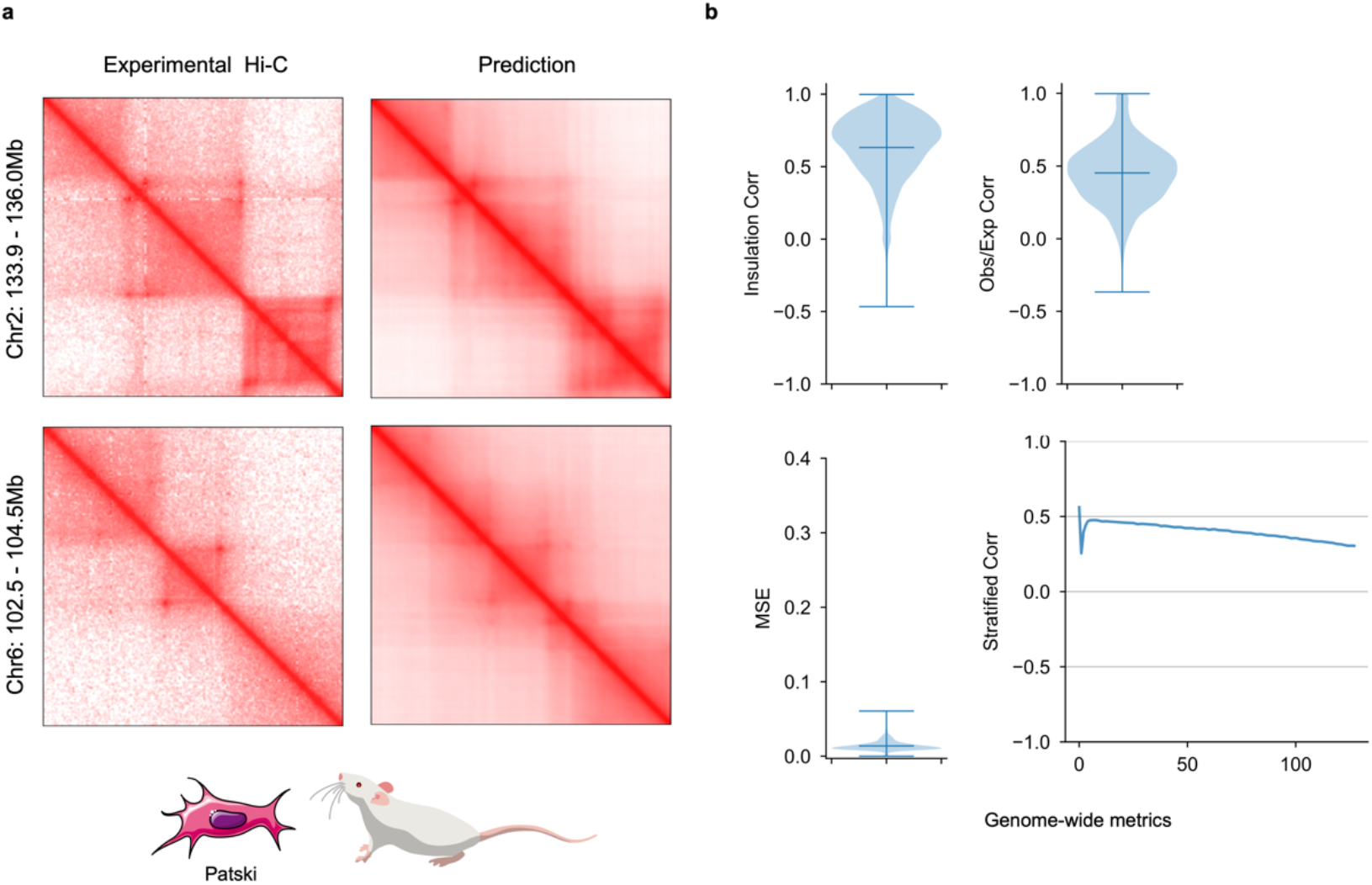
Transferring model trained on human cell type to mouse. **a,** Experimental Hi-C and C.Origami prediction results of two representative loci in hybrid mouse Patski cells. **b**, Genome-wide performance metrics of predicting mouse chromatin organization using C.Origami trained with human data. Presented matrices include insulation score correlation, observed vs expected matrix correlation, mean squared error, and distance-stratified correlation. Error bars in the violin plots indicate minimum, mean and maximum values.

**Supplementary Figure 22:**
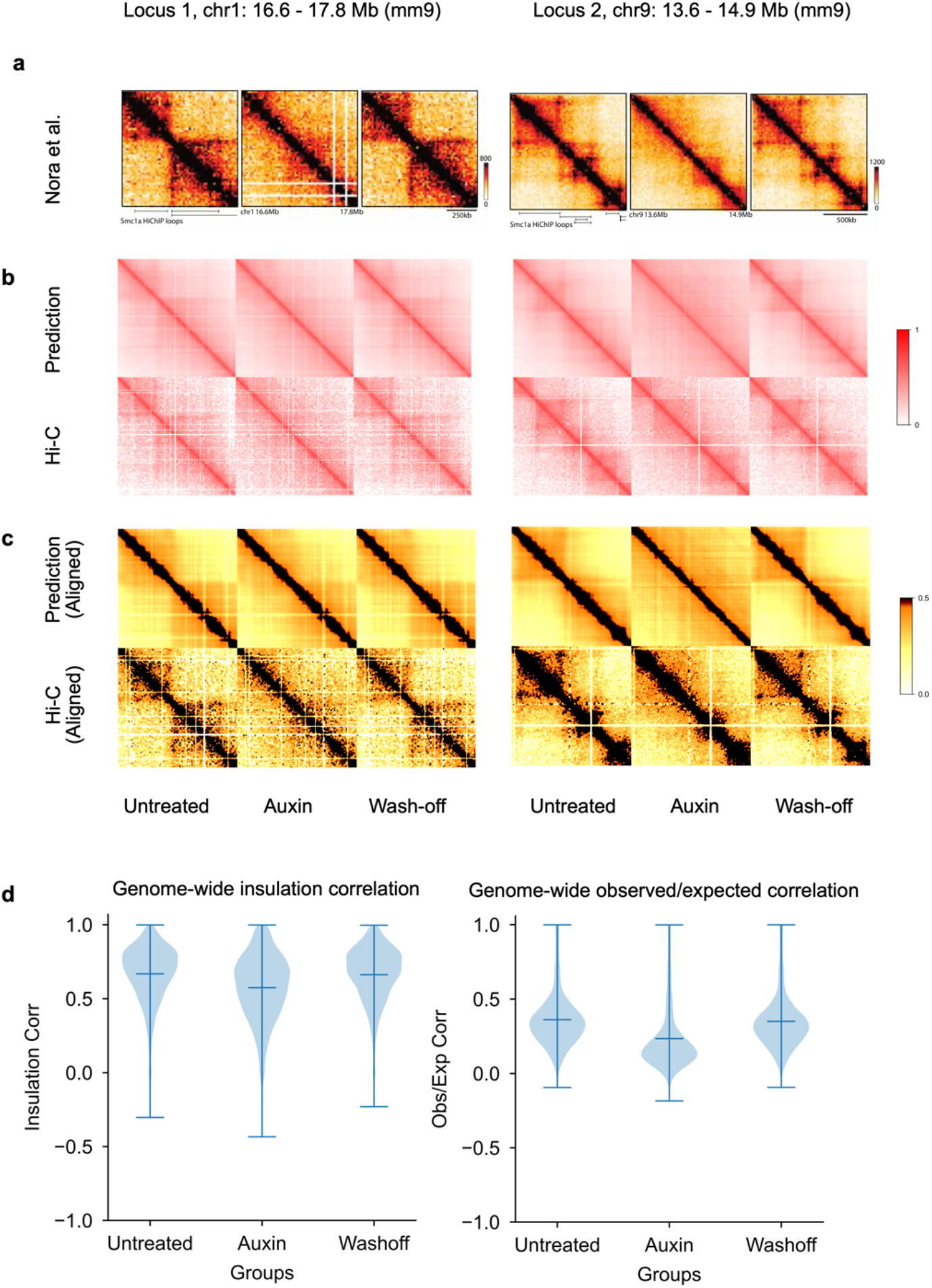
Predicting chromatin organization dynamics upon auxin-induced CTCF depletion and restoration in mESCs. **a**, Experimental results adopted from Nora *et al*^38^. at two loci indicated on top. All plots were visualized in triplicates, indicating conditions of before CTCF depletion (Untreated), CTCF depleted (Auxin), and CTCF restored (Wash-off). **b**, C.Origami prediction at the corresponding 2Mb-wide windows using DNA sequence and CTCF ChIP-seq profiles from Nora *et al.* Corresponding experimental Hi-C matrices from Nora *et al*. were processed by HiC-bench and visualized in parallel. **c**, Adjusted prediction and Hi-C matrices from **b**. Matrix size and location were adjusted to match the exact position from the experimental results as shown in **a**. Colormap was adjusted to match the original figure in Nora *et al*. **d**, Genome-wide performance metrics for evaluating C.Origami prediction upon CTCF depletion and restoration. Presented correlations include insulation score (left panel) and observed vs expected matrix values (right panel). Error bars in the violin plots indicate minimum, mean and maximum values.

**Supplementary Figure 23:**
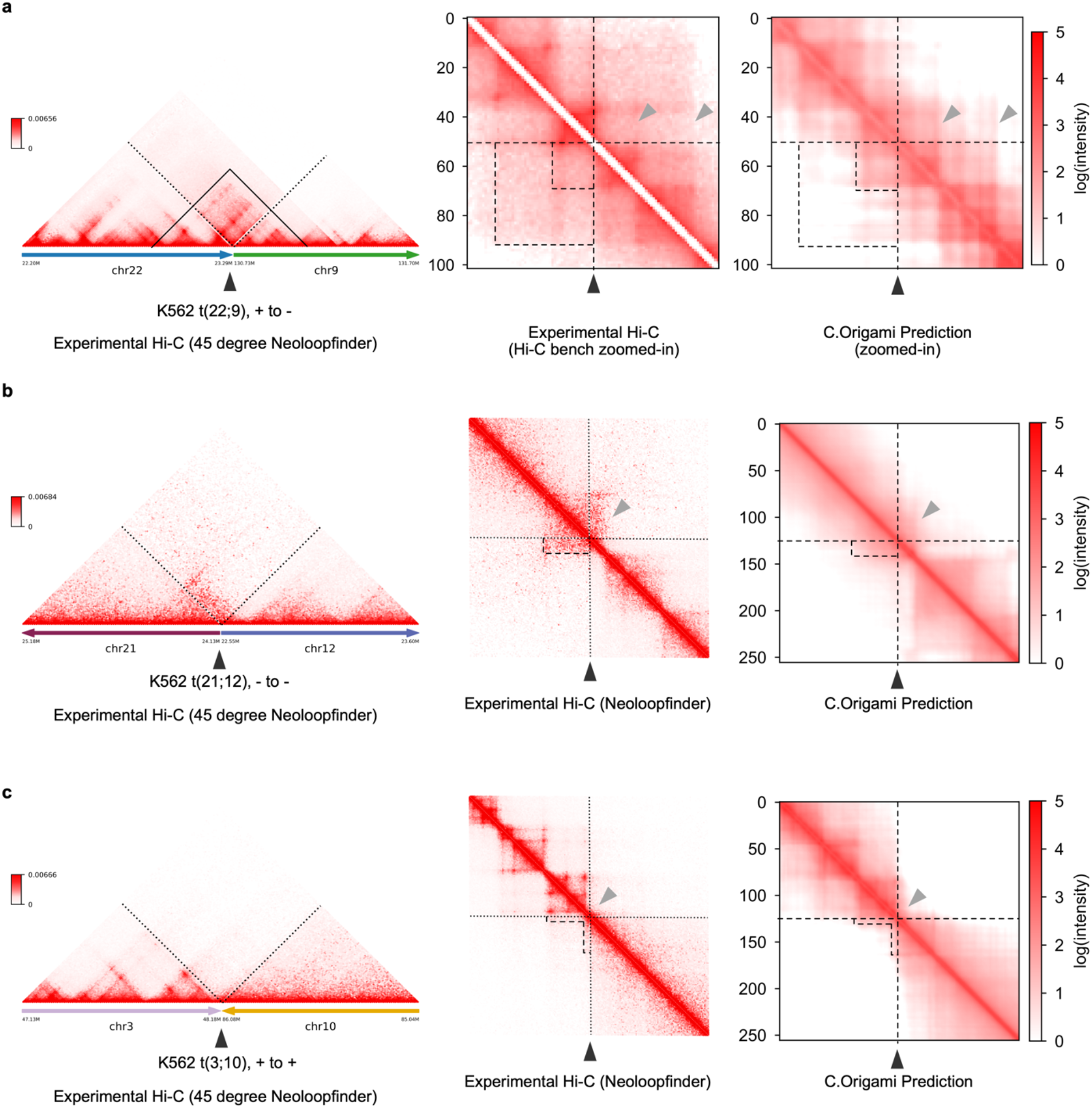
Predicting translocation-induced novel chromatin organizations in K562 cells. **a**-**c,** Experimental and predicted Hi-C matrices at three translocation loci in K562 cells. In each case, chromatin organization structures were first reconstructed using HiC-bench^64^ and NeoLoopFinder^41^, followed by C.Origami prediction at the translocation loci using *in silico* fused genomic information. **a**, t(22;9) translocation, also known as the Philadelphia chromosome, that leads to a fused gene *BCR-ABL1*. **b**, t(21;12) translocation with a stripe interaction. **c**, t(3;10) translocation with a faint “L”-shape interactions as indicated by the dotted contour. Dotted boxes indicate neo-TAD forming at the translocation site. Black arrowhead indicates the translocation site.

**Supplementary Figure 24:**
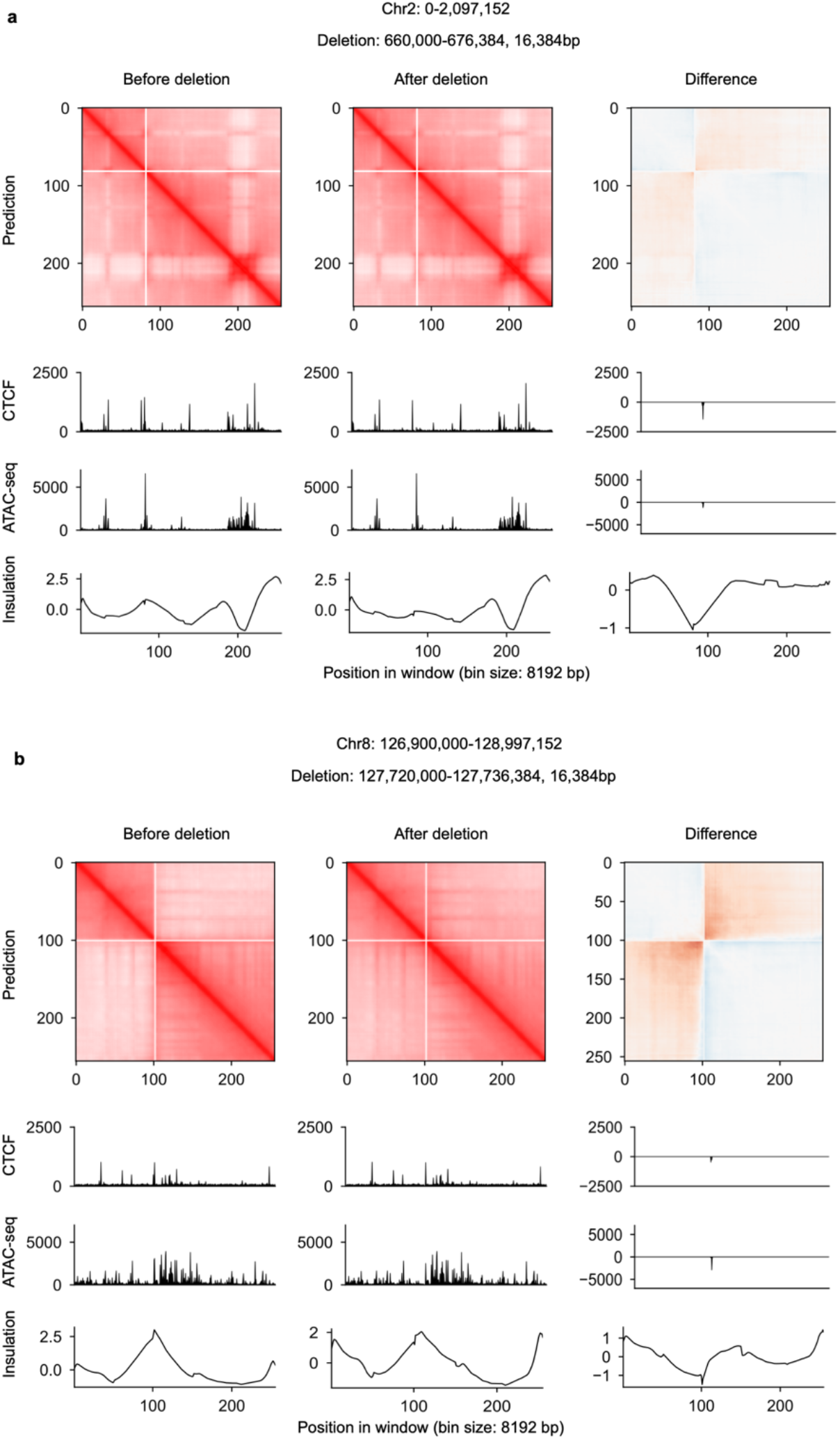
*In silico* genetic experiments performed on IMR-90 cells. Two *in silico* deletion experiments were separately represented in **a** and **b**. Each experiment included the prediction before (left) and after deletion (middle). The difference in chromatin folding after deletion were presented on the right.

**Supplementary Figure 25:**
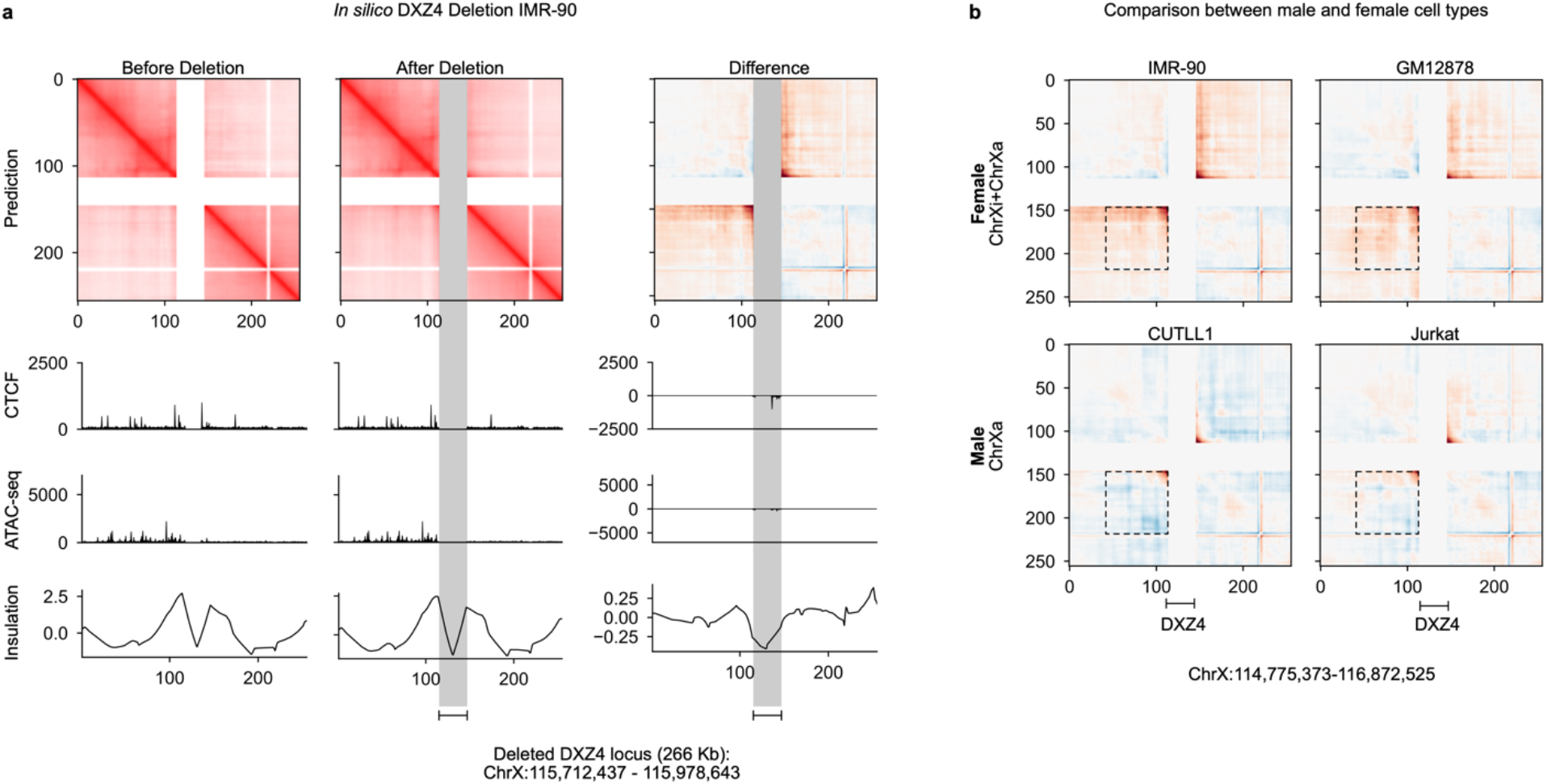
Predicting X chromosome organization changes upon *DXZ4* deletion prediction in male and female cell types. **a**, Chromatin organization changes upon *in silico* deletion of a 266Kb repeats at the *DXZ4* locus in IMR-90, a female cell line. The perturbed region mimics the experimental knock-out in Darrow et al^45^. The deleted region is indicated by a gray bar. **b**, Chromatin organization changes upon *in silico* deletion of the *DXZ4* locus in two female cell lines (IMR-90, GM12878), and two the male cell lines (bottom: CUTLL1, Jurkat). Deleting *DXZ4* locus led to substantial loss of insulation at the two flanking regions of *DXZ4* locus in the female cell lines, while the effect was very minimal in the male cell lines, supporting the role of *DXZ4* in regulation X chromosome inactivation. Interaction regions are denoted by dotted boxes.

**Supplementary Figure 26:**
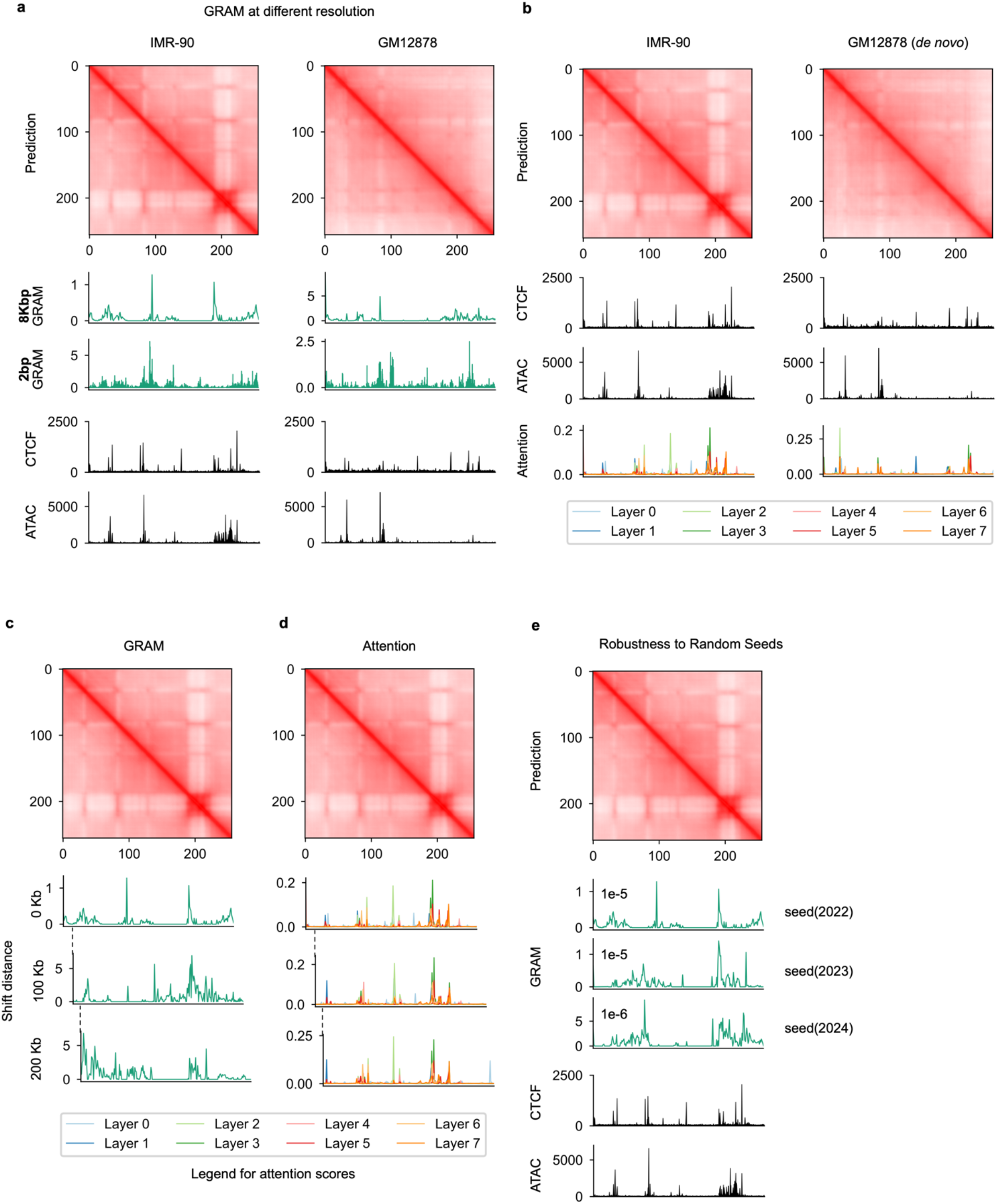
Comparing GRAM and attention scores. **a**, Comparison of GRAM scores at 2bp and 8kb resolution in IMR-90 (left) and GM12878 (right). **b**, Attention scores on IMR-90 and GM12878. Attention scores on different layers were colored according to legends. **c-d**, Comparison between GRAM (**c**) and attention scores (**d**) at three consecutive windows with 100Kb shifts. **e**, GRAM scores generated at different PyTorch random seeds.

**Supplementary Figure 27:**
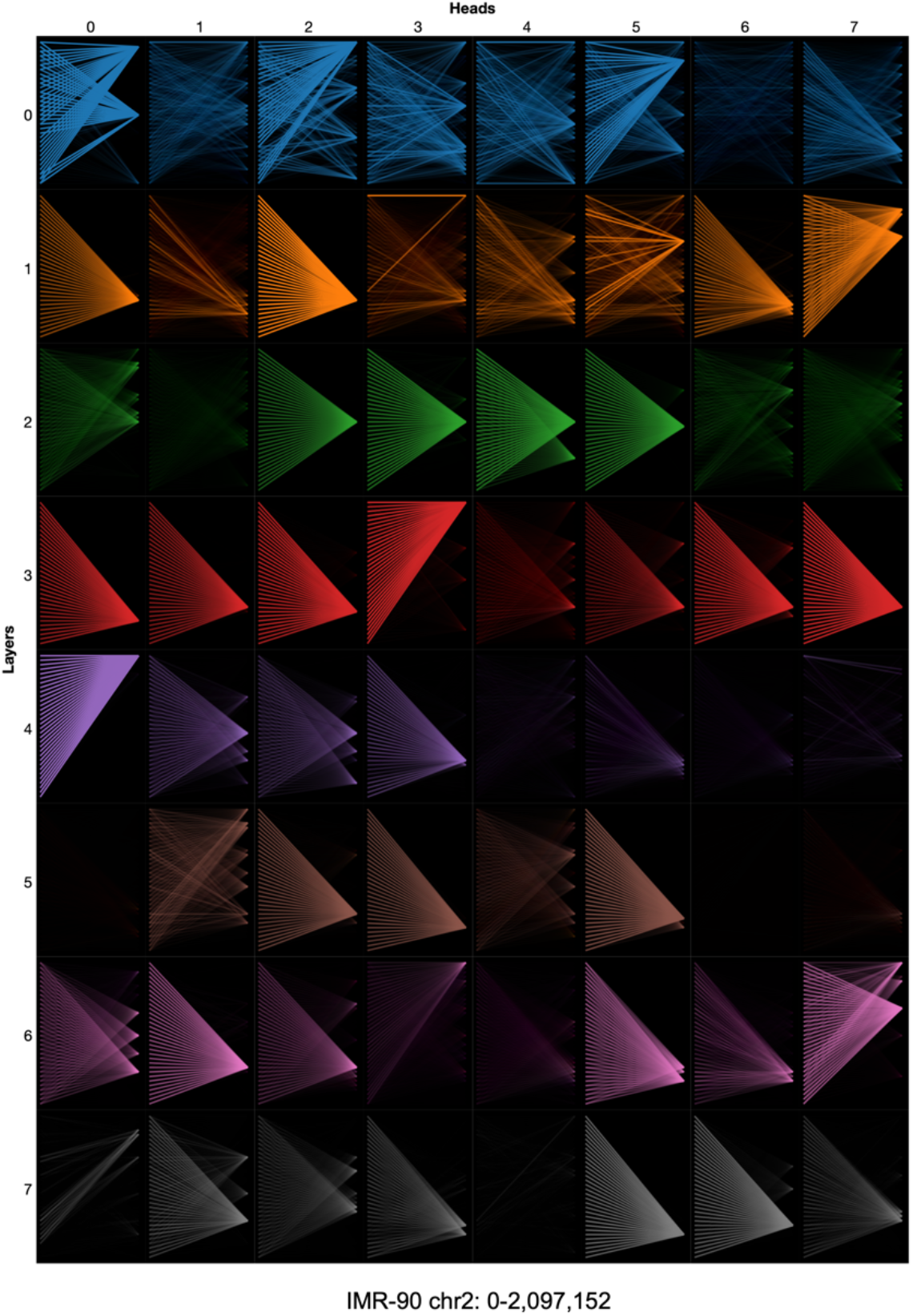
Attention weights generated from the transformer module of C.Origami. A detailed view of the attention weights in eight heads (columns) across eight layers (rows), generated by the BertViz package^73^. The y axis of each row represents a 2Mb genomic distance. Brightness of the line segment between two different locations denotes interaction intensity.

**Supplementary Figure 28:**
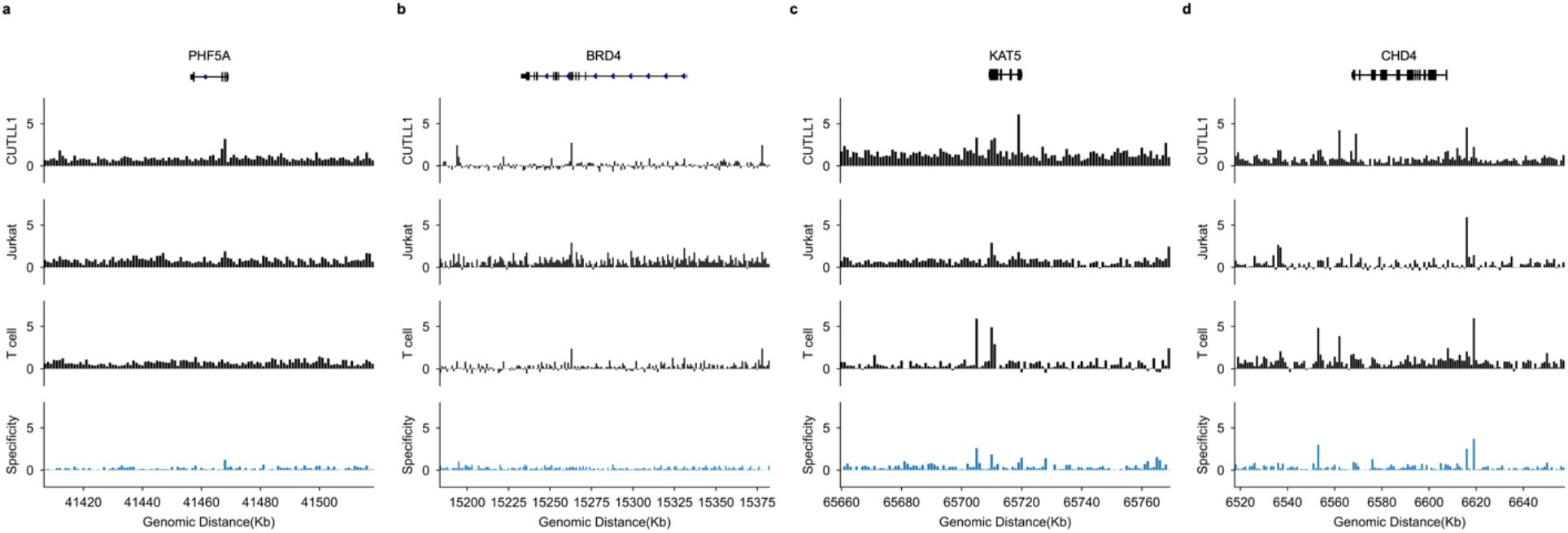
ISGS-identified impact scores at four chromatin remodeler genes in both T-ALL cells and T cells. Impact scores of the DNA elements in T-ALL cells and normal T cells were first calculated independently through ISGS, and then visualized at the four chromatin remodelers genes (*PHF5A*, *BRD4*, *KAT5*, *CHD4,* with 50Kb upstream and 50Kb downstream) which are required for Jurkat and CUTLL1 cell proliferation according to the CRISPR screening experiments. The specificity track (fourth track) was calculated as the difference between T cell impact score and T-ALL impact score (from CUTLL1 or Jurkat, whichever is smaller). *CHD4* has the highest specificity score between T-ALL cells and normal T cells.

**Supplementary Figure 29:**
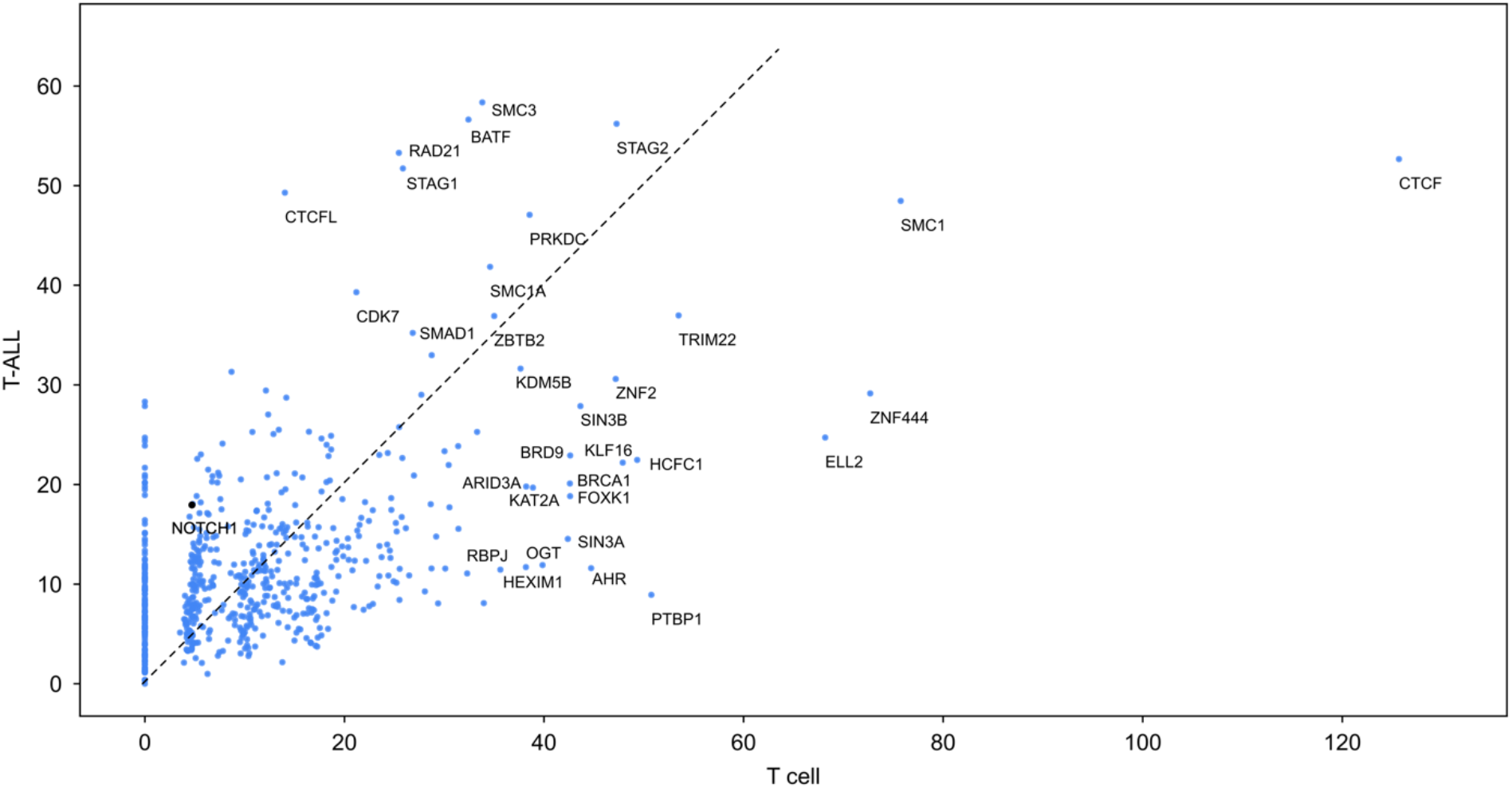
Scatter plot of *trans*-acting factor binding enrichment in ISGS-identified impactful elements in T-ALL and normal T cells. Odds ratio of enrichment between T-ALL and normal T cells were ploted on the y axis and x axis, respectively. T-ALL odds ratio was aggregated from enrichment in CUTLL1 and Jurkat. Only factors with odds ratio larger than 35 were labeled, except NOTCH1 which was highlighted for comparison with CDK7 (referring to Figure 7).

**Supplementary Figure 30:**
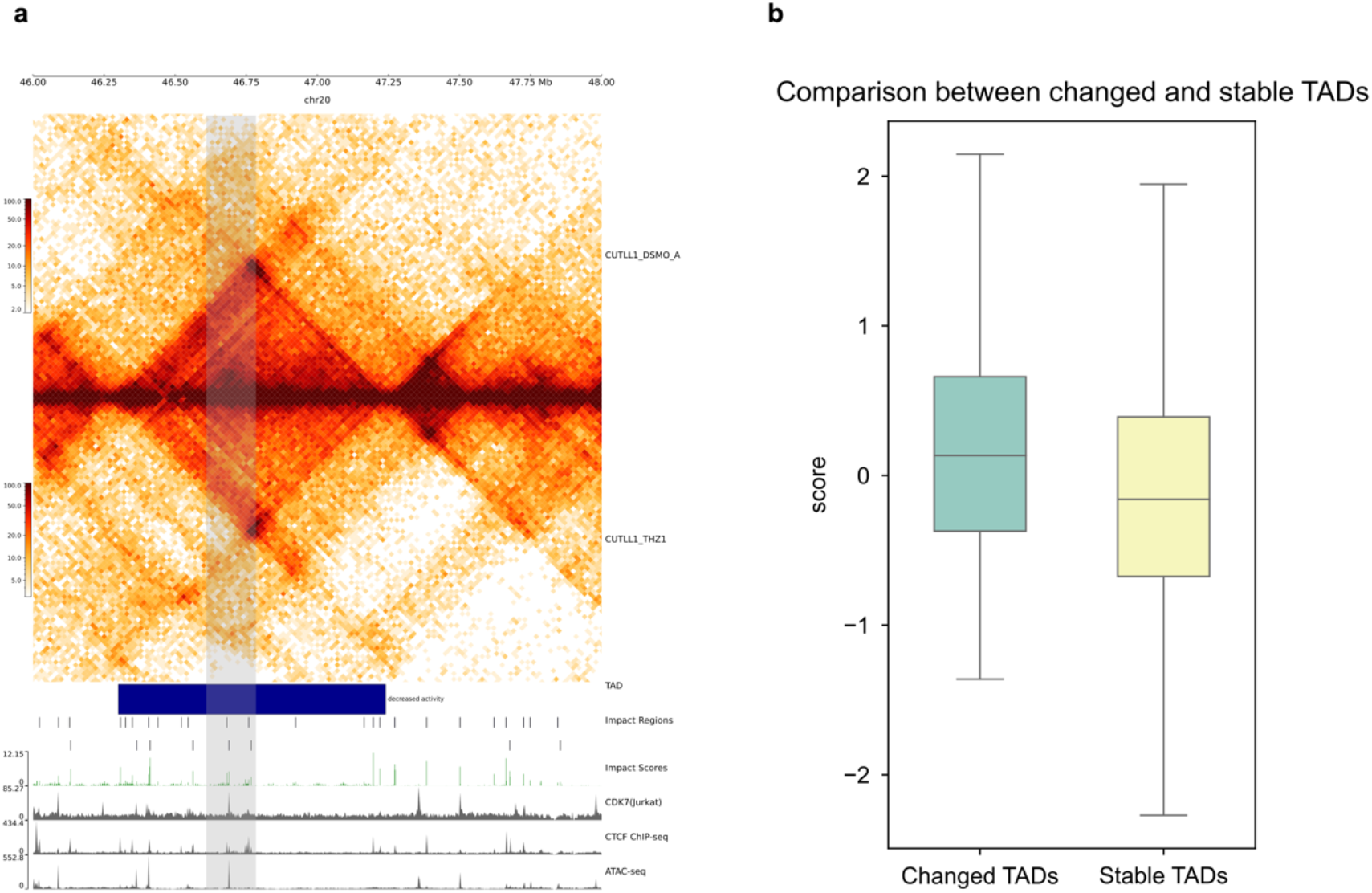
Overlap between impactful elements and CDK7-inhibition induced TAD changes. **a**, An example of TAD with decreased activity. Grey bar indicates a prominent decrease in interaction in the CDK7-inhibition (+THZ1) group. The TAD intensity plots were aligned with impactful regions, impactful scores, CDK7 ChIP-seq, CTCF ChIP-seq, and ATAC-seq signals from top to bottom. **b**, Impact score of DNA elements in changed TADs and stable TADs determined from pharmaceutical inhibition of CDK7. The overall impact scores in the changed TADs are significantly higher (independent t-test, p-value = 1.72e-05).

